# An immunomechanical checkpoint PYK2 governs monocyte-to-macrophage differentiation in pancreatic cancer

**DOI:** 10.1101/2024.11.19.624405

**Authors:** Wenyan Xie, Xin Yu, Qingxin Yang, Nengwen Ke, Ping Wang, Hao Kong, Xiangji Wu, Panpan Ma, Lang Chen, Jie Yang, Xiuqing Feng, Yuan Wang, Hubing Shi, Lu Chen, Yun-Hua Liu, Bi-Sen Ding, Qiang Wei, Hong Jiang

## Abstract

Pancreatic ductal adenocarcinoma (PDAC) is characterized by a fibrotic, stiff tumor microenvironment (TME), where tumor-associated macrophages (TAMs) drive ECM remodeling, progression, and immune evasion. The contribution of mechanical cues to monocyte differentiation into TAMs remains largely unexplored. Here we show that mechanical force is required for monocyte-to-macrophage differentiation. PYK2, as an innovative immunomechanical checkpoint, de facto governs this differentiation process. We demonstrated that PYK2 senses mechanical signals via Piezo1 and integrins, triggering F-actin polymerization and translocating to the nucleus to regulate mechanotransduction and differentiation genes (e.g., *ACTR3*, *RELA*). Targeted deletion of PYK2 impairs the differentiation and polarization of monocyte-derived macrophages, reshapes the PDAC microenvironment, and enhances the efficacy of anti-PD-1 immunotherapy. These findings underscore the critical role of mechanical cues in monocyte differentiation and suggest that targeting PYK2 is a promising strategy to modulate TAM function and improve immunotherapy outcomes in patients with PDAC.

**Statement of significance:** This study identifies PYK2 as an immunomechanical checkpoint that drives monocyte-to-macrophage differentiation in PDAC via Piezo1/integrin-mediated mechanical cues. Targeted deletion of PYK2 reshapes the PDAC microenvironment, and enhances the efficacy of anti-PD-1 immunotherapy, suggesting PYK2 as a promising therapeutic target to overcome immunotherapy resistance.

## INTRODUCTION

The physical features of cancer significantly contribute to its progression and affect its response to therapy (1). One of the most notable mechanical abnormalities in tumors is the increased tissue stiffness (1,2), which is primarily due to the fibrotic nature of tumors characterized by the increased deposition and crosslinking of extracellular matrix (ECM) proteins (3). The rigid tumor microenvironment (TME), shaped by cancer, fosters an immune-suppressive milieu and impedes the efficacy of immunotherapy (2,4). Targeting ECM stiffness has been shown to improve the efficacy of immunotherapy (5–7).

Pancreatic ductal adenocarcinoma (PDAC) is an aggressive and lethal cancer characterized by a distinct, dense and desmoplastic stroma (8,9), reflecting its highly stiff TME. Tumor-associated macrophages (TAMs) are among the most predominant immune cells within the TME and promote ECM remodeling, tumor growth and immune suppression (10,11). Macrophages can originate from either embryonic precursors or adult circulating monocytes with tissue specificity (12). In a murine model of PDAC, it was found that approximately 90% of TAMs were derived from monocytes (13). While it is well established that biochemical signals can trigger the monocyte differentiation into TAMs (14,15), the role of mechanical stimuli and training within the PDAC TME in this process remains unclear. Although increased stiffness in the TME has been linked to driving macrophages toward an alternatively activated state and enhancing their immunosuppressive functions, the precise mechanism behind this process has yet to be fully elucidated (4,16,17). Exploring the impact of mechanical cues on the differentiation and polarization of monocytes/macrophages could provide valuable insights and potential strategies for targeting TAMs.

Mechanosensitive ion channels and cell adhesion molecules, including integrins, selectins, and cadherins, play critical roles in sensing and translating mechanical signals into biochemical responses across various cell types (18–20). Recent research has revealed that macrophages can detect substrate stiffness and respond by activating the mechanosensitive transcription factor yes-associated protein 1 (YAP1) (21). Mechanosensitive ion channels, such as PIEZO channels and TRPV4, have been identified for their involvement in the sensing of stiffness, polarization, and function in macrophages, potentially complementing the adhesive role of integrins (22–26). Circulating monocytes can sense fluid shear stress via Piezo1, and high shear stress promotes their proinflammatory phenotype (27). The viscoelasticity of myelofibrosis contributes to the aberrant differentiation of monocytes into dendritic cells (28). However, whether and how monocytes sense and respond to the biophysical characteristics of the PDAC TME during the differentiation process into macrophages remains largely unexplored.

In this study, we demonstrated that mechanical training is essential for the differentiation of monocytes into macrophages. PYK2, a member of the focal adhesion kinase (FAK) family, is an innovative immunomechanical checkpoint in monocytes that is responsible for their mechanosensing, mechanotransduction and differentiation processes. In addition to its role in generating cellular force, PYK2 also functions as a potential transcription factor that assists in the transcription of genes associated with mechanotransduction and differentiation. Inhibiting PYK2 disrupts the maturation and polarization of monocyte-derived macrophages, modifies the physical properties of the TME in PDAC, and significantly enhances the response to immunotherapy.

## RESULTS

### ECM-associated mechanical stimuli promote differentiation of myeloid precursors into macrophages in PDAC

To investigate the impact of high fibrosis-induced mechanical stimuli on cell activities in PDAC, we generated comprehensive data using nanoindentation (419 measurements from 7 PDAC and 3 adjacent normal samples), sequential immunohistochemistry (IHC) (3 PDAC and 3 adjacent normal samples with 9 markers), and spatial transcriptomics (28,668 spots from 4 PDAC and 2 adjacent normal samples) of human PDAC samples (Fig. 1a; Supplementary Fig. S1a-b; Supplementary Fig. S2a-c; Supplementary Table 1). Additionally, we incorporated an external PDAC atlas single-cell RNA sequencing (scRNA-seq) dataset, comprising 26 PDAC and 19 normal samples, resulting in a total of 167,818 single-cell transcriptomes for detailed cellular analysis (Fig. 1a; Supplementary Fig. S2d-f).

**Fig. 1.**
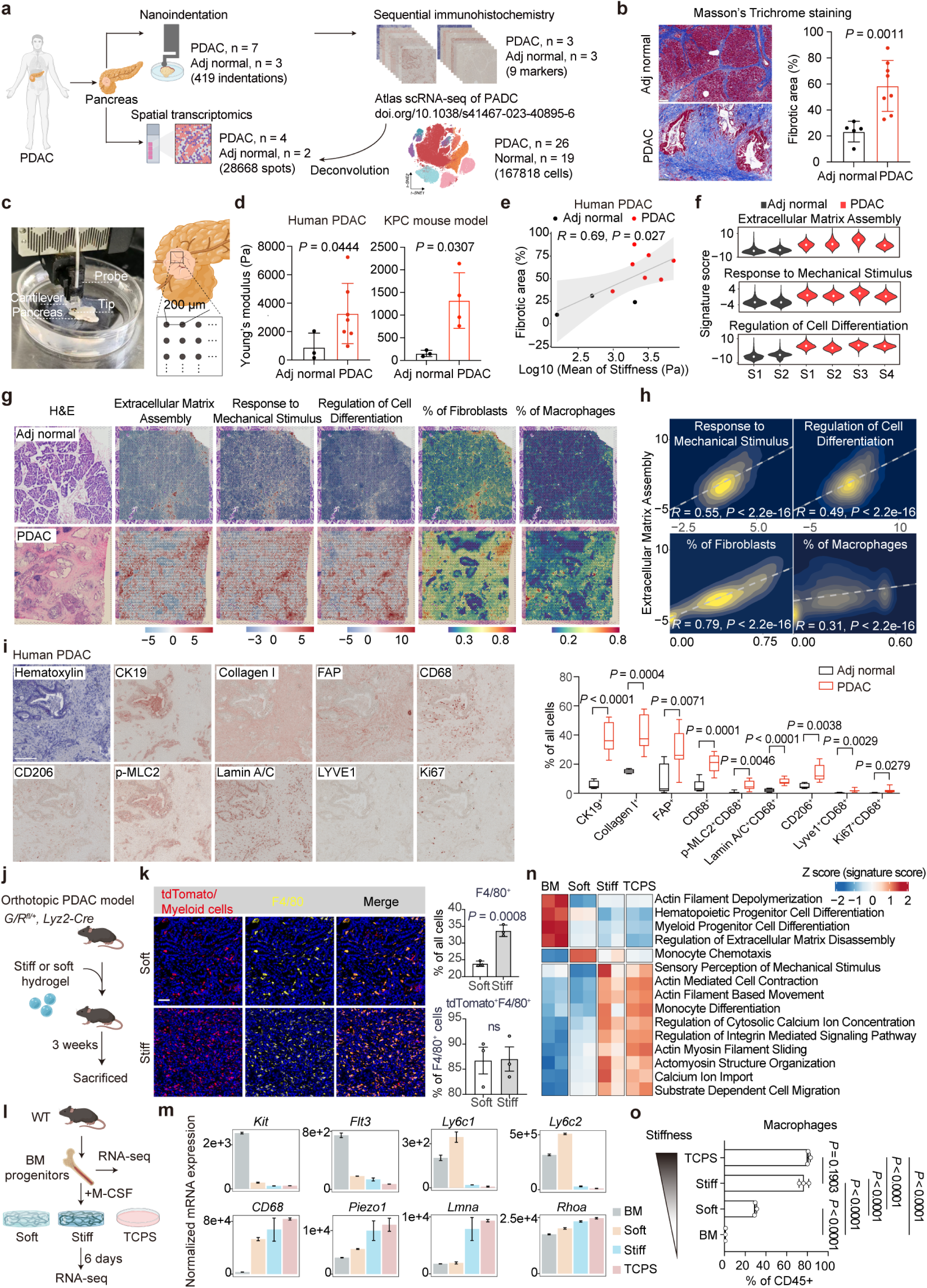
Stiff PDAC TME promotes monocyte differentiation into macrophage. **a**, Experimental workflow of human PDAC samples for nanoindentation, Masson’s trichrome staining, sequential immunohistochemistry (IHC) staining, and spatial transcriptomics (ST) analysis. Fresh adjacent normal and tumor tissues from human PDAC patients were procured and divided into two portions: one portion underwent immediate processing for nanoindentation analysis, followed by formalin-fixation, paraffin-embedding, and subsequent sectioning for Masson’s trichrome staining, and mIHC staining. The other portion was embedded in OCT, frozen and sectioned for ST analysis. Previous scRNA-seq data from human normal pancreas and PDAC was used for ST cell type deconvolution(29). **b**, Representative images of Masson’s trichrome staining (left) and quantification (right) of the fibrotic area in human adjacent normal (n = 5) and PDAC (n = 8) samples using HALO software. Scale bars, 100 μm. **c**, Measurement of Young’s modulus of pancreatic tissues using nanoindentation. The image (left) captures the nanoindenter probe and tissue specimens. Each sample was indented with 200 μm intervals between measurements (right). **d**, The Young’s modulus of adjacent normal (n = 3) and PDAC (n = 7) tissues in human PDAC patients, as well as in the KPC mouse model (3 adjacent normal samples and 4 PDAC samples), was measured via nanoindentation testing. **e**, The scatter plot showing the relationship between the fibrotic area (%) and the log-transformed mean stiffness (Pa) in human adjacent normal and PDAC tissues. Black dots represent adjacent normal tissues (n = 3), and red dots represent PDAC tissues (n = 7). The grey line represents the linear fit, and the shaded grey area indicates the 95% confidence interval of the fit. The Pearson correlation coefficient (*R*) is 0.69 with a *P* value of 0.027. **f-h**, Spatial transcriptomics of human adjacent normal and PDAC samples. **f**, Comparison of the signature scores for three specific pathways, e.g., extracellular matrix assembly, response to mechanical stimulus, and regulation of cell differentiation in spatial transcriptomics data of human adjacent normal and PDAC samples. **g**, H&E staining and spatial distribution of signature scores associated with the indicated pathways from Fig. 1f, along with the distribution of fibroblasts and macrophages in spatial transcriptomics data for the indicated tissue sections of human adjacent normal and PDAC. Scale bars, 1 mm. **h**, Spearman correlation analysis illustrating the interrelation between the signature scores of extracellular matrix assembly and other pathways, as well as the distribution of cell content, in the spatial transcriptomics data of tumor samples. The correlation coefficients (*R*) and *P* values are shown in the plots. The colors in the plot indicate the density of cell distribution. **i**, Representative images (left) and quantification (right) of sequential IHC staining of CK19, Collagen I, FAP, CD68, CD206, p-MLC2, Lamin A/C, LYVE1, and Ki67 in human PDAC tissue. n = 3 patients in each group. Scale bars, 200 μm. **j**, Schematic of establishment of orthotopic PDAC model. Stiff or soft hydrogel was ground, mixed with KPC cells and orthotopically injected into the pancreas of *G/R^fl/+^, Lyz2-Cre* mice. The mice were sacrificed 3 weeks after implantation. **k,** Representative images (left) and quantification (right) of mIHC staining of F4/80 in orthotopic PDAC model generated in *G/R^fl/+^, Lyz2-Cre* mice. n = 3 mice per group. Scale bars, 50 μm. **l**, Schematic of bulk-RNA sequencing of mouse bone marrow (BM) progenitors and macrophages differentiated on surfaces with varying stiffness. **m**, Normalized gene expression of *Flt3*, *Kit*, *Ly6c1*, *Ly6c2*, *CD68*, *Piezo1*, *Lmna* and *Rhoa* in mouse BM progenitors, macrophages differentiated on soft, stiff, and TCPS surfaces for 6 days, as determined from bulk RNA-seq data. **n**, Heatmap of GSVA analysis showing pathway enrichment in mouse BM progenitors, macrophages differentiated on soft, stiff, and TCPS surfaces for 6 days, based on the bulk RNA-seq data. The colors in the heatmap represent the z-scores of the GSVA-derived pathway enrichment scores. **o**, Quantification of flow cytometry for macrophages in mouse BM, and those differentiated on soft, stiff, and TCPS surfaces over a 6-day period cultivation. *P* values in (**b**), (**d**) and (**k**) were determined by two-tailed unpaired Student’s t-test; *P* values in (**i**), and (**o**) were determined by ordinary one-way ANOVA. Data shown in (**m**) and (**n**) are representative of two experiments. Data shown (**o**) are representative of three experiments.

One prominent characteristic of human PDAC is its extensive fibrotic stroma. Masson’s trichrome staining analysis revealed a substantial fibrotic area present in human PDAC samples (∼60%) compared to adjacent normal tissues (∼20%) (Fig. 1b; Supplementary Fig. S1b), underscoring the highly fibrotic nature of PDAC. The dense fibrosis of solid tumors is known to create a rigid TME. Nanoindentation measurements confirmed that human PDAC samples exhibited markedly higher Young’s modulus (1-7 kPa), compared to adjacent normal tissues (0.15-2 kPa; Fig. 1c and d). These stiffness characteristics were also observed in genetically engineered PDAC mouse models (KPC mouse models; Fig. 1d). Furthermore, Pearson correlation analysis revealed a positive correlation (*R* = 0.69; *P* = 0.027) between tissue stiffness and the degree of fibrosis (Fig. 1e), highlighting the strong association between the significant fibrosis in PDAC and its stiff TME.

To further explore the impact of the stiff TME, we performed a spatial transcriptome (ST) analysis on human PDAC and adjacent normal samples. Previous scRNA-seq data from human normal pancreas and PDAC was used for ST cell type deconvolution (29). We found that the signature scores for pathways related to extracellular matrix assembly, response to mechanical stimulus, and regulation of cell differentiation were significantly higher in PDAC tissues compared to adjacent normal tissues (Fig. 1f). Spatially, these pathways exhibited consistent distribution patterns across PDAC tissues (Fig. 1g). Spearman correlation analysis revealed a positive relationship among these pathways (Fig. 1h), underscoring the close association between ECM deposition, cell response to mechanical stimulus and cell differentiation.

Further analysis of cell types within the spatial transcriptome data revealed insights into how different cells contributed to and reacted to the stiff TME (Supplementary Fig. S2g). As expected, fibroblasts, which are major producers of extracellular matrix and are responsive to a rigid TME (30,31) (Fig. 1g). Sequential IHC results confirmed a significant increase in myofibroblast (FAP^+^ cells) presence in PDAC compared to adjacent normal tissues (Fig. 1i; Supplementary Fig. S3a and b).

Notably, macrophages were also predominantly found in these regions (Fig. 1g), suggesting their location in the stiff TEM and their capability to respond to mechanical signals. Indeed, the mechanical markers such as Lamin A/C and phosphorylated myosin II (p-MLC2) were elevated in macrophages (CD68^+^) in PDAC compared to adjacent normal tissues (Fig. 1i). In addition, a much higher abundance of macrophages was observed in the stiff TME (Fig. 1g and i), leading us to speculate that a stiff TME promotes differentiation of monocytes into macrophages.

To determine whether mechanical stimuli are required in monocyte differentiation into macrophage, we mixed either a soft (∼300 Pa) or stiff hydrogel (∼30 kPa) with KPC cells and orthotopically implanted into the pancreas of myeloid tdTomato^+^ mice to simulate a mild or intense mechanical stimuli (Fig. 1j). As a result, we observed a significantly higher number of macrophages (F4/80^+^ cells) in mice implanted with stiff hydrogel compared to the mice implanted with soft hydrogel (Fig. 1k). Moreover, over 80% of these macrophages were tdTomato positive, indicating their myeloid origin within pancreatic tissue (Fig. 1k). Of the macrophages, ∼6% were resident macrophages (LYVE1^+^F4/80^+^; Supplementary Fig. S4a and b), while in human PDAC, resident macrophages (LYVE1^+^CD68^+^) comprised less than 5% of macrophage population (Fig. 1i). These findings indicate that macrophages in PDAC tissues primarily arise from myeloid progenitors and that a stiff TME promotes the differentiation of these myeloid progenitors into macrophages.

To further explore the underlying mechanism, we isolated mouse bone marrow (BM) progenitors and differentiated them into macrophages under varying degrees of mechanical stimulation (Fig. 1l). Cells seeded on soft hydrogel underwent mild mechanical training, whereas cells cultured on stiff hydrogel and TCPS experienced intensive mechanical training. Bulk RNA-seq analysis revealed that BM progenitors exhibited higher expression of common progenitor markers such as *Flt3* and *Kit* (Fig. 1m). Cells subjected to mild mechanical training expressed monocyte markers including *Ly6c1* and *Ly6c2*. In contrast, cells exposed to intensive mechanical training showed significantly increased expression of mechanically related genes (*Piezo1*, *Trpv4*, *Itgb1*, *Lmna* and *Rhoa*), as well as elevated levels of the macrophage marker *CD68* (Fig. 1m; Supplementary Fig. S5). Gene set variation analysis (GSVA) indicated a higher enrichment of gene sets related to cell mechanotransduction, including actin-mediated cell contraction, actomyosin structure organization, and regulation of integrin-mediated signaling pathways, as well as monocyte differentiation, in cells subjected to intensive mechanical training compared to those subjected to mild training (Fig. 1n). These results indicate that monocyte differentiated into macrophages require intensive mechanical training induced by a stiff TME. To validate these findings, we analyzed BM progenitors cultured on various stiff substrates using flow cytometry. Consistently, an increase in matrix stiffness corresponded with a higher number of macrophages (Fig. 1o; Supplementary Fig. S6a and b), highlighting the necessity of intensive mechanical training for progenitor cells to differentiate into macrophages.

### Spatiotemporal analysis reveals PYK2 upregulation in monocytes in response to mechanical stimuli in human PDAC

To pinpoint the key factors involved in sensing and responding to mechanical stimuli during monocyte differentiation into macrophages in PDAC, we analyzed previously published sc-RNA seq data (32) (Supplementary Fig. S7a-c). This dataset consisted of 11 samples, comprising 26,420 cells, featuring both monocytes and macrophages from both the blood and tumor tissues of the same PDAC patient, effectively capturing the entire transition process from circulating monocytes to tumor-infiltrating and differentiating macrophages. Using a chi-square test, we observed that the majority of monocytes resided in the blood, while the majority of macrophages were present in the PDAC tissue, reflecting the pathophysiological process where monocytes circulate in the bloodstream and differentiate into macrophages within the TME (Supplementary Fig. S8a). We categorized monocytes and macrophages based on their tissue origin and identified three types of cells: “monocyte blood”, “monocyte PDAC”, and “macrophage PDAC” (Fig. 2a). Upon biochemical cues, blood monocytes infiltrate into tumor tissues, becoming “monocyte PDAC”, which then interact with mechanical cues from the TME, ultimately differentiating into “macrophage PDAC” cells. The trajectory analysis illustrated this dynamic transition process over time (Fig. 2b and c; Supplementary Fig. S8b).

**Fig. 2.**
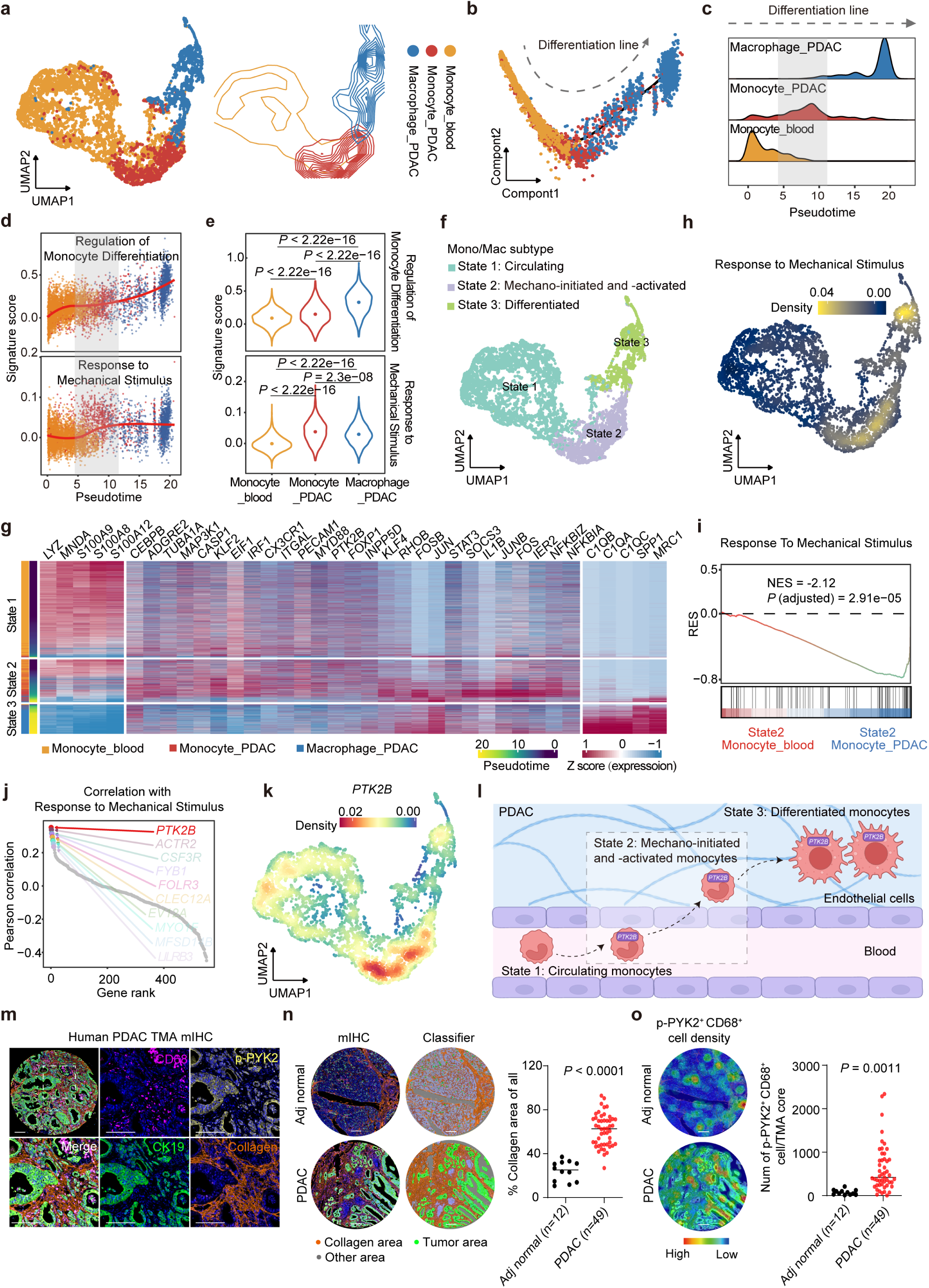
Spatiotemporal analysis of human PDAC reveals *PTK2B* is involved in monocyte mechanotransduction and differentiation. **a,** Uniform manifold approximation and projection (UMAP) visualization displaying cell types in scRNA-seq data of monocytes and macrophages in blood and PDAC tissues obtained from human PDAC patients(32). The left panel displays a UMAP plot where individual cells are represented as points, colored according to their cell type. The right panel presents a density plot based on the UMAP results, illustrating the density distribution of the cells. **b,** Pseudotime trajectory visualization for monocytes and macrophages from Fig. 2a in a two-dimensional state-space as determined by Monocle 2. The differentiation line (dashed) indicates the trajectory of this cellular transformation. **c,** The ridge plots showing the temporal distribution of each cell type from Fig. 2a along the pseudotime axis. The dashed line indicates the differentiation process. **d,** The plots showing the dynamic signature scores of two pathways, regulation of monocyte differentiation (top) and response to mechanical stimulus (bottom), along the pseudotime axis. The color of the dots represents the cell types, consistent with Fig. 2a. The red lines indicate the dynamic expression trends of the pathways, determined using a locally estimated scatterplot smoothing (LOESS) method. The grey area highlights the transition zone between blood monocytes and tumor-associated monocytes. **e,** The violin plots comparing the activity of the pathways, e.g., regulation of monocyte differentiation (top) and response to mechanical stimulus (bottom) across three indicated cell types. The significance of differences between the groups was assessed using t-tests. **f,** The UMAP plot illustrating the clustering results of three Mono/Macro cell types collected previously, using the original Louvain algorithm. Cells were divided into three groups: State 1 (light blue), representing circulating cells; State 2 (purple), representing mechano-initiated and -activated cells; and State 3 (light green), representing differentiated cells. **g,** The heatmap illustrating the dynamic changes in the expression of characteristic genes for monocytes and macrophages, as well as genes involved in the pathways (regulation of monocyte differentiation and response to mechanical stimulus) across the three cell states over pseudotime. **h,** The UMAP plot depicting the density distribution of cells based on their activity in the pathway of response to mechanical stimulus. Higher densities of active cells are shown in yellow, while lower densities are shown in blue. **i,** Gene set enrichment analysis (GSEA) of genes in response to mechanical stimulus pathway ranked by log2 fold change comparing monocytes from blood (red) with monocytes from PDAC (blue) in State 2. **j,** The dotplot displaying the ranking results of the Pearson correlation between the upregulated genes in blood-derived monocytes from state 2 and the activity in the pathway of the response to mechanical stimuli, highlighting the top 10 genes. **k,** The UMAP visualization of the density distribution of *PTK2B* expressed cells. Higher densities of *PTK2B*-expressed cells are shown in red, while lower densities are shown in blue. **l,** Schematic illustrating the infiltration of monocytes from the bloodstream into PDAC tissue, where they differentiate into macrophages. Initially, circulating monocytes breach the endothelial barrier and penetrate the PDAC tissue, triggering the upregulation of *PTK2B* and their response to mechanical cues. Subsequently, these mechano-activated monocytes differentiate into macrophages within the PDAC tissue. **m**, Representative images of mIHC staining of CK19, CD68, p-PYK2 and Collagen I in human PDAC tissue microarray (TMA). Scale bars, 200 μm. **n**, mIHC images and their corresponding digital images generated by HALO-based digital classifier, identifying the collagen, tumor, and background regions (left). Quantification of collagen area in human adjacent normal (n = 12) and PDAC (n = 49) samples using HALO software. Scale bars, 200 μm. **o**, Representative digital images (left) and quantification (right) of cell density of CD68^+^p-PYK2^+^ cells in adjacent normal (n = 12) and PDAC (n = 49) tissues. Scale bars, 200 μm. *P* values in (**n**) and (**o**) were determined by a two-tailed unpaired Student’s t-test.

Pathway analysis over pseudotime revealed that the signature scores for the regulation of monocyte differentiation progressively increased from blood monocytes to PDAC monocytes and further to PDAC macrophages (Fig. 2d and e). Additionally, the signature score for the response to mechanical stimuli showed an incremental trend from blood monocytes to PDAC but remained constant in PDAC macrophage (Fig. 2d and e). Pearson correlation analysis further supported these findings, indicating a significant positive correlation between these pathways and pseudotime progression (Supplementary Fig. S8c). These results suggest that the cellular response to mechanical stimuli is initiated in monocytes as they enter TME, driving their differentiation into macrophages.

Based on their mechanical response and differentiation status, these cells were classified into three distinct states: State 1 (circulating monocytes from the blood), State 2 (mechanically initiated and activated monocytes from both blood and PDAC), and State 3 (fully differentiated macrophages) (Fig. 2f). Gene expression profiles corroborates these states: State 1 exhibited monocyte markers such as *S100A8* and *S100A9*, State 2 showed transcription factors involved in monocyte differentiation like *FOS*, *JUN*, *STAT3*, and genes responding to mechanical stimuli such as *IL1B* and *ITGAL*, while State 3 expressed mature macrophage markers including *C1QA*, *C1QB*, and *C1QC* (Fig. 2g; Supplementary Table 2).

Focusing on State 2, we noted that this population, comprising a mix of blood and PDAC monocytes, represents the early stage of mechanical cue sensing and differentiation initiation. This population included approximately equal proportions of blood-derived monocytes and PDAC monocytes, along with a small number of macrophages from the PDAC tissue (Supplementary Fig. S8d). The UMAP plot of pathways for regulation of monocyte differentiation and response to mechanical stimuli showed activation beginning in State 2 (Fig. 2h; Supplementary Fig. S8e).

To identify key regulatory genes, we performed a differential expression genes (DEGs) analysis comparing blood and tumor-derived monocytes within this state. Notably, blood-derived monocytes in State 2 exhibited higher expression of genes related to actin filament organization, such as *ITGAL*, *ACTB*, *ACTR2*, and *MYL6*, while tumor-associated monocytes had increased expression of genes linked to monocyte differentiation, such as *FOS*, *JUNB*, *STAT3*, and *MRC1* (Supplementary Fig. S8f). GSEA analysis indicated that blood-derived monocytes were enriched in pathways related to actin filament function, whereas tumor-associated monocytes showed enrichment in pathways associated with mechanical stimuli response, myeloid differentiation, and macrophage activation (Fig. 2i; Supplementary Fig. S8g-h). These findings suggest a stepwise process where blood-derived monocytes first respond to mechanical stimuli by upregulating cytoskeletal genes and, upon entering the TME, initiate differentiation by upregulating genes associated with mechanical forces and macrophage maturation.

Among the DEGs linked to mechanical stimuli response, *PTK2B* (encoding PYK2) emerged as the top gene associated with this pathway (Fig. 2j; Supplementary Fig. S8i). *PTK2B* expression was highest in State 2, suggesting it plays a critical role in the initial response to mechanical stimuli, guiding monocyte mechano-activation (Fig. 2k-l). To validate these findings, we performed mIHC staining on a human PDAC tissue microarray (TMA) (Fig. 2m). Approximately 60% of the stained area was collagen-positive, indicating a stiff TME in PDAC (Fig. 2n). Moreover, macrophages in human PDAC samples exhibited significantly higher levels of active PYK2 compared to those in adjacent normal tissues (Fig. 2o). Notably, these p-PYK2^+^ macrophages were predominantly localized in the fibrotic regions (Fig. 2o), reinforcing PYK2’s critical involvement mediating cellular responses to mechanical cues within the TME.

### PYK2 is critical for mechanical training of monocyte differentiation

To experimentally validate whether *PTK2B* was involved in sensing and responding to mechanical stimuli from the environment, THP-1 cells, a widely used human monocytic cell line in monocyte/macrophage research, were primed with PMA and attached to a matrix to undergo mechanical training. Upon attachment, the cells received mechanical training. We observed a significant increase in *PTK2B* expression in cells subjected to mechanical training compared to suspended cells without training (Fig. 3a). As the duration of training progressed, cell stiffness increased from 24 to 48 hours, reaching a consistent level at 72 hours (Fig. 3b). Notably, the protein expression and activation of *PTK2B* followed a similar pattern, continuously increasing up to 48 hours and then plateauing at 72 hours (Fig. 3a).

**Fig. 3.**
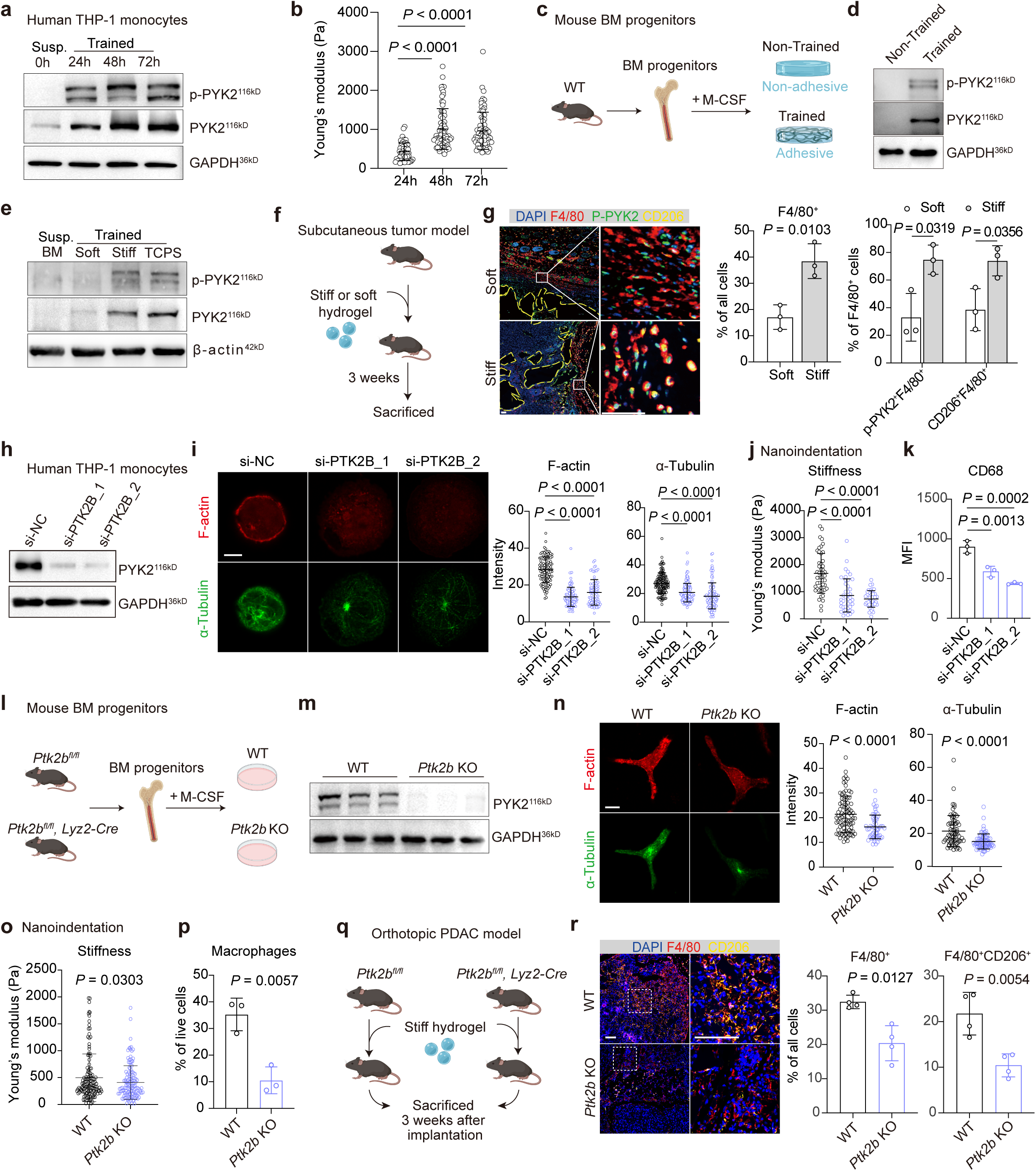
PYK2 serves as an immunomechanical checkpoint coordinating mechanical training of monocyte differentiation. **a**, Western blot analysis of PYK2 and p-PYK2 in suspended THP-1 cells (non-trained), and the cells subjected to mechanical training for 24, 48 and 72 hours. **b**, Assessment of THP-1 cell stiffness following mechanical training durations of 24 (n=55), 48 (n=75), and 72 (n=75) hours. **c**, Schematic illustrating the differentiation of mouse BM progenitors into macrophages, with and without mechanical training. Mouse BM progenitors were isolated and cultured with M-CSF on non-adhesive and adhesive surfaces for 3 days. **d**, Western blot analysis of PYK2 and p-PYK2 in differentiated BM progenitors after 3 days of mechanical training compared to those without training. **e**, Western blot analysis of PYK and p-PYK2 in mouse BM progenitors and differentiated BM progenitors after 6 days of mechanical training on soft, stiff and TCPS surfaces. **f**, Schematic of establishment of subcutaneous tumor model. Stiff or soft hydrogel were mixed with KPC cells and subcutaneously injected into wildtype mice. The mice were sacrificed 3 weeks after implantation. **g**, Representative images (left) and quantification (middle and right) of mIHC staining of F4/80, p-PYK2, CD206 and DAPI of tissues collected 3 weeks after subcutaneous implantation. The location of the hydrogel is indicated by a dashed yellow line. n = 3 mice in each group. Scale bars, 50 μm. **h**, Validation of *PTK2B* silencing in THP-1 cells using RNA interference (RNAi) through Western blot analysis. THP-1 cells were transfected with si-NC (negative control), si-PTK2B_1, and si-PTK2B_2, and stimulated with PMA for 48 hours. Subsequently, the cells were lysed for protein extraction and underwent western blot analysis. **i**, Representative immunofluorescent images (left) and quantification (right) of F-actin and α-tubulin in THP-1 cells with silenced *PTK2B*. Cells were transfected with si-NC, si-PTK2B_1, and si-PTK2B_2, followed by 48-hour PMA stimulation. n = 117 (si-NC), 108 (si-PTK2B_1), and 79 (si-PTK2B_2) cells for F-actin quantification. n = 131 (si-NC), 142 (si-PTK2B_1), and 111 (si-PTK2B_2) cells for tubulin quantification. Scale bars, 10 μm. **j**, Assessment of stiffness of THP-1 cell following *PTK2B* silencing. THP-1 cells were transfected with si-NC, si-PTK2B_1, and si-PTK2B_2, then stimulated with PMA for 48 hours. Cell stiffness was analyzed using nanoindentation testing. n = 49, 43 and 41 cells for si-NC, si-PTK2B_1 and si-PTK2B_2 conditions, respectively. **k**, Flow cytometry analysis of THP-1 cells following *PTK2B* silencing. THP-1 cells were transfected with si-NC, si-PTK2B_1 and si-PTK2B_2, followed by stimulating with PMA for 48h. MFI of CD68 of THP-1 cells gated on live single cells. **l**, Schematic of cultivation process of *Ptk2b* knockout (KO) BMDMs. Mouse BM cells were isolated from *Ptk2b^fl/fl^* (WT) and *Ptk2b^fl/fl^, Lyz2-Cre* (*Ptk2b* KO) mice, and subsequently stimulated with M-CSF for 6 days. **m**, Validation of *Ptk2b* knockout (KO) in BMDMs using western blot. **n**, Representative immunofluorescent images and quantification of F-actin and tubulin in WT and *Ptk2b* KO BMDMs. n = 86 (WT) and 57 (*Ptk2b* KO) cells for F-actin quantification. n = 71 (WT) and 76 (*Ptk2b* KO) cells for tubulin quantification. Scale bars, 10 μm. **o**, Stiffness determination of WT (n = 137) and *Ptk2b* KO (n = 133) BMDMs using nanoindentation testing. **p**, Flow cytometry analysis of macrophages derived from WT and *Ptk2b* KO BM progenitors. Macrophages were gated on CD45^+^ CD11b^+^ Ly6G^-^ Ly6C^-^ F4/80^+^ single live cells. **q**, Schematic of establishment of orthotopic PDAC model in *Ptk2b^fl/fl^* and *Ptk2b^fl/fl^, Lyz2-Cre* mice. Stiff or soft hydrogel were mixed with KPC cells and orthotopically injected into the pancreas of *Ptk2b^fl/fl^* or *Ptk2b^fl/fl^, Lyz2-Cre*. The mice were sacrificed 3 weeks after implantation. **r,** Representative images (left) and quantification (middle and right) of mIHC staining of F4/80, CD206, and DAPI of tissues collected 3 weeks after orthotopically pancreatic implantation in *Ptk2b^fl/fl^* and *Ptk2b^fl/fl^, Lyz2-Cre* mice. n = 4 mice in each group. Scale bars, 100 μm. *P* values in (**g**), (**n**), (**o**) and (**p**) were determined by two-tailed unpaired Student’s t-test; *P* values in (**a**), (**i**), (**j**) and (**k**) were determined by ordinary one-way ANOVA. Data shown in (**b**), (**i**), (**j**), (**k**), (**n**), (**o**) and (**p**) are representative of three experiments.

To further confirm the mechanical response of PYK2, mouse BM progenitors were primed with M-CSF and cultured on adhesive and non-adhesive matrices (Fig. 3c). The adhesive surface, composed of polyethylene glycol (PEG) and gelatin methacryloyl (GelMA), allowed cell adhesion and subsequent perception of mechanical stimuli upon substrate attachment. In contrast, the non-adhesive surface consisting solely of PEG, prevented cell adhesion. Cells cultured on adhesive matrices and subjected to mechanical training exhibited markedly increased protein expression of *Ptk2b* compared to non-trained cells (Fig. 3d). Furthermore, the protein expression of *Ptk2b* and its phosphorylation changed in accordance with the intensity of the mechanical training, with higher levels of expression and activation observed under more intense training conditions (Fig. 3e).

To investigate this phenomenon *in vivo*, we prepared stiff and soft hydrogels, combined them with KPC cells, and subcutaneously implanted them into mice (Fig. 3f; Supplementary Fig. S9a and b). As expected, the stiff TME (∼2 kPa), created by stiff hydrogels, exhibited a higher abundance of macrophages (F4/80^+^) (Fig. 3g; Supplementary Fig. S9c). Additionally, macrophages in the stiff TME exposed to stronger mechanical stimuli showed elevated levels of phosphorylated PYK2 (p-PYK2) and a shift toward an alternatively activated phenotype, indicated by increased CD206 expression (Fig. 3g). These findings collectively suggest that PYK2 can detect and respond to mechanical stimuli from the microenvironment both *in vitro* and *in vivo*.

To further elucidate the role of PYK2 in mechanotransduction and monocyte differentiation, we used small interfering RNA (siRNA) to silence *PTK2B* expression in THP-1 cells. Western blot analysis confirmed effective *PTK2B* silencing, with PYK2 inhibition observed without affecting FAK expression (Fig. 3h; Supplementary Fig. S10a). We then employed immunofluorescence staining to evaluate F-actin and microtubules, key components of the cytoskeleton responsible for cellular force generation (Fig. 3i). In *PTK2B* knockdown THP-1 cells, the assembly of F-actin and expression of α-tubulin were markedly reduced (Fig. 3i), leading to a significant decrease in cellular force generation (Fig. 3j) and impairing macrophage maturation (Fig. 3k).

To explore the effects of endogenous PYK2 elimination on monocytes and macrophages, we generated myeloid-specific *Ptk2b* knockout (KO) mice (*Ptk2b^fl/fl^, Lyz2-Cre*) (Fig. 3l and m; Supplementary Fig. S10b). Consistently, the absence of PYK2 markedly disrupted the organization of F-actin and microtubules (Fig. 3n), resulting in reduced cellular stiffness (Fig. 3o) and a significant decrease in macrophage number (Fig. 3p).

To confirm these findings *in vivo*, stiff hydrogels were combined with KPC cells and implanted orthotopically into the pancreas of myeloid *Ptk2b* KO mice (*Ptk2b^fl/fl^, Lyz2-Cre*) or control mice (*Ptk2b^fl/fl^*; Fig. 3q). PDAC tissue from control mice exhibited a significantly higher presence of total macrophages (F4/80^+^), alongside elevated p-MLC2^+^ F4/80^+^, Lamin A/C^+^ F4/80^+^, and NF-κB^+^ F4/80^+^ subsets, indicating PYK2 deficiency disrupts the mechanotransduction and monocyte differentiation (Fig. 3r; Supplementary Fig. S11a and b). Additionally, there was a notable shift of macrophages toward the alternative activated phenotype (F4/80^+^ CD206^+^) in control mice compared to KO mice, indicating the potential role of PYK2 in modulating macrophage polarization (Fig. 3r).

To further confirm these observations, we exposed *Ptk2b* KO or WT bone marrow-derived macrophages (BMDMs) to LPS, IL-4/IL-13, or a conditioned medium in a stiff microenvironment. *Ptk2b* KO cells exhibited a more inflammatory phenotype, characterized by increased expression of iNOS and TNF-α in response to LPS compared to control cells. Conversely, control cells showed significantly higher levels of alternatively activated macrophage markers such as CD206, Arg-1, and CD163 in response to IL-4/IL-13 or conditioned medium (Supplementary Fig. S12a and b), suggesting that PYK2-mediated mechanotransduction also influences macrophage polarization. Furthermore, *Ptk2b* knockout BMDMs displayed a significantly reduced ability to migrate (Supplementary Fig. S13a-c). These findings underscore the crucial role of PYK2-dependent mechanical training in monocyte differentiation and macrophage function.

### Nuclear PYK2 regulates the transcriptional program of monocyte differentiation upon mechanical training

PYK2, a calcium-sensitive tyrosine kinase, is known for its activation by Ca^2+^ and calmodulin (33). Given that Piezo1 is a well-known mechanical stress-sensitive ion channel that influences macrophage maturation and inflammatory function (23,26,34), we thus investigated whether Piezo1-mediated Ca^2+^ influx contributes to PYK2 activation in monocytes. We first examined the effect of the Piezo1 agonist, Yoda1, on PYK2 activation. Western blot analysis revealed a dose-dependent increase in PYK2 activation, which was effectively blocked in the presence of the Ca^2+^ chelator EGTA (Fig. 4a). Interestingly, EGTA treatment also led to a decrease in PYK2 protein expression (Fig. 4a), possibility due to a weak cell attachment induced by EGTA. To further explore this relationship, we monitored cytosolic Ca^2+^ levels using the Ca^2+^ indicator Fluo-4/AM in both control and *PIEZO1* knockdown THP-1 cells following Yoda1 treatment. Consistent with previous study(26), Yoda1 treatment resulted in a substantial rise in cytosolic Ca^2+^ levels in control cells (Fig. 4b). In contrast, PIEZO1 deficient cells exhibited only a marginal increase in cytosolic Ca^2+^ levels when exposed to Yoda1 (Fig. 4b), confirming the role of Piezo1 in mediating cytosolic Ca^2+^ influx. Notably, silencing *PIEZO1* led to reduced PYK2 activation (Fig. 4c), underscoring the significance of Ca^2+^ influx via Piezo1 in triggering PYK2 activation. Flow cytometry further suggested that Yoda1-induced Piezo1 activation enhanced monocyte-to-macrophage differentiation, while Piezo1 inhibition with GdCl₃ suppressed it (Supplementary Fig. S14a and b), positioning PIEZO1 as a mechanosensitive regulator upstream of PYK2-mediated differentiation.

**Fig. 4.**
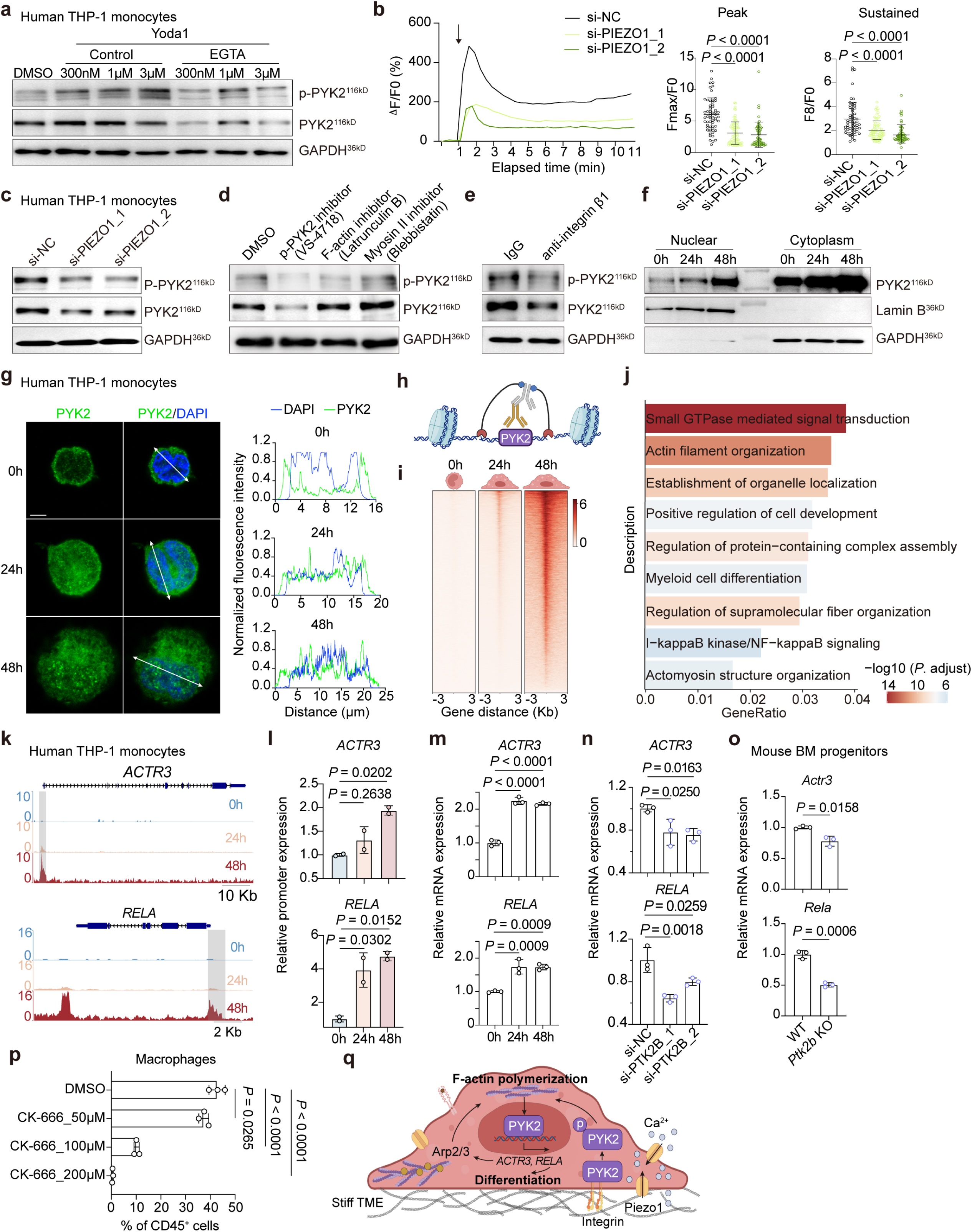
Nuclear PYK2 guides monocyte differentiation at transcriptional level following mechanical training. **a,** Western blot analysis of PYK2 and p-PYK2 in THP-1 cells treated with PMA and Yoda1 (300 nM) in the presence or absence of EGTA (4 mM) for 6 hours. **b,** Evaluation of cytosolic Ca^2+^ levels in THP-1 cells transfected with si-NC or si-PIEZO1. Following transfection of THP-1 cells with si-NC, si-PIEZO1_1, or si-PIEZO1_2, cells were primed with PMA for 48 hours. Stimuli Yoda1 was added into the culture medium at 1 min as indicated by a black arrow. Fluorescence intensities of calcium indicator Fluo-4 AM in individual cells were tracked over time. The relative fluorescence intensities (ΔF/F0) of Fluo-4 AM-loaded THP-1 cells were calculated (left), while the average maximum peak intensity (Fmax/F0) and the intensity at the steady state (F8/F0) were calculated at 7 min post Yoda1 stimulation from si-NC (58 cells), si-PIEZO1_1 (75 cells) and si-PIEZO1_2 (58 cells) treated cells (middle and right). **c,** Western blot analysis of PYK2 and p-PYK2 in THP-1 cells following PIEZO1 silencing. THP-1 cells were transfected with si-NC, si-PIEZO1_1 or si-PIEZO1_2, followed by treatment of PMA for 48 hours. **d-e,** Western blot analysis of PYK2 and p-PYK2 in THP-1 cells treated with PMA and anti-integrin β1 antibody (5 µg/ml), or inhibitors including PYK2 inhibitor VS-4718 (1 µM), F-actin polymerization inhibitor Latrunculin B (5 µM) and Myosin II inhibitor Blebbistatin (10 µM) for 24 hours. IgG antibody or DMSO served as controls. **f,** Western blot analysis of PYK2 in the nuclear or cytoplasmic fractions of suspended THP-1 cells and those primed with PMA for 24h and 48h, respectively. **g,** Investigation of the colocalization of PYK2 with the nucleus in suspended THP-1 cells and those primed with PMA for 24h and 48h, respectively. Nucleus were labeled with DAPI. Representative immunofluorescence images of PYK2 (green) and DAPI (blue) in THP-1 cells are presented (left). Histogram of the normalized fluorescent intensity of PYK2 and DAPI was obtained from the cross-sectional lines through cell nucleus (right). Scale bars, 10 μm. **h,** Schematic of CUT&Tag for investigating the chromatin recruitment of PYK2. **i,** CUT&Tag sequencing analysis of PYK2 in suspended THP-1 cells and those primed with PMA for 24h and 48h, respectively. **j,** The bar plot showing GO enrichment analysis of downstream genes associated with promoter regions (<1 kb) identified from CUT&Tag data in mechanically trained THP-1 cells (48h vs 0h). The color gradient represents the -log10 adjusted *P* value, and the x-axis indicates the GeneRatio. **k,** CUT&Tag tracks of *ACTR3* and *RELA* loci in suspended THP-1 cells and those primed with PMA for 24h and 48h. **l,** qPCR validation of *ACTR3* and *RELA* loci in suspended THP-1 cells and those primed with PMA for 24h and 48h. **m**, Relative mRNA expression of *ACTR3* and *RELA* in suspended THP-1 cells and those primed with PMA for 24h and 48h. **n,** Relative mRNA expression of *ACTR3* and *RELA* in THP-1 cells following *PTK2B* silencing. **o,** Relative mRNA expression of *ACTR3* and *RELA* in WT or *Ptk2b* KO mouse progenitors primed with M-CSF for 3 days. **p,** Flow cytometry analysis of mouse progenitors treated with M-CSF and Arp2/3 inhibitor CK-666 (50, 100 and 200 µM) for 3 days. **q,** Illustration outlining the PYK2-mediated mechanotransduction and differentiation process of monocytes within a stiff microenvironment. Monocytes respond to biochemical cues by interacting with their environment, initiating the differentiation process. Upon adhesion to the rigid matrix, activation of Piezo1 and integrin occurs, leading to the activation of PYK2 and subsequent reorganization of F-actin. Following this, PYK2 translocates to the nucleus, and selectively binds to gene regulatory regions associated with actin cytoskeleton organization and NF-κB signaling, such as *ACTR3* and *RELA*, and others, thereby promoting monocyte differentiation. *P* values in (**o**) were determined by two-tailed unpaired Student’s t-test; *P* values in (**b**), (**l**), (**m**), (**n**) and (**p**) were determined by ordinary one-way ANOVA. Data shown in (**i**), (**k**) and (**l**) are representative of two experiments. Data shown in (**b**), **(m)**, (**n**), (**o**) and (**p**) are representative of three experiments.

The F-actin cytoskeleton serves as the primary mechanotransducer (35), prompting us to investigate the contributions of F-actin polymerization and actomyosin contractility to PYK2 expression and activation. Utilizing VS-4718, a well-known inhibitor of PYK2, as a positive control, we observed a significant reduction in both PYK2 protein expression and activation after 24-hour treatment (Fig. 4d). Subsequent experiments showed that inhibiting F-actin polymerization with Latrunculin B effectively suppressed PYK2 phosphorylation, while inhibition of myosin II activity did not show a significant impact on PYK2 expression and activation (Fig. 4d). Since F-actin is linked to integrins in terms of its organization and dynamics (36), blocking integrin β1 using an anti-integrin β1 antibody resulted in a substantial reduction in both PYK2 expression and activation levels (Fig. 4e). These results suggest that F-actin polymerization, rather than the actomyosin contractility, is the primary factor influencing PYK2 expression and activation.

During monocyte differentiation into macrophage under mechanical training, PYK2 was observed to translocate from the cytoplasm to the nucleus (Fig. 4f and g). Although previous studies have identified the nuclear presence of PYK2, its specific role in this location remains incompletely understood. To fully uncover the genomic maps of PYK2-associated DNA-binding sites in monocytes and macrophages, we conducted CUT&Tag sequencing using a PYK2 antibody in THP-1 cells with or without mechanical training (Fig. 4h). As expected, the chromatin accessibility of PYK2 increased with the process of monocyte differentiation (Fig. 4i). Analysis of promoter regions (<1 kb) with signals detected in the 48h vs 0h CUT&Tag data from mechanically trained THP-1 cells revealed significant enrichment of downstream genes in pathways related to cytoskeleton dynamics, such as small GTPase-mediated signal transduction, actin filament organization, and actomyosin structure organization, as well as NF-κB signaling, which guides monocyte differentiation (37,38) (Fig. 4j; Supplementary Table 3).

Specifically, PYK2 was found to bind to the promoter region of *ACTR3*, an essential component of the Arp2/3 complex involved in actin polymerization (39,40), and *RELA*, a canonical NF-κB signaling pathway component (41,42) (Fig. 4k). Binding to these protomer regions was validated by qPCR (Fig. 4l-o). Mechanical training of THP-1 monocytes resulted in increased expression of *ACTR3* and *RELA* (Fig. 4m). Conversely, PYK2 loss in THP-1 cells or knockout in mouse monocytes significantly reduced mRNA and protein expression of these genes (Fig. 4n and o; Supplementary Fig. S15a). Furthermore, inhibition of the activity of Arp2/3 by CK-666 induced dose-dependent inhibition of macrophage differentiation (Fig. 4p), underscoring the mechanistic link between PYK2-mediated actin dynamics and differentiation.

We next analyzed the binding motifs of PYK2 using the HOMER motif discovery tool (43). This revealed significant enrichment of AP-1 (e.g., Fra2, Fra1, Jun, JunB, Fos) and SP-1 motifs at PYK2-binding sites (Supplementary Fig. S15b), suggesting that PYK2 collaborates with AP-1 and/or SP-1 to regulate gene expression. Focusing on the RELA promoter, we identified eleven SP-1 and eight AP-1 motifs within 1 kb upstream of its transcription start site (Supplementary Fig. S15c). To test this interaction functionally, we cloned a 286 bp fragment of the RELA promoter containing these motifs (RELA response element, RE) into a luciferase reporter vector and transfected it into 293T cells. PYK2 overexpression increased luciferase activity by approximately 2-fold compared to controls (Supplementary Fig. S15d), indicating that PYK2 enhances RELA transcription via these motifs. A DNA pull-down assay further confirmed this, using a 5’-biotinylated DNA probe spanning the AP-1 and SP-1 binding regions of the RELA promoter. PYK2, alongside AP-1 and SP-1, co-precipitated with these sequences in the nuclear extracts from THP-1-derived macrophages and PYK2-overexpressing 293T cells, with specificity validated by a 10-fold excess of non-biotinylated competitor oligonucleotides disrupting binding (Supplementary Fig. S15e). These data suggest that PYK2 physically associates with AP-1 and/or SP-1 at these sites, though additional studies are needed to define the precise protein-protein interactions driving RELA transcription.

Collectively, these findings identify PYK2 as a key regulator of mechanotransduction and differentiation. Upon biochemical stimulation with M-CSF, monocytes attach to their surroundings and undergo mechanical training, activating mechanosensitive ion channels and integrins. This process activates PYK2 and promotes F-actin polymerization, leading to the nuclear translocation of PYK2. PYK2 then drives the transcription of genes crucial for mechanotransduction and differentiation, creating a positive feedback loop that facilitates monocyte differentiation (Fig. 4q).

### Loss of PYK2 in myeloid precursors disrupts ECM-associated mechanics in TME and enhances the efficacy of anti-PD-1 immunotherapy

To explore the therapeutic implications of these findings, we examined whether the genetic loss of *Ptk2b* in monocytes could have a beneficial impact on PDAC. By crossing *Ptk2b^fl/fl^* mice with Lyz2-Cre mice, we generated *Ptk2b^fl/fl^* (WT); *Ptk2b^fl/+^, Lyz2-Cre* (Ptk2b HET), and *Ptk2b^fl/fl^, Lyz2-Cre* (Ptk2b HOM) mice. To replicate the stiff TME of PDAC, we created PDAC models by orthotopically implanting a small piece (∼1mm in diameter) of fibrotic tumor (Supplementary Fig. S16a) into the pancreas of mice (Fig. 5a). During the initial two weeks post-implantation, a dose-dependent reduction in tumor size was evident in myeloid PYK2-deficient mice (Fig. 5b). However, by the third week, tumor sizes in control and *Ptk2b* HET mice were comparable. *Ptk2b* HOM mice consistently exhibited a significant decrease (∼32 days) in tumor size compared to controls, although this difference diminished over time (Supplementary Fig. S16b). Masson’s trichrome staining, IHC analysis of α-SMA, collagen I, and fibronectin, and tumor stiffness assessments demonstrated a dose-dependent decrease in fibrosis (Fig. 5c-f; Supplementary Fig. S16c) and tumor stiffness (Fig. 5d), indicating that loss of PYK2 in BM progenitors disrupts ECM-related mechanics within the TME.

**Fig. 5.**
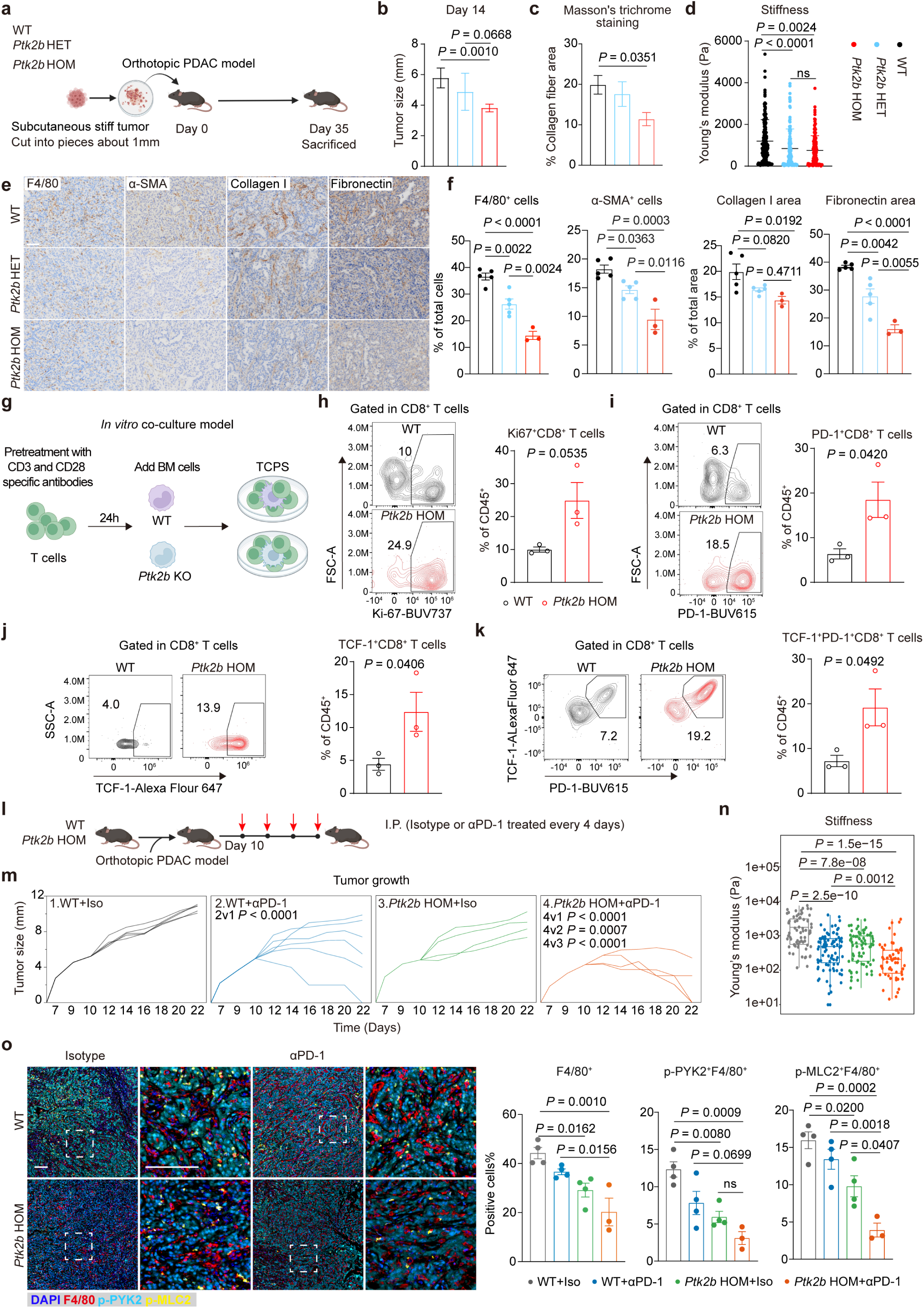
Loss of PYK2 in BM progenitors reduces mechanics in TME and enhances the efficacy of PD-1 immunotherapy. **a**, Schematic diagram of the experimental setup for **b-g**. KPC cells were subcutaneously injected into the wild-type mice, and the stiff tumor tissue was collected two weeks after the injection. The tumor was then sectioned into small pieces (∼1mm) and orthotopically transplanted into the pancreas of *Ptk2b^fl/fl^* (WT), *Ptk2b^fl/+^, Lyz2-Cre* (*Ptk2b* HET) and *Ptk2b^fl/fl^, Lyz2-Cre* (*Ptk2b* HOM) mice. The mice were sacrificed 35 days after implantation. **b**, Tumor diameter of PDAC tissue in WT (n = 6), *Ptk2b* HET (n = 6) and *Ptk2b* HOM (n = 7) at day 14. **c**, Quantification of the fibrotic area stained using Masson’s trichrome staining in PDAC samples from WT (n = 6), *Ptk2b* HET (n = 6), and *Ptk2b* HOM (n=7) mice. **d**, Quantification of tissue stiffness of PDAC in WT (n = 6), *Ptk2b* HET (n = 5), and *Ptk2b* HOM (n=7) mice using nanoindentation. **e**, Representative images and (**f**) quantification of IHC staining of F4/80, Collagen I, α-SMA, and Fibronectin in PDAC in WT (n = 5), *Ptk2b* HET (n = 5) or *Ptk2b* HOM (n = 3) mice. Scale bars, 100 μm. **g,** Schematic of *in vitro* co-culture experiment. CD8^+^ T cells were isolated and stimulated with CD3 and CD28-specific antibodies overnight. WT or *Ptk2b* KO BM progenitors were isolated and cultured with T cells at a ratio of 1:5 with the presence of M-CSF. After a 7-day co-culture period, the cells were collected and analysed by flow cytometry. **h-k,** Flow cytometry analysis of CD8^+^ T cells co-cultured with WT or *Ptk2b* KO monocytes. **l**, Schematic diagram of the experimental setup for **m-o**. KPC cells were orthotopically transplanted into the pancreas of WT and *Ptk2b* HOM mice. The mice received either Isotype control or αPD-1 treatment (200 µg) every 3 days starting from day 10 when the tumor reached approximately 0.5 cm in diameter. The mice were sacrificed after four rounds of injections. **m,** Tumor growth curves of individual tumors in WT or *Ptk2b* HOM mice that received either Isotype control or αPD-1 treatment. n = 5 (WT+Iso), n = 6 (WT+αPD-1), n = 4 (*Ptk2b* HOM+Iso), n = 4 (*Ptk2b* HOM+αPD-1) mice. **n,** Quantification of tissue stiffness of PDAC in WT+Iso (n = 5), WT+αPD-1 (n = 5), and *Ptk2b* HOM+Iso (n = 4), *Ptk2b* HOM+αPD-1 (n = 3) mice using nanoindentation. **o,** Representative images (left) and quantification (right) of mIHC staining of F4/80, p-MLC2, p-PYK2, and DAPI in PDAC in WT or *Ptk2b* HOM mice that received either Isotype control or αPD-1 treatment. n = 4 (WT+Iso), n = 4 (WT+αPD-1), n = 4 (*Ptk2b* HOM+Iso), n = 3 (*Ptk2b* HOM+αPD-1) mice. Scale bars, 100 μm. *P* values in (**b-f**), (**n**), and (**o**) were determined by ordinary one-way ANOVA; *P* values in (**h-k**) were determined by two-tailed unpaired Student’s t-test; *P* values in (**m**) were determined by ordinary two-way ANOVA.

To comprehensively explore the impact of PYK2 loss in myeloid cells and lymphocytes, BM progenitors, monocytes, macrophages, and T cells were analyzed (Fig. 5e; Supplementary Fig. S17a-e). The deficiency of PYK2 in myeloid cells led to an increase in the number of progenitor cells, such as MDPs (Supplementary Fig. S17f), potentially leading to an elevation of monocytes in the bloodstream (Supplementary Fig. S17g). However, the macrophages showed a dose-dependent reduction in the PDAC tissue of PYK2 deficient mice (Fig. 5e), indicating that PYK2 loss influences monocyte differentiation in PDAC *in vivo*.

Analysis of T cells, which are crucial for tumor cell killing, revealed a tendency for an increased CD8^+^ T cells in PDAC of both *Ptk2b* HET and *Ptk2b* HOM mice, along with a concurrent tendency for a rise in exhausted PD1^+^ CD8^+^ T cells (Supplementary Fig. S17d and h). *In vitro* co-culture experiments showed that PYK2 deficiency in monocytes directly stimulated CD8^+^ T cell proliferation (Fig. 5g and h; Supplementary Fig. S18a and b). Although these CD8^+^ T cells eventually became exhausted (Fig. 5i), there were significantly increased expression levels of TCF-1 protein in T cells from *Ptk2b* HOM mice compared to control, suggesting T cell functionality was preserved due to the maintenance of its stemness (Fig. 5j and k). Additionally, we observed an increased number of T cells in Draining lymph nodes (DLN) from *Ptk2b* HOM mice, indicating PYK2 depletion in myeloid cells may enhance the priming of T cells in DLN (Supplementary Fig. S17e and i), while reduced tumor stiffness may facilitate the infiltration of these T cells into tumors. These findings indicate the potential benefit of combined αPD-1 antibody therapy.

To assess the impact of αPD-1 therapy, treatment was administered to both WT and *Ptk2b* HOM mice once tumors reached approximately 0.5 cm in size. Tumor size was monitored every other day, and mice were euthanized after receiving four injections (Fig. 5l). Tumors in control mice grew rapidly, whereas αPD-1 treatment slowed growth compared to the isotype control (Fig. 5m). Remarkably, tumors in *Ptk2b* HOM mice receiving αPD-1 treatment began to shrink after the second injection, with tumors nearly vanishing in half of the mice after four treatments (Fig. 5m; Supplementary Fig. S19a), highlighting the effectiveness of combining αPD-1 therapy.

Consistently, the fibrotic area and tissue stiffness were reduced in myeloid PYK2-deficient mice compared to controls (Fig. 5n; Supplementary Fig. S19b and c). Notably, αPD-1 treatment further intensified these effects, with a minimal fibrotic area (∼5%) and tissue stiffness restored to levels similar to normal tissues (∼300 Pa). These results underscore the significant alteration of ECM-related mechanics within the TME through the combination of myeloid PYK2 loss and αPD-1 treatment.

Further mIHC analysis revealed that the absence of PYK2 led to a significant decrease in macrophage populations and reduced myosin II activation. αPD-1 treatment in *Ptk2b* HOM mice further diminished macrophage numbers and myosin II activation in tumors (Fig. 5o). These findings suggest that the deficiency of PYK2 disrupts mechanotransduction in monocytes, impeding their differentiation, while αPD-1 treatment exacerbates this effect. Ultimately, these changes in differentiated monocytes reshape the TME of PDAC, offering a promising opportunity for improved PDAC treatment.

### Loss of PYK2 in myeloid precursors enhances cytotoxic CD8^+^ T cell response and extends mouse survival

To comprehensively explore the mechanisms underlying the improved efficacy of αPD-1 treatment upon myeloid PYK2 loss, we conducted scRNA-seq on tumor tissues from four cohorts: WT+Iso, WT+αPD-1, *Ptk2b* HOM+Iso, and *Ptk2b* HOM+αPD-1. After stringent quality control, transcriptomes from 47,124 high-quality single cells were obtained. Unsupervised clustering and canonical marker-based annotation identified 13 major cell types, revealing distinct distribution patterns of immune and malignant epithelial cells in each cohort (Fig. 6a; Supplementary Fig. S20a-f; Supplementary Table 4). Generally, there was a reduction in malignant epithelial cells in the *Ptk2b* HOM+Iso cohort compared to the WT+Iso cohort, and this reduction was further enhanced by the combination of αPD-1 treatment. Regarding changes in the tumor immune microenvironment (TIME), the *Ptk2b* HOM+Iso cohort showed an increase in monocytes and a decrease in macrophages compared to the control cohort, along with a notable increase in T cells. The αPD-1 treatment further augmented T cell infiltration (Fig. 6a). These changes in both tumor cells and TIME reinforce the beneficial outcomes of the combined therapy.

**Fig. 6.**
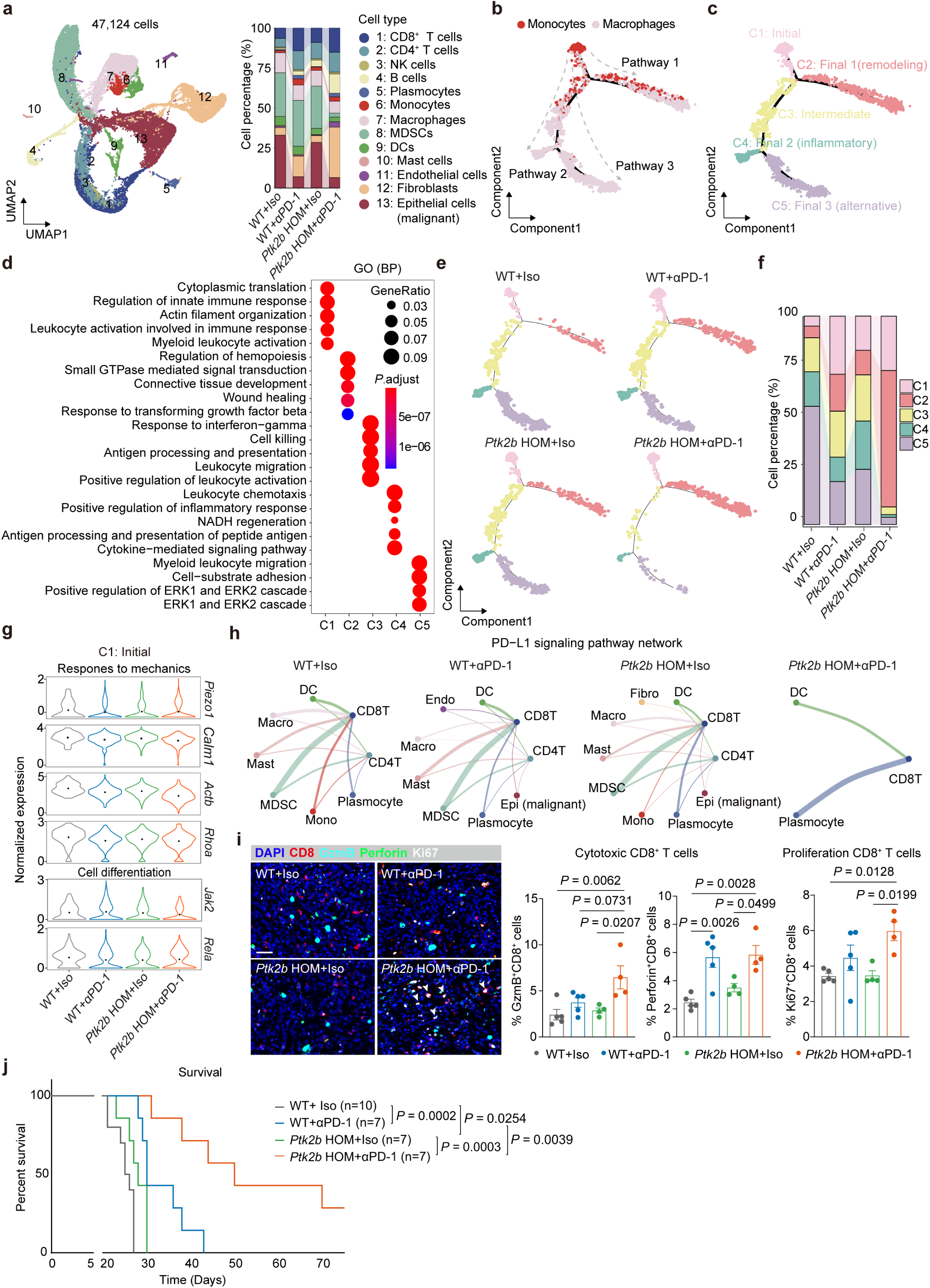
Loss of PYK2 in myeloid precursors improves effector T cell infiltration and extends mouse survival. **a**, Profiling of scRNA-seq data of WT+Iso, WT+αPD-1, *Ptk2b* KO+Iso, and *Ptk2b* KO+αPD-1 groups. The left panel represented the UMAP visualization displaying single-cell clustering of cells from all four groups, consisting of 47,124 cells from 8 samples. The identified cell types included CD8^+^ T cells (1), CD4^+^ T cells (2), NK cells (3), B cells (4), Plasmocytes (5), Monocytes (6), Macrophages (7), MDSCs (8), Dendritic Cells (9), Mast cells (10), Endothelial cells (11), Fibroblasts (12), and Malignant Epithelial cells (13). The right panel represented the stacked bar plot showing the variation in the proportion of these 13 cell types across the 4 groups. Each color represented a different cell type. **b,** Monocle analysis showed the developmental trajectory of monocytes and macrophages in WT+Iso, WT+αPD-1, *Ptk2b* KO+Iso, and *Ptk2b* KO+αPD-1 groups. The analysis identified three pathways, labeled Pathway 1, Pathway 2, and Pathway 3, with monocytes (red) transitioning into macrophages (pink) along distinct developmental pathways. **c,** Cell grouping and naming from Monocle trajectory analysis in Fig. 6b, identified five clusters: C1 (Initial), C2 (Final 1, remodeling), C3 (Intermediate), C4 (Final 2, inflammatory), and C5 (Final 3, alternative). **d,** GO terms and pathways enriched for up-regulated genes (adjusted *P* value < 0.05, log2 fold change > 0.25) in C1, C2, C3, C4, and C5 cells. The color scale indicates adjusted *P* values, derived from a hypergeometric test. The size of the symbols is proportional to the gene counts enriched in the corresponding GO terms. **e,** Monocle analysis of monocyte and macrophage development in WT+Iso, WT+αPD-1, *Ptk2b* KO+Iso, and *Ptk2b* KO+αPD-1 groups, corresponding to the clusters in Fig. 6c. **f,** Stacked bar chart showing the variation in the proportion of these five sub-cell types from monocle analysis across the four groups. Each color represents a different cell type. **g,** Violin plots displayed the expression levels of response to mechanics genes (*Piezo1*, *Calm1*, *Actb*, and *Rhoa*) and cell differentiation genes (*Cd44*, *Cebpb*, *Cxcr4* and *Jak2*) in C1 cluster from Fig. 6c across WT+Iso, WT+αPD-1, *Ptk2b* KO+Iso, and *Ptk2b* KO+αPD-1 groups. **h,** The potential intercellular communication of PD-L1 signaling pathway based on the calculation of interactive ligands-receptors between Endo/Fibro/Epi (malignant) cells and immunocytes in the WT+Iso, WT+αPD-1, *Ptk2b* KO+Iso, and *Ptk2b* KO+αPD-1 groups (from left to right). The lines connect to the cell types that express the cognate receptors or ligands. The thickness of line and interaction weight score indicate the number and expression level of ligand-receptor pairs. **i,** Representative images (left) and quantification (right) of mIHC staining of CD8, Granzyme B (GzmB), Perforin, Ki67, and DAPI in PDAC in WT or *Ptk2b* HOM mice that received either Isotype control or αPD-1 treatment. n = 5 (WT+Iso), n = 5 (WT+αPD-1), n = 4 (*Ptk2b* HOM+ Iso), n = 4 (*Ptk2b* HOM+αPD-1) mice. Scale bars, 20 μm. **j**, Survival curves of WT and *Ptk2b* HOM mice in response to Isotype control or αPD-1 treatment. The mice received intraperitoneal administration of either Isotype control or αPD-1 treatment (200 µg) every 4 days once the tumor reached approximately 0.5 cm in diameter. The experiments were concluded when the tumor size reached 1.5 cm or if there was a 20% loss in body weight. n = 10 (WT+Iso), n = 7 (WT+αPD-1), n = 7 (*Ptk2b* HOM+Iso), n = 7 (*Ptk2b* HOM+αPD-1) mice. *P* values in (**i**) were determined by ordinary one-way ANOVA; *P* values in (**j**) were determined by Log-rank (Mantel-Cox) test.

To investigate the impact of PYK2 loss on the differentiation of monocytes *in vivo*, we conducted an integrated and pseudotime analysis of monocytes and macrophages. This analysis identified three distinct differentiation pathways—Pathway 1, Pathway 2, and Pathway 3—which were further divided into five clusters (Fig. 6b and c). To clarify the functions of the five clusters, we conducted enrichment analysis on their highly up-regulated expressed genes (Fig. 6d). Cluster 1, representing the initial state, was enriched with genes associated with monocyte activation and was composed almost entirely of monocyte cells. As pseudotime progressed (Fig. 6d; Supplementary Fig. S21a and b), Cluster 1 differentiated into either Cluster 2 or Cluster 3. Cluster 2 indicated a remodeling state, enriched in genes related to hemopoiesis regulation, connective tissue development, and wound healing. Cluster 3, serving as the intermediate state, branched into either inflammatory cells (Cluster 4), which were enriched in genes linked to the positive regulation of inflammatory responses, or alternatively activated cells (Cluster 5), which showed enrichment in processes such as cell-substrate adhesion and the ERK1/ERK2 cascade, crucial for the polarization of alternatively activated macrophages (44).

Notably, loss of PYK2 in myeloid precursors led to a significant increase of M1-like inflammatory macrophages and reduction of alternatively activated macrophages *in vivo*, suggesting that PYK2 may not only regulate monocyte differentiation but also influence macrophage polarization (Fig. 6e and f). This shift from "pro-tumor" to "anti-tumor" macrophages, induced by PYK2 loss, potentially contributes to the reduced tumor burden in this cohort. Interestingly, there was a significant increase of remodeling macrophages in tumors from the *Ptk2b* HOM mice receiving αPD-1 treatment, indicating that the recruited monocytes/macrophages may play a supportive role in tissue recovery and reconstitution.

To further confirm the influence of mechanical status on the activation of monocytes, we examined genes related to mechanotransduction in the initial monocytes (cluster 1). As expected, a reduction in genes such as *Piezo1*, *Calm1*, *Actb*, and *Rhoa* was observed in the cohort with PYK2 loss (Fig. 6g). This reduction also extended to key cell differentiation-related genes such as *Rela* and *Jak2* (Fig. 6g). Overall, these findings, consistent with previous results, suggest that inhibiting PYK2 disrupts the mechanosensitive pathways affecting monocyte differentiation.

Moreover, the impact of PYK2 loss extends beyond monocyte differentiation, affecting physical environment and other cell types as well. In *Ptk2b* HOM mice, tumor fibrosis and stiffness decreased significantly (Fig. 5c–f). scRNA-seq analysis identified fibroblasts as the primary ECM producers, with the highest “Extracellular Matrix Assembly” and “Collagen Trimer” pathway scores (Supplementary Fig. S22a), while macrophages also substantially contribute to fibrosis (45,46), with elevated “Collagen Trimer” scores (Supplementary Fig. S22b). In these mice, fibroblasts exhibited reduced ECM pathway scores alongside reduced expression of *Col1a1* and *Lox* (Supplementary Fig. S22c). Similarly, macrophages exhibited lower “Collagen Trimer” score (Supplementary Fig. S22c), alongside reduced expression of pro-fibrotic mediators (e.g., *Il1b*, *Pdgfa*, *Ccl2*) in combined monocyte and macrophage population (Supplementary Fig. S22d). Notably, TGF-β signaling, a critical fibrosis driver between fibroblasts and macrophages (47–49), was markedly suppressed (Supplementary Fig. S22e, red arrow). *In vitro* Transwell co-culture experiments further confirmed that *Ptk2b* KO macrophages suppress CAF proliferation and activation, as demonstrated by reduced protein expression of Ki67 and α-SMA (Supplementary Fig. S22f). Collectively, *Ptk2b* deficiency in myeloid cells attenuates fibrosis by directly limiting ECM synthesis and indirectly suppressing fibroblast activation.

This altered stomal landscape ties closely to fibroblast behavior across treatment groups. In the WT+Iso cohort, fibroblasts were highly enriched in pathways related to collagen biosynthesis and chemokine response, indicating myofibroblast and inflammatory phenotypes (50). In contrast, in *Ptk2b* HOM mice, either alone or combined with αPD-1 treatment, fibroblasts displayed lower activation levels in terms of ECM production and inflammatory secretion, with the combo group showing the lowest activation levels (Supplementary Fig. S23a). This reduced fibroblast activity potentially contributed to the decreased stiffness of the TME in the combo-treated mice.

These changes in monocyte differentiation, macrophage function and physical properties finally led to a noticeable increase in cytotoxic T cell infiltration in the *Ptk2b* HOM+αPD-1 cohort. This combined therapy effectively disrupted the PD-L1 signaling pathway between CD8^+^ T cells and other cells (Fig. 6h), leading to a substantial increase in cytotoxic CD8^+^ T cells (Fig. 6i; Supplementary Fig. S24a and b), and a decrease in terminally exhausted CD8^+^ T cells (Supplementary Fig. S24c and d), ultimately aiding in the eradication of tumor cells in the combined targeted cohort.

Finally, the overall survival of mice was monitored. None of the WT+Iso mice survived PDAC as it is a highly lethal cancer. Myeloid PYK2 loss or αPD-1 treatment extended their lifespan but still, most mice were unable to overcome the disease. Notably, the combination of αPD-1 treatment and targeted deletion of PYK2 in myeloid precursors markedly enhanced the overall survival rate (Fig. 6j), indicating the promising therapeutic approach of concurrently targeting myeloid PYK2 and PD-1 in the treatment of PDAC patients.

## DISCUSSION

PDAC is characterized by a dense fibrotic ECM that creates a rigid TME. In response to this stiffened and fibrotic TME, macrophages adopt an alternatively activated phenotype that contributes to immune dysfunction. Our findings, consistent with previous studies (13,51), indicate that the predominant macrophage population in PDAC originates from monocytes and significantly influences disease progression. However, the precise impact of the mechanical properties of the tumor ECM on monocyte differentiation remains insufficiently understood. Our investigation revealed that macrophages located within the fibrotic regions of PDAC exhibit elevated expression of genes associated with responses to mechanical stimulus and regulation of cell differentiation. Through both *in vivo* and *in vitro* experiments, we demonstrated that substantial mechanical cues are essential for driving monocyte differentiation into macrophages, with actin filament-mediated activities playing a particularly crucial role. These findings shed light on the pivotal role of mechanical signals in orchestrating the differentiation of monocytes into macrophages within TME of PDAC.

PYK2, encoded by *PTK2B*, is a non-receptor kinase that belongs to the FAK family, sharing homology with FAK in terms of amino acid sequence and domain structure. Both PYK2 and FAK contain an N-terminal band 4.1/Ezrin/Radixin/Moesin (FERM) domain, a protein tyrosine kinase (PTK) domain, three proline-rich domains, and a C-terminal focal adhesion targeting (FAT) domain (52). While FAK is ubiquitously expressed in most cell types, PYK2 expression varies across different cell types, with notably higher levels in neurons and hematopoietic lineages (53,54). Despite sharing some activation mechanisms, PYK2 possesses distinctive characteristics, such as its high sensitivity to Ca^2+^ (55). In this study, PYK2 was identified as a critical immunomechanical checkpoint that drives the differentiation of monocytes into macrophages.

Our study unveiled that the expression and activation of PYK2 are dependent on mechanical forces, with activation triggered by the mechanosensitive ion channel Piezo1 or integrin β1-mediated adhesion. Notably, mRNA levels of *Trpv4* increase with substrate stiffness in differentiating monocytes (Supplementary Fig. S5), aligning with its stiffness-sensing role in macrophages (56,57). Yet, TRPV4 activation typically relies on upstream mechanosensors, such as PIEZO1 or integrins^58,59^, suggesting it acts primarily as a downstream Ca^2+^ amplifier (58,59). Future studies will investigate whether TRPV4 synergizes with PIEZO1 or integrins to amplify PYK2 activation and facilitate its nuclear translocation.

This mechanical regulation extends to cytoskeletal dynamics, where inhibiting F-actin polymerization with Latrunculin B more potently disrupted PYK2 activation than inhibiting myosin II activity with Blebbistatin. Furthermore, loss of PYK2 in monocytes led to significant disruption in the arrangement of F-actin and microtubules. It has been reported that PYK2 interacts with several receptors, such as integrin and chemokine receptors (60–64), which engage various downstream events including small G protein activation and focal adhesion formation that play a role in actin reorganization (65,66). As a pivotal tyrosine kinase in hematopoietic cells, PYK2 greatly influences the organization of the actin cytoskeleton. Consistent with our findings, osteoclasts lacking PYK2 exhibited a marked reduction in microtubule acetylation and stability (67). These results suggest that PYK2 serves as a mechanical checkpoint in sensing and transducing mechanical forces by coordinating rearrangements in cytoskeletal proteins.

During monocyte differentiation, we observed the nuclear accumulation of PYK2, a phenomenon also reported in neurons, tumor cells, and macrophages (68–70). While the exact mechanism and function of nuclear PYK2 remain elusive, it is thought to modulate gene expression in a cell type-specific manner (69,70). Our research has identified specific gene sites associated with nuclear PYK2, revealing its direct or indirect interaction with multiple promoters that regulate genes crucial for actin cytoskeleton dynamics and NF-κB signaling. *ACTR3*, which encodes ARP3—a core component of the Arp2/3 complex responsible for nucleating branched actin filaments—was identified as a downstream gene regulated by PYK2. Arp2/3 has a critical function in F-actin assembly and cell migration (71,72). Our study has shown that inhibiting the Arp2/3 complex activity hindered monocyte differentiation, highlighting the critical role of F-actin polymerization in this process. Rel/NF-κB transcription factors have been shown to orchestrate survival and differentiation in various hematopoietic lineages (73). Mice deficient in myeloid RelA exhibited reduced inflammation in a nephritis model, while restricting RelA activity suppressed macrophage alternative activation under oncogenic stress (74,75). The loss of PYK2 resulted in a reduction in *RELA* gene expression, which led to impaired monocyte differentiation and altered macrophage polarization. Consequently, nuclear PYK2 facilitates the expression of transcripts involved in F-actin polymerization and cell differentiation, establishing a regulatory network that triggers a positive feedback loop, enhancing cytoskeletal organization and promoting cell differentiation.

In addition to influencing cell differentiation, the inhibition of PYK2 resulted in altered macrophage polarization and decreased cell motility. Macrophages are highly dynamic cells that undergo rapid cellular remodeling during processes such as polarization, migration, and phagocytosis, which are intricately regulated by changes in actin cytoskeleton organization and focal adhesions (76). The deficiency of PYK2 significantly disrupted F-actin assembly, reduced myosin II activation, and decreased ARP3 expression, consequently impairing cell migration and chemotactic responses. Consistent with our findings, the loss of PYK2 hindered the migration of macrophages and cytotoxic T lymphocytes in response to chemokine stimulation (77,78). Previous studies have shown that increased stiffness can shift macrophages from classically activated phenotype to alternatively activated phenotype (79,80). Herein, we demonstrate that PYK2 directed the polarization of macrophages. The deficiency of PYK2 proteins led to a decrease in cell stiffness, promoted the pro-inflammatory phenotype, and suppressed the alternatively activated phenotype of macrophages. These alterations ultimately brought significant changes in the TME of PDAC.

Desmoplasia in PDAC fosters immune evasion by limiting cytotoxic T cell infiltration and impeding drug delivery. The interplay between CAFs and alternatively activated macrophages sustains this fibrotic response (81–85). Modulating macrophage function offers an alternative strategy for anti-fibrosis strategy (86). In *Ptk2b* HOM mice, fibrosis reduction results from diminished macrophage ECM production and disrupted macrophage-CAF crosstalk via TGF-β and other mediators (e.g., IL-1β, PDGF, CCL2). While CAFs are the primary ECM source, PYK2 in macrophages critically regulates their activation, indirectly attenuating CAF function and stromal stiffness.

TAMs also drive T cell dysfunction, contributing to immunosuppression (87,88). *Ptk2b* deletion reprograms macrophages to promote CD8^+^ T cell proliferation and preserve their stem-like properties, enhancing αPD-1 efficacy. Although αPD-1 directly reinvigorates T cells, its success in PDAC hinges on a permissive TME. Myeloid-specific PYK2 loss facilitates this by: (i) reducing fibrosis to improve T cell infiltration and αPD-1 delivery, and (ii) shifting macrophage function to preserve CD8^+^ T cell stemness and support priming. Thus, *Ptk2b* HOM and αPD-1 synergize through a division of labor: *Ptk2b* deletion optimizes the stromal and immune niche, amplifying αPD-1’s ability to unlock T cell anti-tumor potential.

A limitation of this study lies in the use of palpation-based monitoring to assess tumor size during growth. Given the deep abdominal location of orthotopic pancreatic tumors, this method demands highly skilled technicians to ensure accuracy. Its reliability was confirmed by correlating palpation measurements with those obtained at the study endpoint via surgical dissection (Supplementary Fig. S25a-b). Palpation-based measurement is widely employed in PDAC research (89–93) due to its simplicity, frequent applicability, and minimal interference with tumor progression compared to imaging modalities such as ultrasound or micro-CT (94–96). Nonetheless, its subjective nature and challenges in precisely detecting deep-seated tumors introduce potential variability. To enhance precision, future studies could combine biweekly palpation with weekly ultrasound imaging, striking a balance between accurate monitoring and minimal disruption to the TME.

In conclusion, PYK2 acts as an immunomechanical checkpoint in PDAC, orchestrating mechanical activation of monocytes that drives their differentiation and polarization into pro-tumorigenic macrophages within stiff tumor niches. These findings underscore the therapeutic potential of myeloid-specific PYK2 inhibitors, which could disrupt mechanotransduction in monocytes, reduce TAM abundance, and reshape their phenotype. Such strategies hold promise for overcoming the immunosuppressive and fibrotic barriers in PDAC, paving the way for more effective immunotherapies.

## METHODS

### Human PDAC samples

Human adjacent normal tissues and tumor tissues were collected from primary PDAC patients with informed written consent, and under the approval of local medical ethics from West China Hospital Medical Committee, Sichuan University (2021-1190). Freshly resected human PDAC samples were kept in PBS containing protease inhibitor (cOmplete Tablets EDTA-free, *EASYpack*, Roche) on ice, and ready for transport. Detailed information about the donors, including age, sex and diagnosis is reported in Supplementary Table 1.

### Animals

The *Kras^G12D^* (B6/JGpt-*Kras^em1Cin(G12D)^*/Gpt, strain NO. T007054), B6*-p53^R172H^* (B6/JGpt-*Trp53^em1Cin(LSL-R172H)^*/Gpt, strain NO. T007671), B6*-G/R* (B6/JGpt- *H11^em1Cin(CAG-LoxP-ZsGreen-Stop-LoxP-tdTomato)^*/Gpt, strain NO. T006163), *Ptk2b-flox* (B6/JGpt-*Ptk2b^em1Cflox^*/Gpt, strain NO. T012869), *Ptf1a-CreERT2* (B6/JGpt- *Ptf1a^em1Cin(P2A-CreERT2)^*/Gpt, strain NO. T006208), *Lyz2-Cre* mice (B6/JGpt-*Lyz2^em1Cin(iCre)/Gpt^*, strain NO. T003822) were purchased from GemPharmatech Co. Ltd (Nanjing, China). The KPC mice (*Kras^G12D/+^; TP53^R172H/+^; Ptf1a-CreERT2*) were bred using the following strategy: *Kras^G12D^* mice were crossed with B6*-p53^R172H^*mice and pancreas-specific *Ptf1a-CreERT2* mice to yield mice that possessed a conditional p53 mutation and endogenous levels of mutant *Kras^G12D^*expressed by pancreatic tissue. Upon reaching 8-10 weeks of age, the KPC mice underwent a 5-day treatment with Tamoxifen. For experimental purposes, pancreatic tumors were harvested when the KPC mice lost approximately 20% in body weight, and the tumor burden had reached a diameter of about 1.5 cm.

The myeloid tdTomato^+^ mice were obtained through the crossing of *Lyz2-Cre* mice with *B6-G/R* mice. In the offspring, the presence of Cre recombinase led to the deletion of Zsgreen in the mouse genome, turning on the expression of tdTomato with red fluorescence. *Ptk2b* HET (*Ptk2b^fl/+^, Lyz2-Cre*) mice were obtained by crossing *Lyz2-cre* mice with *Ptk2b^fl/fl^* mice, while *Ptk2b* HOM mice (*Ptk2b^fl/fl^, Lyz2-Cre*) were generated by mating *Ptk2b* HET mice. The genotyping primers are listed in Supplementary Table 5. All mice were housed in specific pathogen-free (SPF) conditions at the West China Hospital Animal Center. All mouse experiments were approved and supervised by the Ethics Committee of Animal Experiments of the West China Hospital of Sichuan University for Animal Care and Use (#2020361A).

### Hydrogel synthesis and preparation

Gelatin methacrylate (GelMA) was synthesized as described previously (97). Briefly, 5g gelatin (Sigma) was dissolved into 50 mL Dulbecco’s phosphate buffered saline (DPBS) at 60 °C. 1 mL of methacrylic anhydride was added dropwise to the gelatin solution under stirring at 50 °C and left to react for 2 hours. The reaction was then quenched by adding 100 mL of pre-warmed (40 °C) DPBS, following which the mixture underwent dialysis against distilled water using 12-14 kDa cutoff dialysis tubing for 4 days at 40 °C. Finally, the water was removed through lyophilization, and the GelMA was stored at 4 °C until further use.

Poly (ethylene glycol) diacrylate (PEGDA, 700 molecular weight; Sigma) and GelMA were used to prepare hydrogels. Briefly, PEGDA and GelMA were dissolved in DPBS. PEGDA (4% and 60% w/v) was mixed with an equal volume of GelMA (8% w/v) to generate “soft” and “stiff” hydrogels, respectively. An equal volume of PEGDA (60% w/v) and DPBS were mixed to fabricate “stiff antifouling” hydrogel. After adding the photoinitiator lithium phenyl-2,4,6-trimethylbenzoylphosphinate (LAP, Sigma, US) with a concentration of 0.05 mg/mL, the polymer was crosslinked under UV exposure for 2 min. The gels were hydrated with PBS at 4 °C overnight before use.

### *In vivo* implantation of hydrogel

To simulate the TME of PDAC with various stiffness, stiff or soft hydrogel was prepared and homogenized in PBS (at a volume ratio of 1:2) using the Bullet Blender Homogenizer (Next advance, US) until no visible particles were present. The homogenized hydrogels were combined with 3×10^5 KPC cells suspended Matrigel (Corning, US) at a volume ratio of 2:1. For subcutaneous simulation, 100 μL of the mixture was injected subcutaneously into the left and right back of mice, respectively. The tissues were collected for nanoindentation and mIHC staining three weeks after implantation. For orthotopic simulation, 50 μL of the mixture was orthotopically injected into the head of the pancreas, and the tissues were collected three weeks after implantation.

### Orthotopic model of PDAC and immunotherapy

In order to simulate PDAC within a well-established stiff TME, 1×10^6 KPC cells suspended in Matrigel (Corning, US) were subcutaneously injected into WT mice. Subsequently, after a 2-week interval, the subcutaneous tumors were collected, dissected into pieces of approximately 1mm, and then implanted into the pancreas of WT, *Ptk2b* HET, or *Ptk2b* HOM mice to establish PDAC mouse models. The PDAC tumors were palpated and tumor size was monitored every 2 or 3 days using vernier calipers. The mice were euthanized 35 days after implantation.

To evaluate the efficacy of αPD-1 treatment, 5 × 10^5 KPC cells suspended in Matrigel (Corning, US) were injected into the pancreatic head of WT and *Ptk2b* HOM mice using an insulin syringe with a needle (KRUUSE, 0.5 mL, 0.3×12mm; US). The tumor diameter was assessed using calipers every other day starting from 7 days after injection. Once tumors reached an approximate diameter of 5 mm, mice were randomly divided into two treatment groups based on their tumor size. Each group received either an IgG2a isotype control (RRID: AB_1107769; BE0089, BioXCell, US) or αPD-1 antibody (RRID: AB_10949053; BE0146, BioXCell, US) treatment via intraperitoneal (i.p.) injection every 4 days at a dose of 200 μg, as previously described (98). Similarly, tumor growth was monitored longitudinally *in vivo* every 2-3 days using vernier calipers. For the time-point experiment, the mice were sacrificed after four rounds of injections. For the end-point experiment, the mice were sacrificed when the tumor size reached 1.5 cm or if there was a 20% loss in body weight.

Tumor size was determined through a palpation-based measurement protocol. The tumor, located near the spleen in the left upper quadrant, was gently palpated and immobilized between the thumb and forefinger. The distance between the fingers was measured using digital calipers, with measurements repeated 2-3 times for accuracy (Supplementary Fig. S25a). All assessments were conducted by a single trained technician to ensure consistency and reproducibility.

### Nanoindentation of tumor tissues and cells

Tissues were freshly dissected and kept in ice-cold PBS until measurement within 8 hours. Mechanical testing was conducted using Piuma Nanoindenter (Optics11 life, Amsterdam, The Netherlands). To obtain a flat surface, the tissues were affixed to the bottom of 3.5 cm diameter petri dishes with super glue (Pattex, China). A spherical probe with a radius of 50 µm and a cantilever stiffness of 0.5 N/m was utilized. The applied indentation protocol involved a loading phase of 2 s at an indentation depth of 10000 nm, which was held for one second, followed by a 2 s unloading phase.

To determine elastic properties of cells, BMDMs or THP-1 cells were cultured in 3.5 cm diameter petri dishes. Prior to measurement, the medium was replaced with PBS. The indentation protocol consisted of a loading phase of 2 s at an indentation depth of 3000 nm, held for one second, and then an unloading phase of 2 s. All individual indentation values were calculated using Piuma Data Viewer version 2.2 (Piuma; Optics11, Amsterdam, The Netherlands).

### Histology staining

Human PDAC and adjacent normal tissues, as well as mouse tumor tissues, were preserved in 4% paraformaldehyde and embedded in paraffin following standard protocols. Human PDAC and adjacent normal tissues were also mounted in OCT for frozen sectioning. H&E and Masson’s trichrome staining were performed using standard protocols. Images were analyzed using HALO software to quantify the fibrotic area across the entire slides.

### Multiplex immunohistochemistry staining

The human pancreatic tissue microarray (HPanA180Su03; Shanghai Outdo Biotech Company, China) was utilized for mIHC staining. Mouse subcutaneous tumor or orthotopic PDAC tumor tissues underwent mIHC staining as well. The staining for mIHC was carried out with the Opal Polaris 7 color IHC Detection Kit (Akoya Biosciences, USA) according to the manufacturer’s protocol. Briefly, tissue slides were deparaffinized with xylene, rehydrated through a graded ethanol series to water.

A microwave treatment in the AR6 buffer (Akoya Biosciences) was applied for antigen retrieval. For the removal of endogenous peroxidase, the slides were incubated at room temperature for 10 min with H_2_O_2_. After blocking for 10 minutes at room temperature, the tissue was incubated with the primary antibody for 1 hour. The primary antibodies used are listed in Supplementary Table 6. A secondary HRP antibody (Akoya Biosciences) was added and incubated at room temperature for 10 min. Signal amplification was achieved using TSA working solution diluted at 1:150 in 1 × amplification diluent (Akoya Biosciences) and incubated at room temperature for 10 min. The multispectral imaging was collected by Vectra Polaris™ Automated Quantitative Pathology Imaging System (Akoya Biosciences) at 20× magnification and analyzed by HALO software (Indica Labs).

### Sequential IHC, image processing and analysis

Sequential IHC was performed as per a previously established protocol (99). Briefly, the formalin-fixed paraffin-embedded human PDAC tissue sections were deparaffinized by Xylene and gradient alcohol solutions and stained with hematoxylin (S3301, Dako, US) for 1 min. The whole slide was scanned using Pannoramic MIDI (3D HISTECH). The sections were subsequently blocked with H_2_O_2_ for 30 minutes, followed by heat-mediated antigen retrieval in a pH6.0 citrate solution (HK080-9K, BioGenex, US). A total of 9 staining, scanning, and chromogenic stripping cycles were executed for the visualization of 9 markers. Details regarding the primary antibodies, incubation process, and AEC reaction times for staining and color detection of sections can be found in Supplementary Table 7.

For sequential IHC image processing and analysis, the tumor epithelial localization was identified by the presence of CK19 markers on individual slides. The same processing was applied to other markers as well. At least three regions of interest (ROIs) were chosen within each sample for analysis, with each containing 5000 × 5000 pixels. Visualization was performed by converting coregistered images into individually pseudocolored single-marker staining images through ImageJ version 1.48 (imageJ, US) and Aperio ImageScope version 12.4.6 (Leica Biosystem, Germany). To obtain single-cell segmentation and quantify the signal, the image was processed using CellProfiler version 2.1.1 pipeline (US). Subsequently, the image underwent quantitative analysis using FCS Express and Image Cytometry software version 7.10.0007 (De Novo Software, US).

### Cell culture

The THP-1 cell line (RRID: CVCL_0006) was purchased from Cell Bank/Stem Cell Bank, Chinese Academy of Sciences (Shanghai, China) in June 2022. KPC cells were kindly offered by Dr. Keyu Li (West China Hospital, Sichuan University, China). THP-1 cells and KPC cells were cultured with RPMI1640 medium (Gibco, US) supplemented with 10% FBS (ZETA LIFE, US), and 1% penicillin-streptomycin (Gibco, US). All cells were cultured in a humidified 5 % CO2 incubator at 37 °C. THP-1 cells were routinely tested for Mycoplasma using PCR-based detection (HUABIO, China), while KPC cells for *in vivo* studies were confirmed Mycoplasma-free before implantation. All cells were used within 15 passages.

### Isolation and differentiation of mouse bone marrow cells into macrophages

Primary mouse BM cells were isolated by flushing femurs and tibias from mice. After erythrocyte lysis with ACK buffer (Gibco, US) for 5 min at RT, cells were washed twice with PBS and passed through a filter of 70 μm. BM cells were cultured in RPMI1640 (Gibco, US) containing 10% FBS (ZETA LIFE, US), 1% penicillin-streptomycin (Gibco, US), and 20 ng/mL macrophage colony-stimulating factor (M-CSF; PeproTech, US) on the plastic petri dish for 6 days to induce differentiation into macrophages. The fresh medium was added 3 days after culture.

### Western blot

Cells were homogenized in RIPA buffer (Thermo Fisher Scientific, US) supplemented with cOmplete Protease Inhibitor Cocktail (Roche, US) and PhosSTOP (Roche, US). The samples were vortexed, incubated on ice for 15 min, and centrifuged at 12000 rpm for 15 min at 4 °C. BCA assay (Life-iLab, China) was employed to measure the protein concentration. Equal amounts of proteins were separated on SDS-PAGE, transferred to 0.2 µm NC membranes (Cytiva, US), and blocked with 5 % BSA in Tris-buffered saline containing 0.1 % Tween-20 (TBS-T) for 1 hour at RT. The membranes were incubated overnight at 4 °C with primary antibodies including antibody to PYK2 (RRID: AB_777566; ab32571, Abcam, UK), antibody to p-PYK2 (RRID: AB_2173988; ab4800, Abcam, UK), antibody to FAK (RRID: AB_2269034; #3285, CST, US), or antibody to GAPDH (RRID: AB_2630358; ab181602, Abcam, UK) antibodies. After three washes with TBS-T, the membranes were incubated with HRP-conjugated secondary antibodies (anti-rabbit, Invitrogen, US) for 1 hour. The protein bands were visualized using an ECL detection kit (UElandy, China) on the Tanon 4600 imaging system (Tanon, China). The full blot images and quantification are presented in Supplementary Fig. S26a-l.

### Individual cell-tracking assay and data analysis

BMDMs were seeded at a density of 1×10^5 cells per well on 6-well plates and allowed to grow overnight. The culture medium was then replaced with a conditioned medium obtained from KPC cells. The cells were immediately placed under a phase-contrast microscope (Leica, Germany) inside a CO_2_ microscope cage incubator set at 37 °C and 5% CO_2_. The time-lapse imaging was captured at 10 min intervals for a total duration of 14 hours. Cell tracking analysis was performed using the “Manual tracking” plugin (Fabrice Cordelières, Institut Curie, France) in ImageJ version 2.9.0 (ImageJ, US). Approximately 30-40 cells were manually tracked, and the parameters such as directionality, speed and the distance traveled by individual cells were quantified using the “Chemotaxis and Migration Tool” plugin (ibidi GmbH, Germany).

### siRNA transfection

The siRNA transfection experiment utilized negative control siRNA (si-NC), as well as specific siRNAs targeting *PTK2B* (si-PTK2B), and *PIEZO1* (si-PIEZO1). These siRNAs were obtained from RiboBio (Guangzhou, China), and their sequences are provided in Supplementary Table 5. THP-1 cells were seeded in 24-well plates at a concentration of 2.5 × 10^5 cells per well. Transfection was performed using Invitrogen™ Lipofamine RNAiMAX Reagent (Thermo Fisher Scientific, US) with siRNAs (50 nM), following the manufacturer’s instructions. After 24 hours of transfection, the medium was replaced with PMA-containing medium (50 ng/ml) to induce monocyte differentiation into macrophages.

### Immunofluorescence staining of cells

The cells were fixed in 4% paraformaldehyde for 15 min at RT. After blocking in a solution of 1× PBS / 5% BSA / 0.3% Triton™ X-100, the cells were incubated with primary antibodies diluted in the blocking buffer overnight at 4 °C. Following three washes with PBS, the cells were incubated with secondary antibodies (RRID: AB_2896347; A48283, Invitrogen, US), phalloidin (ab176759, Abcam, UK), and DAPI (AbMole, US) diluted in the blocking buffer for 1 hour at room temperature while being protected from light. The cells were rinsed three times with PBS and observed using a confocal laser scanning microscope (Leica, Germany). The primary antibodies used are listed in Supplementary Table 6.

### RNA isolation and qRT-PCR

Total RNA was extracted using RNAsimple Total RNA Kit (Tiangen, China) according to the manufacturer’s instructions. The first-strand cDNA synthesis was performed using the HiScript III RT SuperMix (Vazyme Biotech, China). Quantitative real-time RT-PCR was conducted on a LightCycler 96 Instrument (Roche, US) using the Taq Pro Universal SYBR qPCR Master Mix (Vazyme Biotech, China). To verify PCR product specificity and avoid false-positive signals, a melting curve analysis was performed. Relative mRNA levels were normalized to the housekeeping gene GAPDH and quantified using the relative quantification 2^ (-ΔΔCt) method. The primer sequences can be found in Supplementary Table 5.

### Co-culture system of T cells and monocytes/macrophages

CD8^+^ T cells were isolated from the spleens of WT mice following the manufacturer’s instructions (MojoSort™ Mouse CD8 T Cell Isolation Kit). Briefly, the spleen was placed on a sterile 70 µm cell strainer mesh in a 50 mL tube filled with ice-cold RPMI 1640 medium and then finely mashed using the plunger of a 5 mL syringe. After washing with PBS, red blood cells were lysed by ACK buffer (Gibco, US) for 5 minutes at room temperature. The cells were then resuspended in MojoSort buffer, and a biotin-antibody cocktail was added and incubated on ice for 15 min. Streptavidin nanobeads were introduced and incubated on ice for an additional 15 min. Subsequently, the MojoSort buffer was added, and using a magnet, the CD8^+^ T cells that were not bound to the beads were separated.

The isolated CD8^+^ T cells were suspended in T cell culture medium [1640 RPMI Medium (Gibco, US) supplemented with 10% FBS (ZETA LIFE, US), 1% penicillin-streptomycin (Gibco, US), MEM, 1 × non-essential amino acids (Gibco, US), 1 mM sodium pyruvate (Sigma, US), 0.02 M HEPES (Gibco, US), 0.05 mM β-mercaptoethanol (Sigma, US), and 10 ng/mL IL-2 (PeproTech, US)]. The T cells were seeded into a 24-well plate at a density of 2 × 10^5 cells/well and activated with CD3-specific (RRID: AB_1107634; BE0001, BioXCell, US) and CD28-specific antibodies (RRID: AB_1107628; BE0015, BioXCell, US) for 24 hours. Subsequently, BM cells were extracted from either WT or *Ptk2b* KO mice following the previously described methods. A total of 4 × 10^4 WT or *Ptk2b* KO BM cells were added to the T cells along with M-CSF (20 ng/mL; PeproTech, US). After a 7-day culture period, the T cells were harvested, and stained with antibodies for flow cytometry analysis (Supplementary Table 8; Cytek Aurora, Cytek Biosciences, US).

### Co-culture system of CAFs and monocytes/macrophages

CAFs were isolated from the pancreas of KPC mice 2 weeks post-tamoxifen induction, following a previously described protocol (100). Pancreatic tissue was washed with PBS, minced into 1-2 mm² fragments, and digested in RPMI-1640 medium supplemented with 3 mg/mL collagenase A (Roche, USA) and 1 mg/mL DNase I (Roche, USA) for 15 minutes at 37°C with gentle agitation. Red blood cells were lysed using RBC Lysis Buffer (eBioscience, USA) for 10 minutes at room temperature. The resulting cell suspension was filtered through a 70 μm cell strainer (Falcon, Corning) and resuspended in RPMI-1640 medium supplemented with 10% fetal bovine serum (FBS). CAFs were identified by IF staining with antibodies against α-SMA (RRID: AB_2734735; #19245, CST) and fibronectin (RRID: AB_2941028; ab268020, Abcam).

For the Transwell co-culture system, BM cells were seeded in Transwell inserts (0.4 μm pore size; Labselect, China), and CAFs were plated in the culture wells of 12-well plates at a 2:1 ratio (BM cells:CAFs) to establish non-contact co-culture. M-CSF (20 ng/mL; PeproTech, US) was added to the BM cells to induce monocyte differentiation into macrophages. After 3 days of co-culture, CAFs were fixed with 4% paraformaldehyde, permeabilized, and stained with antibodies against Ki67 (RRID: AB_302459; ab16667, Abcam) and α-SMA (RRID: AB_2734735; #19245, CST) for IF analysis.

### Flow cytometry

Mouse BM derived cells were stained with Fixable Viability Stain 700 (BD Biosciences, US), followed by blocking with mouse CD16/32-specific antibody (RRID: AB_1574975; Biolegend, US). The fluorescein-conjugated anti-mouse antibodies (Supplementary Table 8) were used for subsequent staining.

To generate a single cell suspension from murine tissues, tumor tissues were chopped and digested by shaking in digestion buffer [3mg/ml collagenase A (Roche, US) and 1 mg/ml DNase I (Roche, US) in RPMI1640 medium] for 30 min. The red blood cells were lysed for 10 min using RBC lysis buffer (eBioscience, US). The cells were then filtered through a 70 μm strainer and resuspended in an appropriate buffer for downstream application. DLN was placed on a sterile 70 µm cell strainer mesh in a 50 mL tube containing ice-cold RPMI 1640 medium and mashed into very fine parts using the plunger of a 5 ml syringe. Blood was collected from the orbital sinus using a capillary tube and red blood cells were lysed for 10 min using RBC lysis buffer (eBioscience, US). For flow cytometry, two million cells from blood, DLN or tumor were stained with Fixable Viability Stain 700 (BD Biosciences, US). After blocking with mouse CD16/32-specific antibody (RRID: AB_1574975; Biolegend, US), the cells were incubated with surface antibodies at 4 °C for 40 min in the dark. For staining intracellular and intranuclear proteins, cells were fixed and permeabilized using Ture-Nuclear Transcription Factor Buffer Set (Biolegend, US), and incubated with fluorescein conjugated anti-mouse antibodies (Supplementary Table 8). Flow cytometry was performed on BD FACSymphony™ A5 Cell Analyzer (BD Biosciences, US) and analyzed using FlowJo (version 10; BD Biosciences, US).

### Ca^2+^ measurement

THP-1 cells were seeded into a 24-well plate at a density of 1 × 10^5 cells/well and primed with PMA for 48 hours. The cells were washed three times with DPBS, and incubated with 5 μM of Fluo-4 AM (Beyotime Biotechnology, China) and 0.1% pluronic F-127 (Beyotime Biotechnology, China) in Hank’s Balanced Salt Solution (HBSS; Biosharp, China) for 30 min. Following this, the Fluo-4 AM-loaded cells were rinsed three times with Ca2^+^-free HBSS (Biosharp, China) and incubated at 37°C for 40 minutes. The fluorescence units were captured using a laser scanning microscope (Leica, Germany). After initially incubating for 1 min, Yoda1 (300 nM) was introduced to the cells, and the change in fluorescence was tracked for 10 min.

### Luciferase assay

A DNA oligonucleotide containing SP-1 and AP-1 binding motifs from the RELA promoter (Supplementary Fig. S15) was cloned into the pGL4.0 plasmid. 293T cells were seeded in 24-well plates and incubated overnight. Cells were then transfected with the pGL4.0 firefly luciferase reporter plasmid and the pRL Renilla luciferase control plasmid (Promega), along with either an empty vector or a PYK2 overexpression plasmid. Transfections were performed using (Lipofectamine LTX, Invitrogen) according to the manufacturer’s protocol. After 24 hours of culture, cells were lysed and harvested for luciferase activity measurement using the Dual-Luciferase Reporter Assay System (#E1910, Promega) per the manufacturer’s instructions.

### DNA pull-down assay

To investigate PYK2’ binding to SP-1 and AP-1 motifs, a DNA pull-down assay was performed as previously described (101,102). A 5’-biotinylated oligonucleotide encompassing the SP-1 and AP-1 binding sites from the RELA promoter (Supplementary Fig. S15) was conjugated to streptavidin-coated magnetic beads (Dynabeads™ M-280, Invitrogen). Nuclear extracts from PMA-treated (100 ng/mL; 48h) THP-1 cells and PYK2 overexpressed 293T cells were prepared using a nuclear extraction kit (Beyotime Biotechnology, China) and incubated with the DNA-conjugated beads for 2 hours at room temperature with constant rotation. Bound proteins were eluted in sample loading buffer, denatured at 95°C for 5 minutes, and resolved by SDS-PAGE. Western blot analysis was conducted using antibodies against PYK2 (ab226798, Abcam), Fra2 (AP-1 subunit; RRID: AB_2722526; #19967, CST), and SP-1 (RRID: AB_3073501; ab227383, Abcam). Binding specificity was validated by a competition assay using a 10-fold molar excess of non-biotinylated oligonucleotides.

### Inhibitors

Inhibitors in the cell culture were used at the following concentrations: 1 µM VS-4718 to inhibit PYK2 (Selleckchem); 5 µM Latrunculin B to inhibit F-actin (Abcam); 10 µM Blebbistatin to inhibit myosin II (MCE); 50, 100 or 200 µM CK-666 to inhibit Arp2/3 (MCE); and 5 µg/ml monoclonal integrin-β1 antibody (RRID: AB_775726; Abcam, ab30394). Vehicle-alone controls for these inhibitors were as follows: DMSO for VS-4718, Latrunculin B, Blebbistatin, CK-666; and IgG nonspecific antibody (Beyotime, A7028) for integrin-β1 antibody. VS-4718, Latrunculin B, Blebbistatin and CK-666 were added to the culture medium directly. Integrin-β1 antibody was incubated with THP-1 cells at 37 °C for 30 min and then primed with PMA (Sigma, 50 ng/ml) for differentiation.

### Generation and processing of scRNA-seq data

#### ScRNA-seq library preparation and sequencing

Mouse tumor samples were collected and dissociated as described above. The scRNA-seq libraries were created using the DNBelab C4 system (BGI-Research, China). Briefly, the single-cell suspension of tumor tissues, mixed with functionalized beads and lysis buffer, was added into a pressure-driven microfluidic device to generate emulsion droplets. The single cell in the droplet was lysed, and mRNA transcripts were released and captured by barcoded beads. Subsequently, the mRNA capture beads were collected, followed by reverse transcription, PCR amplification, and purification. The library’s concentration was measured by Qubit (ThermoFisher, US). The library was sequenced on an MGIseq-2000 platform (BGI-Research, China).

#### Data processing

The raw FASTQ files were processed using the DNBelab_C4scRNA (v1.0.1) software (https://github.com/MGI-tech-bioinformatics/DNBelab_C_Series_scRNA-analysissoftware) and were converted to a format compatible with cellranger (103). Subsequently, the sequencing data were analyzed with STAR software (v2.5.3) (104) using default parameters for mapping to the mm10 reference genome. The aligned reads were then screened to identify valid cell barcodes and unique molecular identifiers (UMIs), facilitating the generation of a gene-cell matrix for subsequent analysis.

#### Quality control of scRNA-seq data

The gene-cell matrix was loaded using the Read10X function of the R package Seurat (v4.1.1) (105). Cells were filtered based on four criteria, which included the number of UMIs, the number of genes, the percentage of mitochondrial (Mt) genes, and the percentage of ribosomal proteins large/small (Rpl/s) genes. The percentage of Mt/Rpl/s genes was calculated using the PercentageFeatureSet function of the R package Seurat. Cells were excluded if they had UMIs less than 900 or greater than 60000, if the number of genes was less than 600 or greater than 8000 if the Mt gene percentage exceeded 10%, or if the Rpl/s percentage was above 25% (Supplementary Fig. S20). Following quality control, 10,737 single cells from WT+Iso experiments, 12,514 single cells from WT+αPD-1 experiments, 13,770 single cells from *Ptk2b* KO+Iso experiments, and 10,103 single cells from *Ptk2b* KO+αPD-1 experiments were included for downstream analysis.

#### Multiple scRNA-seq data normalization and integration

To address batch effects, the Seurat package was used to combine multiple distinct scRNA-seq datasets into a unified and unbatched dataset. Each sample was first normalized using the NormalizeData function, followed by the identification of 2,000 features with high cell-to-cell variation using the FindVariableGenes function. Anchors between individual datasets were identified with the FindIntegrationAnchors function, and these anchors were input into the IntegrateData function to create a batch-corrected expression matrix of all cells, enabling the integration and joint analysis of cells from different datasets.

#### Dimensionality reduction and cell type annotation of scRNA-seq

After quality control and batch effect removal, dimensionality reduction and unsupervised clustering were performed using the R package Seurat (v4.1.1). The integrated dataset was then normalized for principal component analysis, leveraging the first 20 principal components to construct a shared nearest neighbor (SNN) network. The Louvain algorithm was utilized for cell cluster identification, with resolution settings adjusted based on the distinctive characteristics of each dataset. Visualization of these clusters in two-dimensional space was achieved using either UMAP or t-SNE, implemented through the RunUMAP or RunTSNE functions, respectively. Markers for each cluster were identified using the FindAllMarkers function in Seurat with the parameters (min.pct = 0.1, logfc.threshold = 1, only.pos = TRUE). To categorize each cluster into a specific cell type, we selected well-known markers of immune cells, epithelial cells, endothelial cells, and fibroblasts. The cell types were annotated using a dot plot.

#### Single-cell level differential expression analysis

A differential gene expression analysis was conducted using the FindMarkers function from the Seurat (v4.1.1) R package. Genes were selected for further analysis only if they had a Bonferroni-adjusted *P* value below 0.05 and an absolute log2 (fold change) exceeding 0.1. Significance was assessed based on the nonparametric Wilcoxon rank-sum test.

#### Monocle2 pseudotime trajectory inference

Cell differentiation for monocytes and macrophages was inferred using a reversed graph embedding method implemented through the R package Monocle2 (v2.16.0) (106) with default parameters as recommended by the developers. The raw gene-cell matrix from each cell type was first extracted using the GetAssayData function of Seurat into Monocle to construct a CellDataSet employing the newCellDataSet function in Monocle2 (v2.16.0). Subsequently, the expression matrix was normalized utilizing the estimateSizeFactors and estimateDispersions functions in Monocle2 (v2.16.0). The highly variable genes recognized by the FindVariableFeatures function in Seurat (v4.1.1) were employed to order cells in Monocle. This ordering was then utilized to generate the monocyte differentiation to macrophage DDRTree trajectory plot through the setOrderingFilter and reduceDimension functions in Monocle2 (v2.16.0).

#### Cell-cell communication network construction

We utilized the R package in CellChat (v2.1.2) to infer and visualize potential intercellular communication networks (107). The CellChat database contained extensive information on cell-cell communication interactions in murine, including the three categories of Secreted Signaling, ECM-Receptor, and Cell-Cell Contact, with a total of 120 pathways. The normalized expression matrix calculated by Seurat was imported, and a CellChat object was created with the createCellChat function. The computeCommunProb and computeCommunProbPathway functions were employed to calculate ligand-receptor interaction probabilities at both signaling gene and pathway levels. The aggregateNet function was then used to compute the aggregated cell-cell communication network. To visualize significant signaling pathways, circle plots were generated using the netVisual_individual function.

### Generation and processing of spatial transcriptomic data

#### Spatial transcriptomics library preparation and sequencing

Adjacent normal tissues and PDAC tissues were immediately cut into tissue blocks approximately 6.5 mm × 6.5 mm in size embedded in OCT media (Tissue-Tek, US) and frozen in a dry ice slurry. A 10 μm frozen tissue section was cut and mounted onto the ST slides (10× Genomics, US). The tissue was fixed in 4% formaldehyde and stained with hematoxylin-eosin reagents. After taking brightfield images, the tissue was permeabilized, and the RNA was isolated and captured by using spatially barcoded mRNA-binding oligonucleotides on a Visium Spatial platform (10× Genomics, US). The cDNA libraries were prepared and sequenced on an MGIseq-2000 (BGI-Research, China).

#### Data processing

The sequencing data were demultiplexed and mapped to the reference genome hg38 using the SpaceRanger software (v1.2.0) (103) with default parameters. Gene expression was quantified based on the unique molecular identifier (UMI). Count matrices were loaded into Seurat (v4.1.1) for all subsequent data filtering, normalization, and visualization. For quality control, we removed low-quality spots whose number of UMI < 200 or the number of genes < 300. Following quality control, 4823 spots from adjacent normal_1, 4814 spots from adjacent normal_2, 4799 spots from PDAC_1, 4873 spots from PDAC_2, 4760 spots from PDAC_3, and 4599 spots from PDAC_4 were included for downstream analysis. Data normalization was performed on independent tissue sections using the variance-stabilizing transformation method implemented in the SCTransform function in Seurat.

#### Spatial deconvolution

To observe the spatial distribution of immune cells (including T cells, B cells, DCs, monocytes, and macrophages), epithelial cells, endothelial cells, and fibroblasts in adjacent normal tissues and pancreatic ductal adenocarcinoma, RCTD (v2.2.1) (108) was used to deconvolute the transcriptome of each spot into the likely constituent cell types. First, the Reference function was utilized to construct a reference object, and the SpatialRNA function was used to construct a spatial RNA object for deconvolution. Subsequently, the create.RCTD function was employed to create an RCTD object from the reference object and the spatial RNA object. Finally, the run.RCTD function was used to obtain the deconvolution results with the parameter doublet_mode set to ‘full’. The deconvoluted ratios of individual cell types were visualized using Seurat’s (v4.1.1) SpatialFeaturePlot function.

### Generation and processing of bulk RNA-seq data

#### Bulk RNA-seq library preparation and sequencing

Total RNA was isolated from mouse BM-derived cells using TRIzol Reagent following the manufacturer’s instructions (Invitrogen, US). The RNA quality was assessed using Agilent 2100 (Agilent, US). The cDNA libraries were prepared and sequenced on an MGIseq-2000 platform (BGI-Research, China).

#### Processing of bulk RNA-seq data

Initially, the Fastq files underwent a quality control step using FastQC (v0.12.1). The raw reads were then trimmed with Trimmomatic (v0.39) to obtain cleaned reads. The cleaned reads were subsequently mapped to mm10 genomes using STAR (v2.5.3) with default parameters. Quantification of the cleaned reads was performed using GenomicAlignments (v1.24.0) (109). Genes with raw total expression levels below 10 across all samples were removed. The quality-controlled gene expression matrix was then normalized using the DESeq2 (v1.24.0) package (110).

### Generation and processing of CUT&Tag-seq data

#### CUT&Tag library preparation and sequencing

DNA binding sites of PYK2 were profiled using CUT&Tag assays with the Hyperactive Universal CUT&Tag Assay Kit for Illumina (Vazyme Biotech, #TD904, China). In brief, 1 × 10^5 THP-1 cells, mouse monocytes, or BMDMs were harvested and washed. Subsequently, they were bound to ConA beads for 10 minutes at room temperature. The cells were then incubated overnight at 4 °C with a PYK2 antibody (Abcam, ab226798). A secondary antibody was applied and incubated for 1 hour at room temperature. After three washes with Dig-wash buffer, the cells were treated with 0.04 μM pA/G-Tnp Pro for 1 hour at room temperature. Following another wash with Dig-300 buffer, the cells were resuspended in TTBL buffer and incubated at 37 ℃ for 1 hour. SDS and DNA Spike-in (1 pg) were added and incubated at 55 °C for 10 min. The nucleic acid was released into the liquid by vigorous pipetting. The liquid was collected and mixed with DNA Extract Beads Pro, incubated for 20 min at room temperature, washed, and resuspended in double-distilled water (ddH_2_O). The library was generated by PCR amplification (11 cycles) and purified using VAHTS DNA Clean Beads (Vazyme Biotech, #N411, China). The library concentration was quantified using real-time PCR. The library quality was assessed using Fragment Analyzer (Agilent, US). Finally, the libraries were sequenced on a Novaseq PE150 platform (Illumina, US).

#### Processing of CUT&Tag-seq data

CUT&Tag data processing and analysis were performed following a standard protocol (111,112). Initial quality checks on the sequencing reads were conducted using FastQC (v0.12.1) (https://github.com/s-andrews/FastQC) and MultiQC (v1.12) (113). Subsequently, raw reads were processed through stringent bioinformatics pipelines, which included trimming and filtering for quality using Trimmomatic (v.0.39) (114). Following this, paired-end reads were aligned using Bowtie2 (v2.5.1) (115) against hg38 genomes downloaded from Ensembl (https://asia.ensembl.org/index.html) and spike-in sequences from *E. coli* genomes with the parameters --local, --very-sensitive-local, --no-mixed, --no-discordant, --phred33, -I 10, -X 700, -p 15. Aligned reads were sorted, and non-uniquely mapping reads were annotated using Picard (v3.0.0) (http://broadinstitute.github.io/picard/). PCR duplicates and reads with low mapping quality (MAPQ < 10) were filtered using SAMtools (v1.18) (116). Peak calling was then identified using MACS2 (v2.2.7.1) (117), utilizing the following parameters with pre-processed BAM files: -t pathway of the treatment cohort BAM files, -c pathway of the control cohort BAM files, -f BAMPE, -g hs, -B, -p 0.05. Motif analysis was performed using HOMER (v5.1) with default parameters (43). Additionally, motif searching was conducted using the MEME (v1.13.1) R package, specifically the runFimo function, with a threshold parameter (thresh = 1e-3) (118). Peaks were annotated using the R package ChIPseeker (v1.24.0) (119), and visualizations were generated using deepTools (v3.5.4) (120) and pyGenomeTracks (v3.8) (121). Additionally, the enrichment of gene ontology terms for peak sets was conducted using ClusterProfiler (v4.9.0.002) (122).

### Gene set enrichment and gene ontology analysis

To conduct Gene Ontology (GO) and Kyoto Encyclopedia of Genes and Genomes (KEGG) analyses, we utilized the enrichGO and enrichKEGG functions within the R package ClusterProfiler (v4.9.0.002). The org.Hs.eg.db (v3.18.0) and org.Mm.eg.db (v3.18.0) packages were employed for these analyses. The results were visualized using the dotplot function from the enrichplot (v1.20.0) package (https://github.com/YuLab-SMU/enrichplot). For Gene Set Enrichment Analysis (GSEA), genes were initially ranked based on their log2 (fold change), and enrichment analysis was performed using the gseGO function with the org.Mm.eg.db (v3.18.0) package. Visualization of the results was achieved using the GseaVis (v0.0.5, https://github.com/junjunlab/GseaVis) package.

### Calculating gene set signature scores

To observe the expression of mechanosensitive and differentiation-related pathways, gene sets derived from the GO Biological Process ontology were used from the MSigDB (v2023.2) (123) and were exported using the R package GSEABase (v1.50.1). For scRNA-seq data, gene set signature scores were calculated for each cell using the Seurat function AddModuleScore. For spatial data, gene set signature scores for each spot were computed using the runPAGEEnrich function from the R package Giotto (v1.1.2) (124). To analyze bulk RNA-seq data, we calculated gene set signature scores for each sample using the gsva function from the R package GSVA (v1.36.3) (125), employing the ssgsea method.

### Public scRNA-seq data analysis

To perform deconvolution analysis on spatial transcriptomics data, we downloaded a previously published reproducible PDAC scRNA-seq atlas (29). To investigate the process of monocytes infiltrating tumor tissue from the vasculature and differentiating into macrophages in PDAC patients, we obtained single-cell sequencing data of blood and tumor tissue from the same PDAC patient from the National Genomics Data Center (NGDC, https://ngdc.cncb.ac.cn/) under GSA-human: HRA003672 (32).

### Statistics and reproducibility

All statistical analyses were conducted using R (v4.2.1) and Prism10 (GraphPad). Sample sizes were not predetermined based on statistical power calculations. Mice were allocated to experiments randomly and processed in arbitrary order. Various statistical tests including two-tailed Student’s t-test, one-way ANOVA test, two-way ANOVA test, nonparametric Wilcoxon rank-sum test, or chi-square test were applied as specified in the figure legends. The precise number of biological or experimental replicates (n) can be found in the figure legends, and the exact *P* value was presented in the graphed data. A *P* or adjusted *P* value < 0.05 was considered statistically significant. All the experiments were carried out at least two or three times, and the key findings were reliably reproduced.

## Data and Code Availability

All the raw mouse sequencing data have been deposited in the CNCB (https://ngdc.cncb.ac.cn/). Spatial transcriptomics data under accession number: PRJCA029235. ScRNA-seq data under accession number: PRJCA029236. Bulk RNA-seq data under accession number: PRJCA029237. CUT&Tag data under accession number: PRJCA029614. All sequencing data are publicly available as of the date of publication. The original code used for data analysis is available at https://codeocean.com/capsule/4495066/tree. Additional information required to reanalyze the data reported in this study can be obtained from the corresponding author upon request.

## Authors’ Contributions

H.J. and Q.W. designed the research; B.S.D. and Y.H.L. supervised in vivo experimental designs. B.S.D. and H.B.S. supervised scRNA-seq analyses; Q.W. supervised in vitro ECM model designs. W.Y.X., P.W. and H.J. wrote and edited the paper; W.Y.X., J.Y. and N.W.K. collected clinical samples; W.Y.X., X.Y., and Q.X.Y. analyzed clinical sample data and scRNA-seq; P.W., P.P.M and Lang C. performed in vitro assays; W.Y.X., X.Y., X.Q.F. and X.J.W performed in vivo mouse models; X.Y., H.K., and P.W. performed mIHC and analyzed data; L.C. helped with bioinformatic analysis. Y.W. proofread the paper.

## Acknowledgments

We would like to thank all members of the Hong Jiang laboratory (Sichuan University) for their insightful comments regarding this work. We thank Ting Cao (Sichuan University) for her assistance in conducting flow cytometry. We acknowledge the Core Facilities of West China Hospital, Sichuan University. This work was supported by the National Natural Science Foundation of China (82073158, to Hong Jiang; 82072615 and 82273274, to Yun-Hua Liu) and 1.3.5 project for disciplines of excellence from West China Hospital of Sichuan University (ZYYC24005, to Hong Jiang).

