## Supplemental Figures for "An immunomechanical checkpoint PYK2 governs monocyte-to-macrophage differentiation in pancreatic cancer"

**Supplementary Fig. S1**

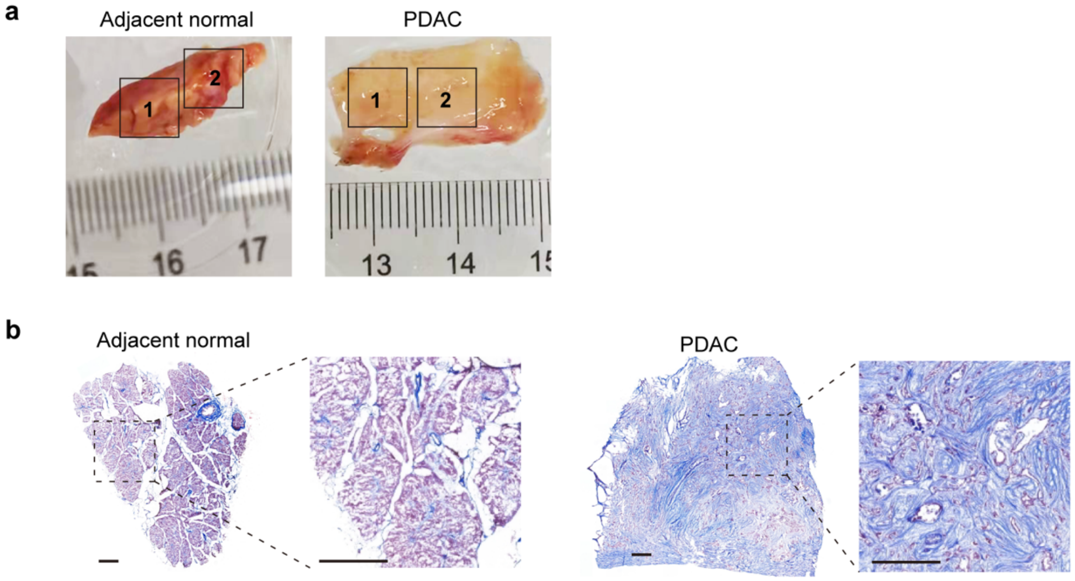


**Supplementary Fig. S1 Processing of human PDAC samples.**

**a,** Representative images of human adjacent normal and PDAC tissues. Two portions were cut from the fresh tissues. Portion No. 1 was processed for nanoindentation analysis, followed by Masson's trichrome staining and mIHC staining. Portion No. 2 was embedded in OCT, frozen and sectioned for ST analysis.

**b,** Representative images of Masson's trichrome staining of human adjacent normal and PDAC tissues. Scale bars, 100 μm.

Supplementary Fig. S2

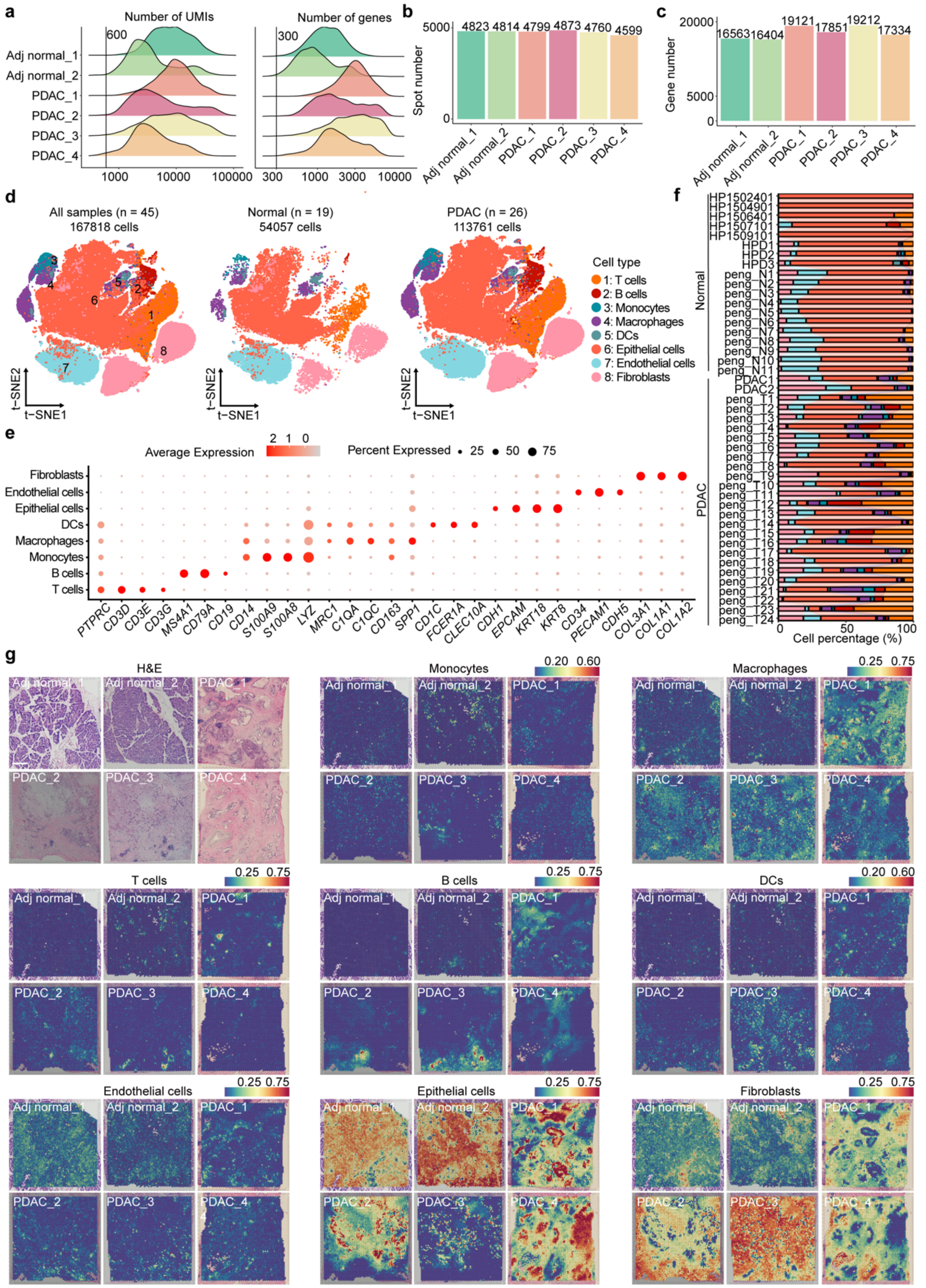


**Supplementary Fig. S2 Spatial transcriptomics and single-cell analysis of adjacent normal and PDAC tissues from human patients.**

**a-c,** Quality control results of spatial transcriptomics data from human PDAC samples.

**a,** The density plots showing the distribution of the number of unique molecular identifiers (UMIs) and the number of genes detected per spot for each sample. The black lines represent the respective cutoff values used for QC.

**b,** The bar graph depicting the number of spots retained after QC for each sample.

**c,** The bar graph displaying the number of genes detected after QC for each sample.

**d,** The t-distributed stochastic neighbor embedding (t-SNE) plots illustrating the distribution of major cell partitions in three categories: all samples (left), normal samples (middle), and PDAC samples (right). The left plot represents all samples (n = 45), containing a total of 167,818 cells; the middle plot represents normal samples (n = 19), containing a total of 54,057 cells; and the right plot represents PDAC samples (n = 26), containing a total of 113,761 cells, as previously described[28]. The colors of the dots represent different cell types: T cells (orange), B cells (red), Monocytes (blue-green), Macrophages (purple), Dendritic cells (pink), Epithelial cells (light blue), Endothelial cells (cyan), and Fibroblasts (peach).

**e,** Dot plot showing the expression levels of specific markers in each cell type.

**f,** Relative fraction of each cell type in human normal pancreas and PDAC tissues across all 45 samples, as shown in Supplementary Fig. S2d.

**g,** H&E staining, the spatial distribution of monocytes, macrophages, T cells, B cells, DCs, Endothelial cells, Epithelial cells and Fibroblasts in spatial transcriptomics data of human adjacent normal and PDAC tissue sections. Scale bars, 1 mm.

### Supplementary Fig. S3

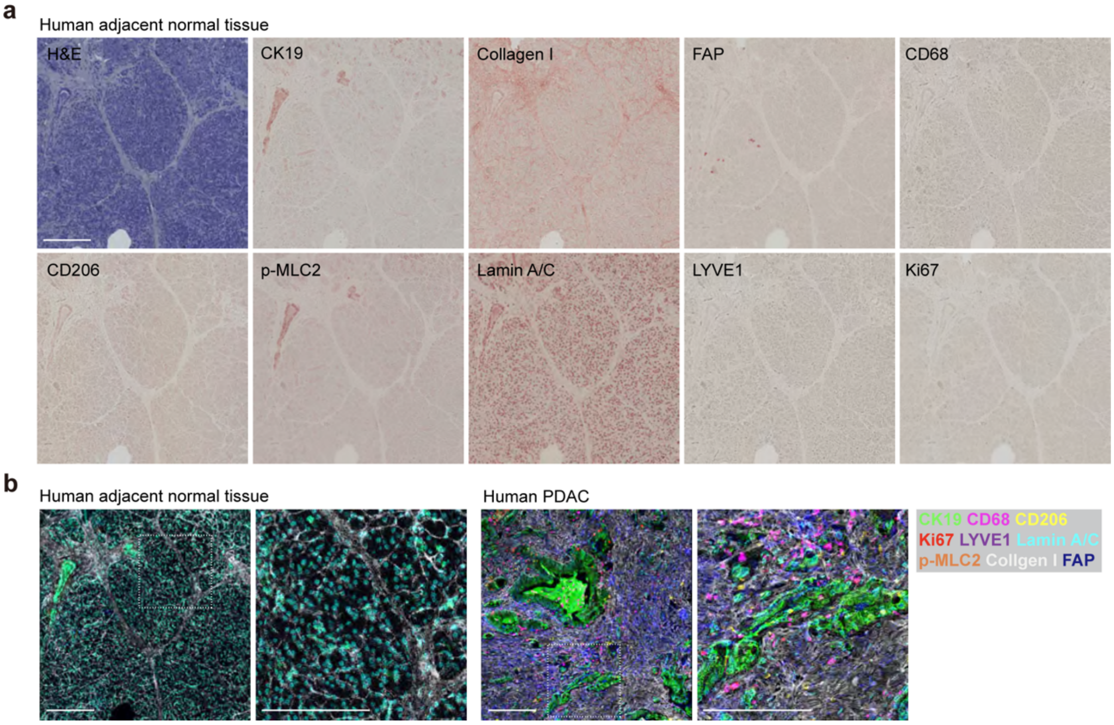


**Supplementary Fig. S3 mIHC staining of human adjacent normal and PDAC tissues**.

**a,** Representative images of human adjacent normal tissue staining with hematoxylin, CK19, Collagen I, FAP, CD68, CD206, LYVE1, Ki67, p-MLC2 and Lamin A/C. Scale bars, 200 μm.

**b,** Overlaid images of markers assigned with pseudocolors in one representative adjacent normal and one representative PDAC area. Scale bars, 200 μm.

### Supplementary Fig. S4

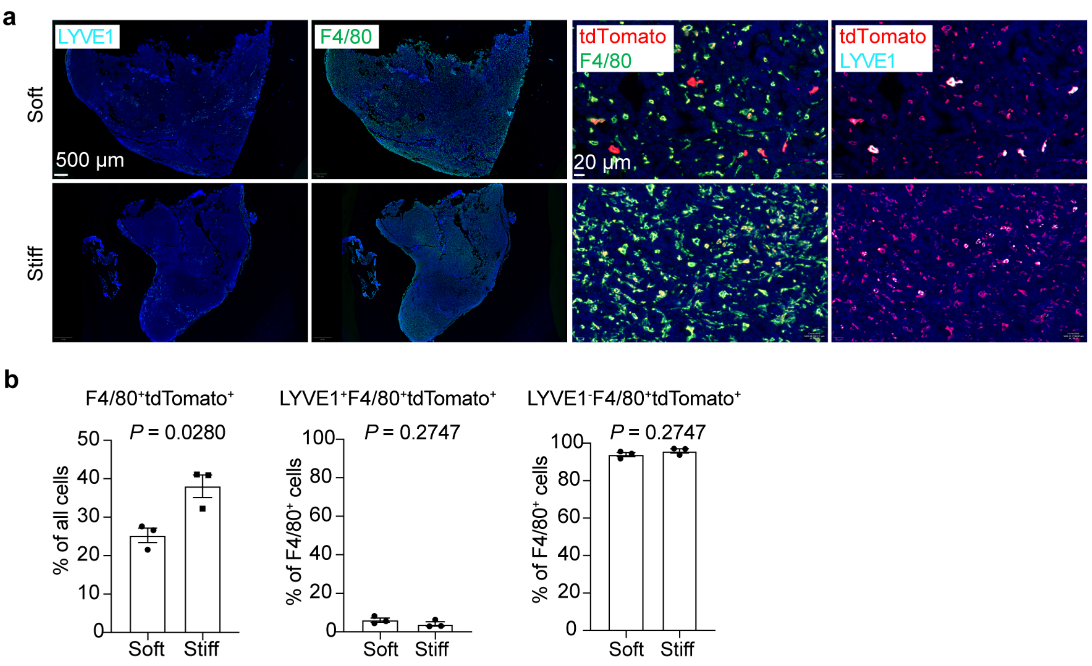


**Supplementary Fig. S4 Quantification of LYVE1$^{+}$ macrophages in mouse PDAC models.**
**a-b,** Stiff or soft hydrogel was minced, combined with KPC cells, and orthotopically injected into the pancreas of G/R$^{fl/+}$, Lyz2-Cre mice. Representative mIHC images (a) and quantification (b) of LYVE1 and F4/80 in orthotopic PDAC tumors from G/R$^{fl/+}$, Lyz2-Cre mice. n = 3 mice per group. *P* values were determined by unpaired two-tailed Student's t-test.

**Supplementary Fig. S5**

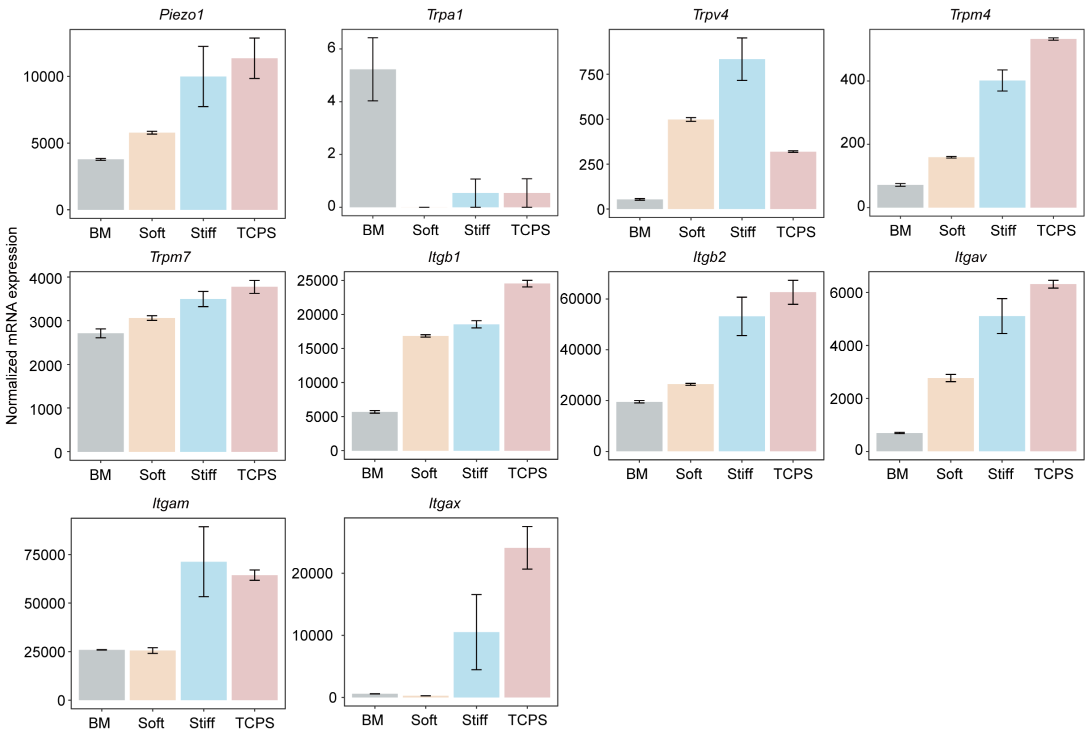


**Supplementary Fig. S5 Stiffness-dependent upregulation of mechanosensitive ion channels and integrins in monocyte differentiation**.

Normalized mRNA expression levels of *Piezo1*, *Trpa1*, *Trpv4*, *Trpm4*, *Trpm7*, *Itgb1*, *Itgb2*, *Itgav*, *Itgam* and *Itgax* in mouse BM progenitors, monocytes differentiated on soft, stiff, and TCPS surfaces for 6 days, assessed by bulk RNA-seq. Data represents two independent experiments.

**Supplementary Fig. S6**

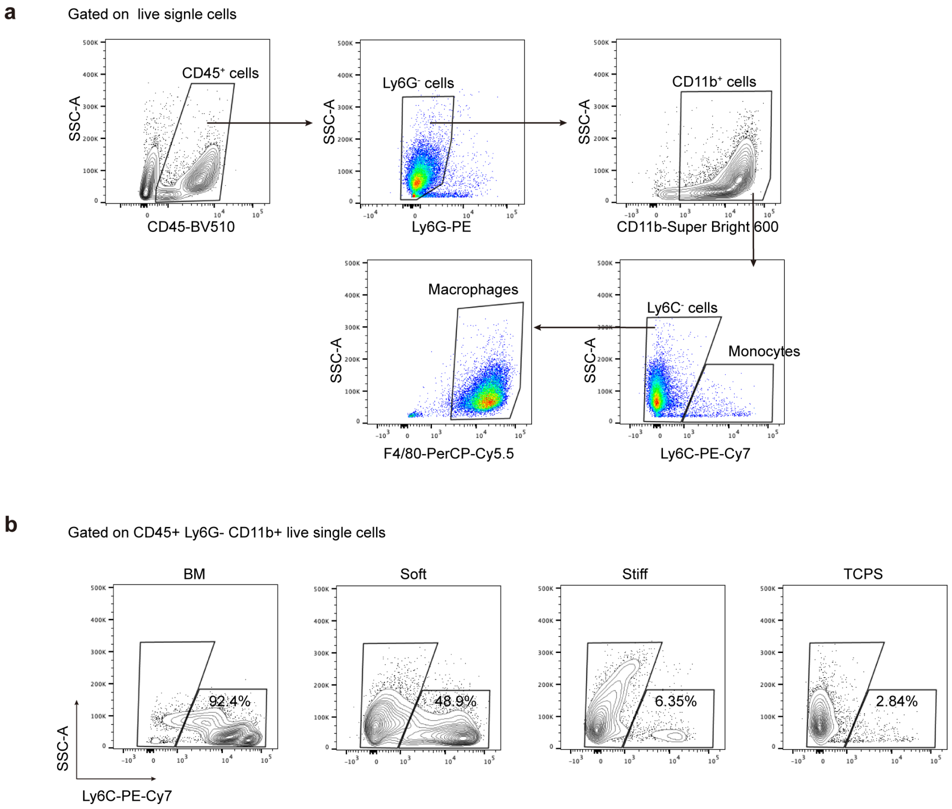


**Supplementary Fig. S6 Flow cytometry analysis of mouse bone marrow derived-monocytes and macrophages.**

**a,** Illustration of the gating strategy employed to identify mouse BM derived-monocytes and macrophages.

**b,** Flow cytometry data showing Ly6C- cells and monocytes in freshly isolated mouse BM cells, and the BM cells stimulated with M-CSF and cultured on soft, stiff, and TCPS surfaces for 6 days.

**Supplementary Fig. S7**

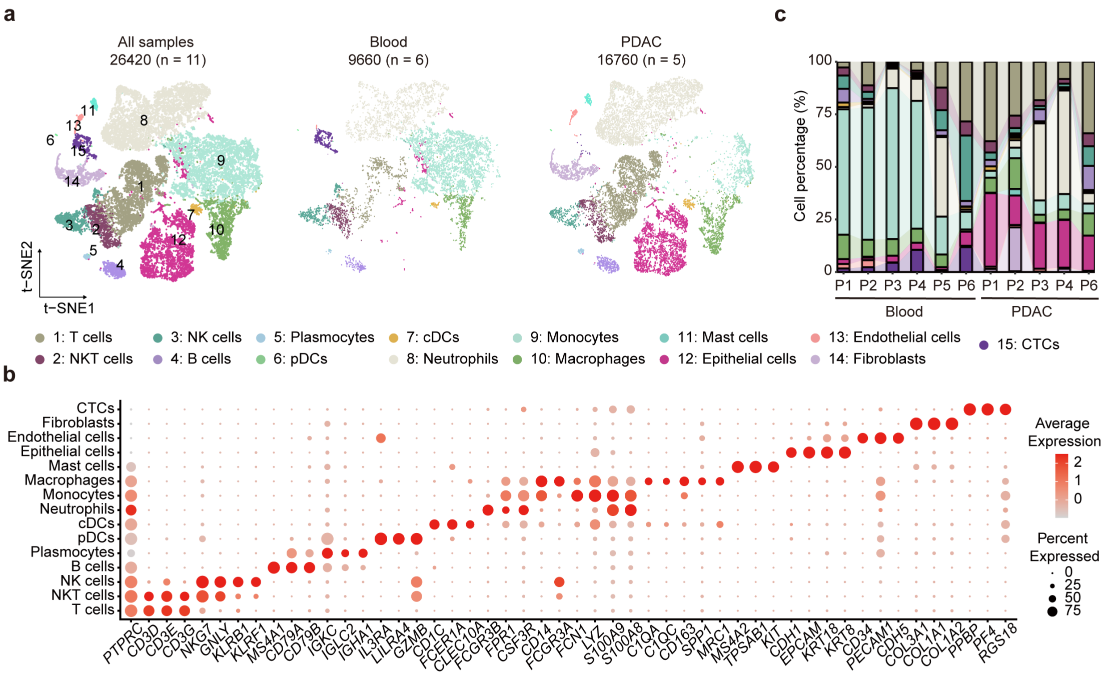

**Supplementary Fig. S7 Quality control results of scRNA-seq data from human blood and PDAC samples.**

**a,** The uniform manifold approximation and projection (UMAP) plots illustrating single-cell clustering of cells from blood and PDAC samples, as described[32]. The left panel represents data from all samples, consisting of 26,420 cells from 11 samples. The middle panel illustrates data from 6 blood samples, containing 9,660 cells. The right panel displays data from 5 PDAC samples, including 16760 cells. The identified cell types include T cells (1), NKT cells (2), NK cells (3), B cells (4), Plasma cells (5), pDCs (6), cDCs (7), Neutrophils (8), Monocytes (9), Macrophages (10), Mast cells (11), Epithelial cells (12), Endothelial cells (13), Fibroblasts (14), and CTCs (15).

**b,** Dot plot showing the expression of representative signature genes across the cell types presented in Supplementary Fig. S7a. The color and size indicate the scaled mean expression and the percentage of cells expressing each gene, respectively.

**c,** Stacked bar chart showing the variation in the proportion of 15 cell types across 11 samples. The samples were categorized into Blood (P1-P6) and PDAC (P1-P4, P6). Each color represents a different cell type, and the height of each color segment indicates the percentage of that cell type within each sample.

**Supplementary Fig. S8**

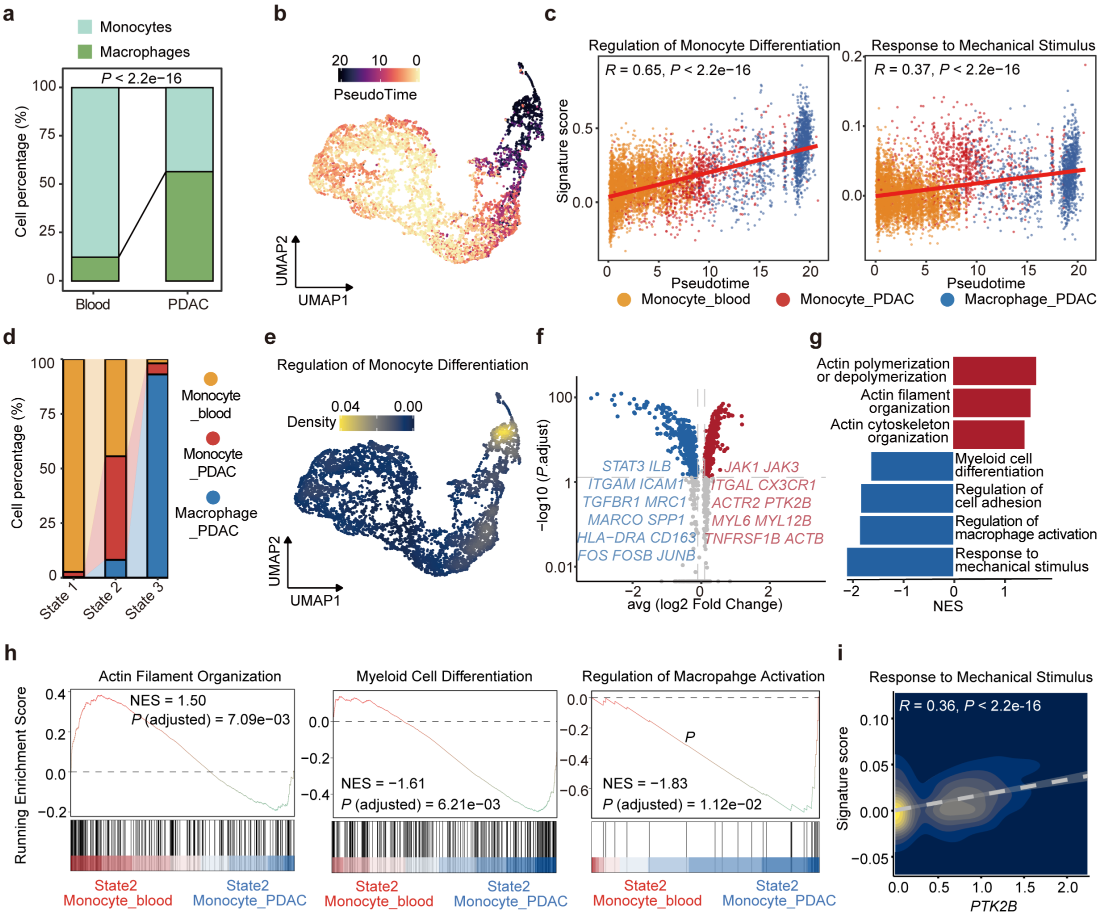


**Supplementary Fig. S8 Spatiotemporal analysis of monocytes and macrophages in human PDAC patients.**

**a,** Bar chart comparing the relative proportions of monocytes and macrophages in Blood and PDAC samples. The proportions were analyzed using a Chi-squared test, with a *P* value of less than 2.2e-16. Monocytes are represented in light green and macrophages in dark green.

**b,** UMAP visualization showing the results of Monocle2 pseudotime analysis, with cells colored according to their pseudotime values.

**c,** Plots showing the dynamic expression of two pathways, Regulation of Monocyte Differentiation (left) and Response to Mechanical Stimulus (right), along the pseudotime axis. The red lines indicate the dynamic expression trends of the pathways, determined using a linear regression model. The Regulation of Monocyte Differentiation pathway had an *R* of 0.65, and the Response to Mechanical Stimulus pathway had an R of 0.38, both with $P < 2.2e\text{-}16$. The colors of the dots correspond to the cell types: Monocyte blood (orange), Monocyte PDAC (red), and Macrophage PDAC (blue).

**d,** Stacked bar chart comparing the relative proportions of three cell states (State 1 to State 3) among three cell types: Monocyte blood, Monocyte PDAC, and Macrophage PDAC. The colors represent the cell types: Monocyte blood (orange), Monocyte PDAC (red), and Macrophage PDAC (blue).

**e,** UMAP plot depicting the density distribution of cells based on their activity of the Regulation of Monocyte Differentiation pathway. Higher densities of active cells are shown in yellow, while lower densities are indicated in blue.

**f,** Volcano plot showing the differential gene expression in State 2 when comparing monocytes from blood with monocytes from PDAC. These differential genes were identified using the Wilcox test with criteria of *P*.adjust < 0.05, |log2 fold change| > 0.1, and FDR correction. Genes upregulated in PDAC monocytes are highlighted in red, while those upregulated in blood monocytes are highlighted in blue. The x-axis represents the average log2 fold change, and the y-axis represents the -log10 adjusted *P* value.

**g,** Bar chart showing the results of Gene Set Enrichment Analysis (GSEA) using differential genes from the comparison of monocytes from blood with monocytes from PDAC in State 2. Enriched pathways in PDAC monocytes are highlighted in red, while enriched pathways in blood monocytes are highlighted in blue. The x-axis represents the normalized enrichment score (NES).

**h,** The GSEA results of myeloid cell differentiation (left), actin filament organization (mid) and actin cytoskeleton organization (right).

**i,** Pearson correlation analysis of *PTK2B* and the signature score of response to mechanical stimulus in PDAC monocytes in State 2. The white dashed line represents the linear regression result, with $R = 0.36$ and $P < 2.2e{-16}$. The color in the plot indicates the density of cell distribution.

### Supplementary Fig. S9

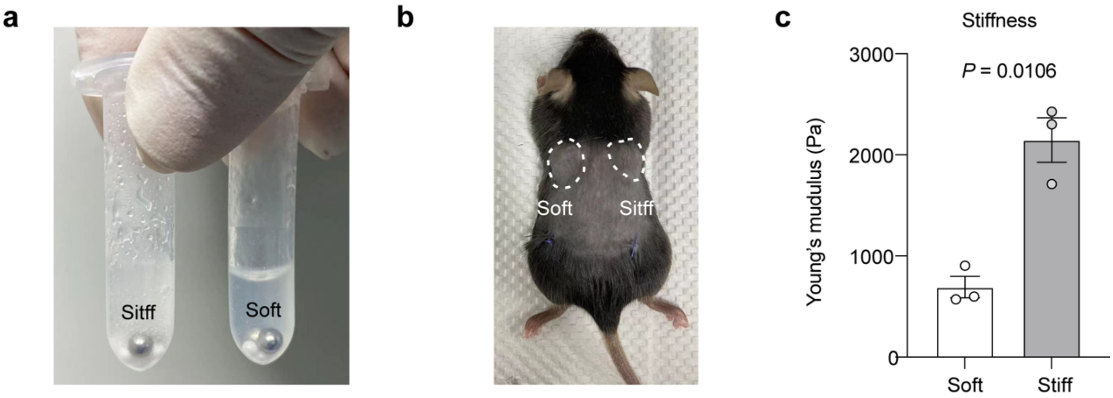


**Supplementary Fig. S9 Subcutaneously simulation of PDAC models.**

**a,** Representative image of stiff or soft hydrogels after grinding.

**b,** Representative image of mice subcutaneously implanted with a mixture of stiff or soft hydrogels and KPC cells. The white dashed lines indicate the implanted hydrogel and KPC cells.

**c,** The Young's modulus of simulated PDAC tissues measured 3 weeks after implantation. n = 3 mice per group. *P* value was determined by a two-tailed unpaired Student's t-test.

### Supplementary Fig. S10

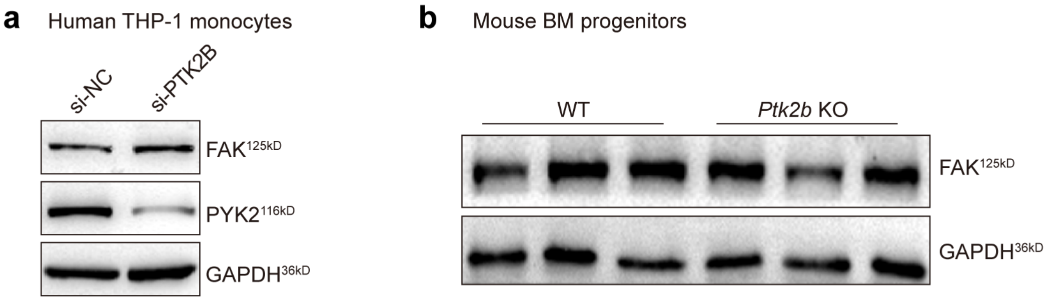


**Supplementary Fig. S10 Western blot analysis of FAK in *PTK2B* silenced THP-1 cells and *Ptk2b* knockout BMDMs.**

**a,** Western blot analysis of FAK in *PTK2B* silenced THP-1 cells. THP-1 cells were transfected with si-NC or si-PTK2B, and then stimulated with PMA for 48 hours. The protein expression of FAK and PYK2 were analyzed by western blot.

**b,** Western blot analysis of FAK in *Ptk2b* knockout BMDMs. Mouse BM cells were isolated from *Ptk2b*$^{fl/fl}$ (WT) and *Ptk2b*$^{fl/fl}$*, Lyz2-Cre* (*Ptk2b* KO) mice, and subsequently stimulated with M-CSF for 6 days. The protein expression of FAK in BMDMs was analyzed by western blot.

### Supplementary Fig. S11

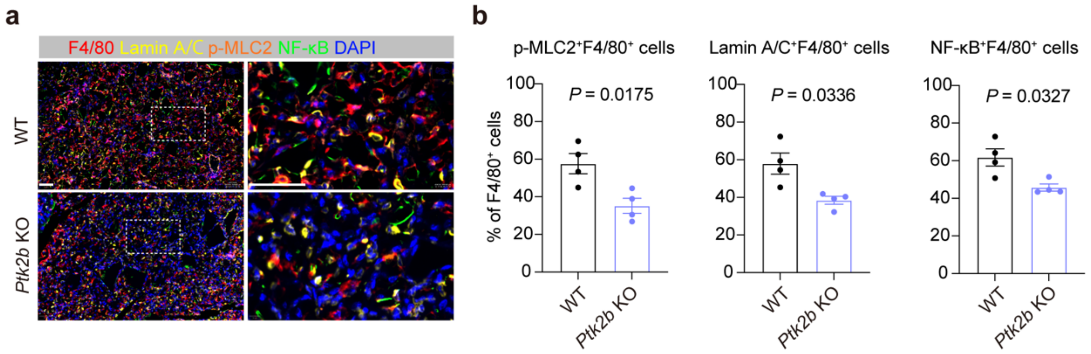


**Supplementary Fig. S11 PYK2 disrupts mechanosignaling during monocyte-to-macrophage differentiation in stiff TME.**

**a-b,** Representative images (a) and quantification (b) of mIHC staining of F4/80, Lamin A/C, p-MLC2, NF-κB p65 and DAPI of tissues collected 3 weeks after orthotopically pancreatic implantation in *Ptk2b*$^{fl/fl}$ and *Ptk2b*$^{fl/fl}$, Lyz2-Cre mice. n = 4 mice in each group. Scale bars, 50 μm.

**Supplementary Fig. S12**

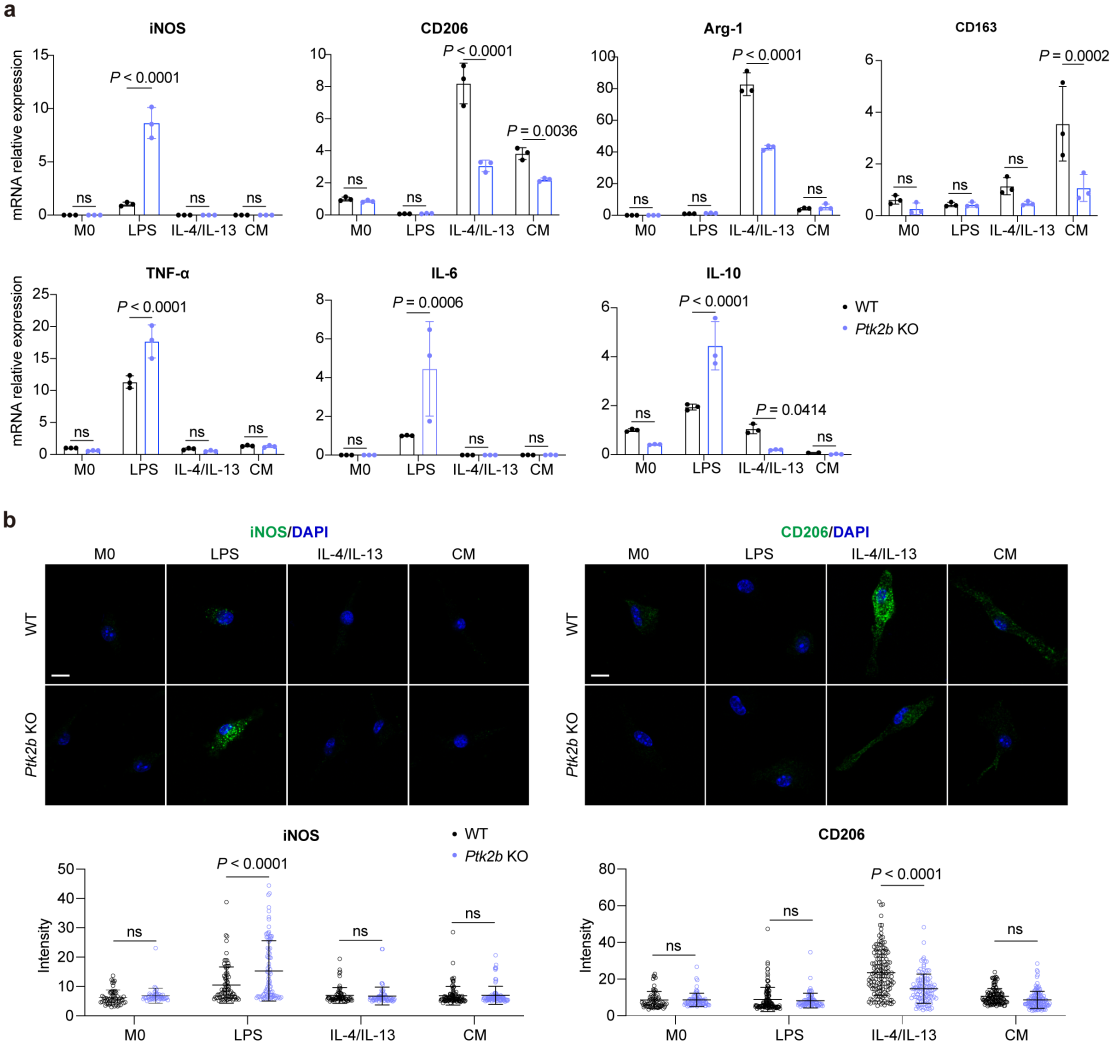


**Supplementary Fig. S12 Polarization of WT and *Ptk2b* KO BMDMs.**

**a,** mRNA relative expression of iNOS, CD206, Arg-1, CD163, TNF-α, IL-6 and IL-10 in WT and *Ptk2b* KO BMDMs treated with LPS, IL-4/IL-13 and condition medium (CM), respectively.

**b,** Representative immunofluorescent images and quantification of iNOS and CD206 in WT and *Ptk2b* KO BMDMs treated with LPS, IL-4/IL-13 and CM, respectively.

*P* values were determined by ordinary two-way ANOVA. Data shown are representative of three experiments.

**Supplementary Fig. S13**

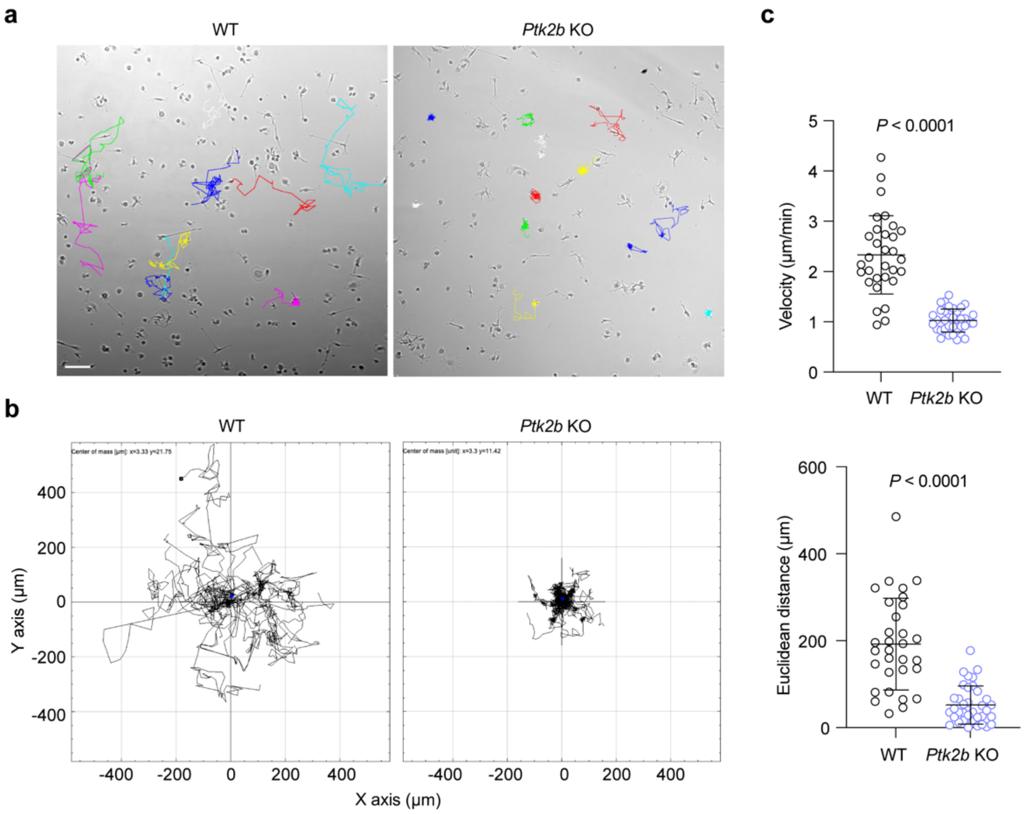


**Supplementary Fig. S13 PYK2 impacts migration of BMDMs.**

**a,** Representative phase-contrast images of WT and *Ptk2b* KO BMDMs captured by time-lapse microscopy at 10-minute intervals. Cell trajectory tracks for WT and *Ptk2b* KO BMDMs were manually traced using the Manual Tracking plugin of ImageJ and marked in color. Scale bars, 100 µm.

**b,** Trajectory plots the trajectories of WT and *Ptk2b* KO BMDMs. All tracks were set to a common origin using Chemotaxis plugin of ImageJ.

**c,** Quantitative analysis of Euclidean distance (µm) and velocity (µm/min) of WT and *Ptk2b* KO BMDMs. Analysis was conducted on WT (31 cells) and *Ptk2b* KO (36 cells) BMDMs from three independent experiments. *P* values were determined by unpaired two-tailed Student's t-test.

### Supplementary Fig. S14

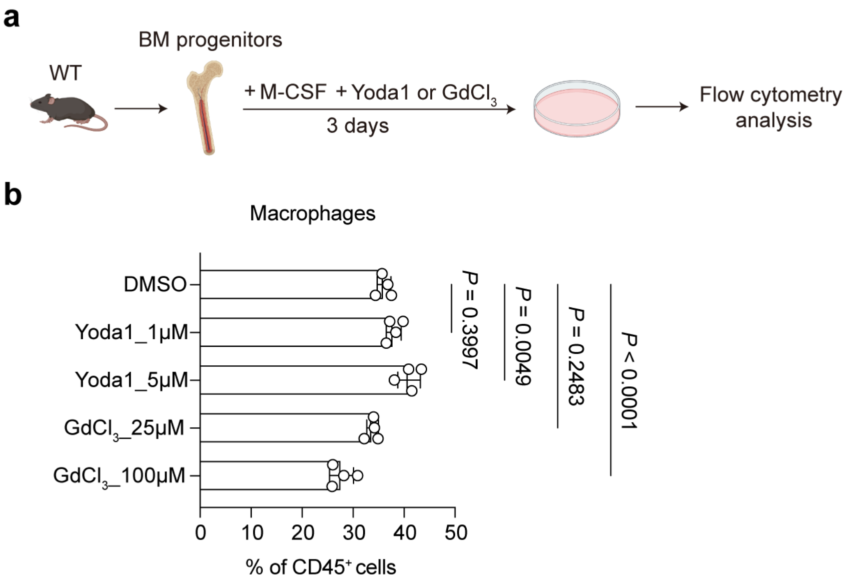


**Supplementary Fig. S14 Role of PIEZO1 in monocyte differentiation.**

**a,** Schematic representation of the differentiation of mouse BM progenitors into macrophages using M-CSF, with the application of PIEZO1 agonist (Yoda1) or inhibitor ($GdCl_3$).

**b,** Flow cytometry analysis of mouse progenitors treated with M-CSF and Yoda 1 (1 and 5 μM) or $GdCl_3$ (25 and 50 μM) for 3 days. Data represents four independent experiments. *P* values were calculated using one-way ANOVA.

**Supplementary Fig. S15**

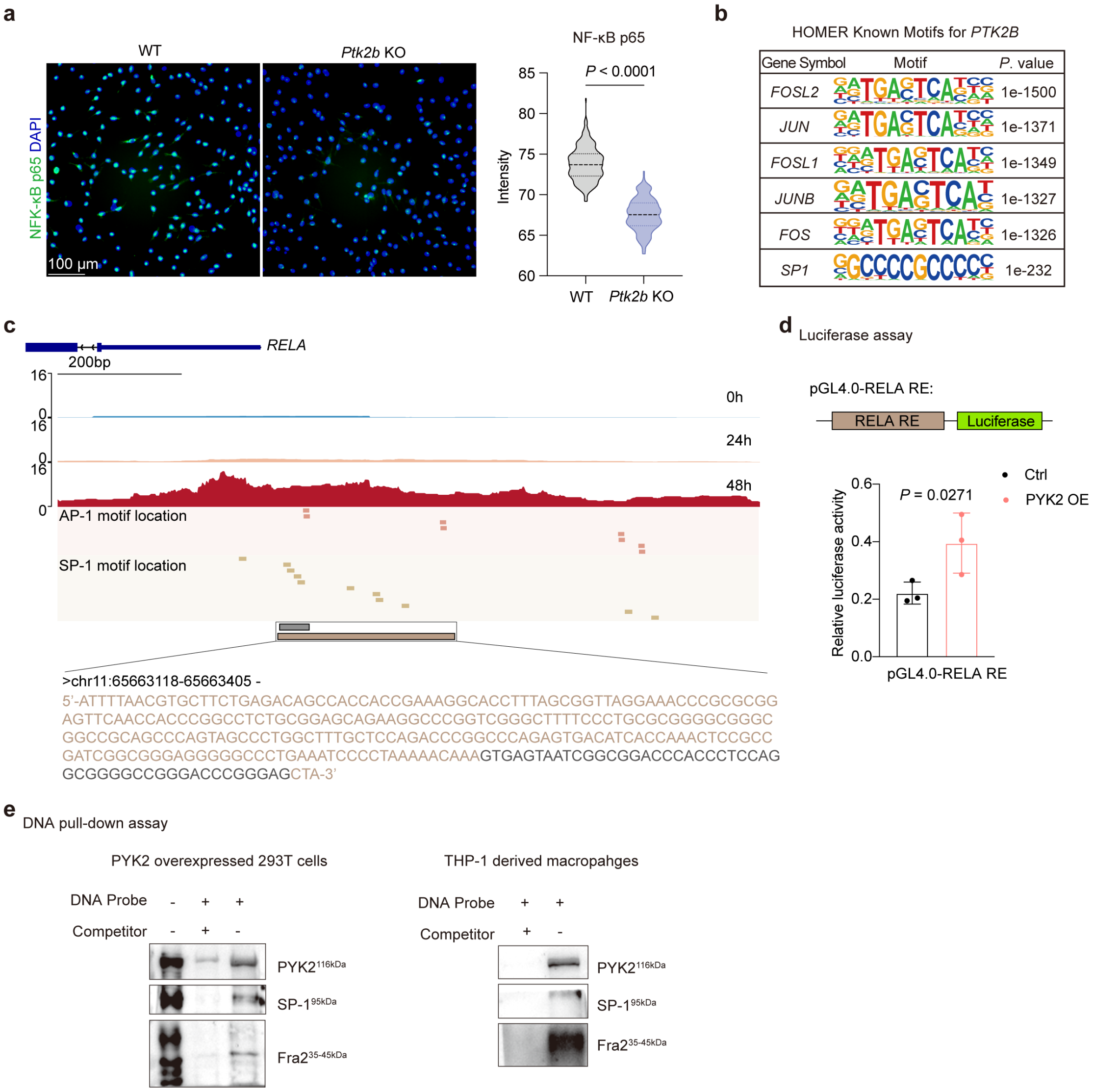


**Supplementary Fig. S15 PYK2 as a transcriptional co-regulator with AP-1 or/and SP-1.**

**a,** Representative immunofluorescent images (left) and quantification (right) of NF-κB p65 expression in WT and *Ptk2b* KO BMDMs. n = 234 (WT) and 180 (*Ptk2b* KO) cells from three independent experiments. Scale bars, 100 μm.

**b,** The most significantly enriched motifs in PYK2-binding regions in THP-1 derived macrophages, identified by HOMER motif analysis.

**c,** CUT&Tag profiles of the RELA promoter locus in suspended THP-1 cells and those primed with PMA for 24 h or 48 h, spanning ~1 kb upstream of the transcription start site. Middle tracks highlight predicted AP-1 (red) and SP-1 (yellow) binding sites. Bottom tracks denote regions used for luciferase assay (brown) and DNA pull-down assay (gray), with corresponding sequences.

**d,** Luciferase assay using a pGL4.0 plasmid containing a DNA oligonucleotide with SP-1 and AP-1 binding motifs from the RELA promoter (RELA response element, RE), and pRL Renilla luciferase control plasmid transfected into 293T cells or the PYK2-overexpressing 293T cells. Data represent three independent replicates.

**e,** DNA pull-down assay to detect PYK2 binding to AP-1 and SP-1 motifs. A 48 bp biotinylated DNA probe was conjugated to magnetic beads. A non-biotinylated DNA with an identical sequence was used as a competitor. Nuclear extracts from PMA-treated (48 h) THP-1 cells or PYK2-overexpressing 293T cells were incubated with the beads. Bound proteins were resolved by SDS-PAGE and analyzed by Western blot with antibodies against PYK2, Fra2, and SP-1.

**Supplementary Fig. S16**

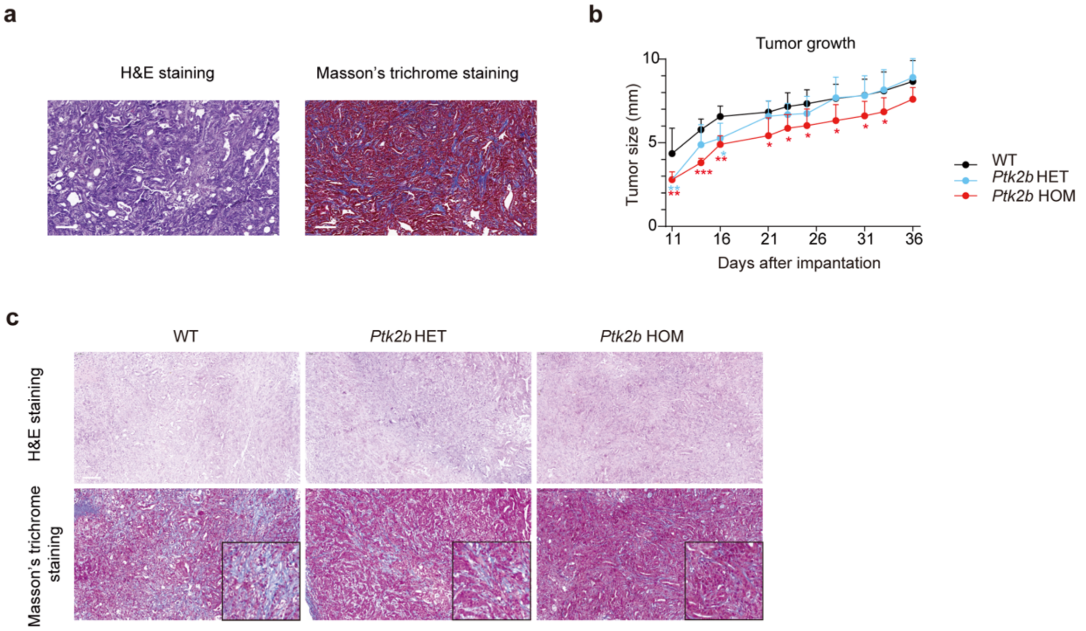


**Supplementary Fig. S16 Loss of PYK2 in myeloid precursors alters TME.**

**a,** Representative images of H&E and Masson's trichrome staining of subcutaneous tumor used for subsequent orthotopic implantation. KPC cells were subcutaneously injected into the WT mice, and the tumor tissue was collected two weeks after the injection. One portion of the tissue was formalin-fixed, paraffin-embedded, and sectioned for H&E and Masson's trichrome staining. The remaining portion was divided into fragments and orthotopically implanted into the pancreas of recipient mice. Scale bars, 100 μm.

**b,** Tumor growth curves of PDAC in WT (n = 6), *Ptk2b* HET (n = 6), and *Ptk2b* HOM (n = 7) mice. *P* values were determined by ordinary two-way ANOVA. *$P$<0.05, **$P$<0.01, ***$P$<0.001.

**c,** Representative images of H&E staining and Masson's trichrome staining of PDAC in WT, *Ptk2b* HET, and *Ptk2b* HOM mice. Scale bars, 100 μm.

### Supplementary Fig. S17

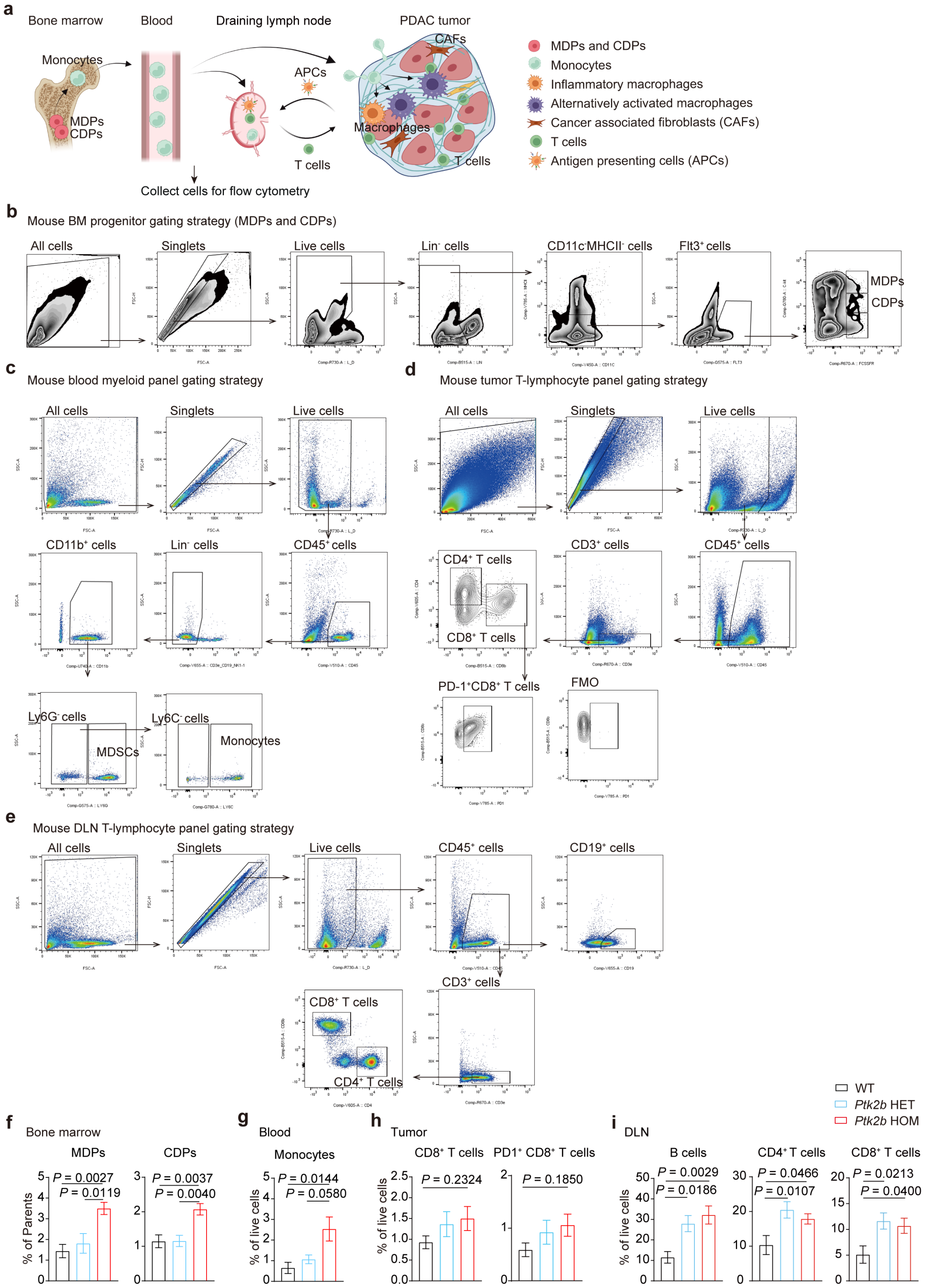


**Supplementary Fig. S17 Flow cytometry analysis of mouse tissues.**

**a,** Schematic diagram illustrating the development and distribution of monocytes and macrophages during PDAC progression. Monocytes are continually produced in the bone marrow from monocyte-dendritic cell progenitors (MDPs). These monocytes enter circulation, eventually infiltrating the PDAC tumor tissue, where they undergo differentiation into macrophages. Within the TME, these macrophages engage with cancer-associated fibroblasts (CAFs) and T cells, influencing and contributing to the progression of PDAC.

**b,** Gating strategy for MDPs and CDPs in mouse BM cells.

**c,** Gating strategy for myeloid cells in mouse blood.

**d,** Gating strategy for T cells in mouse PDAC tissue.

**e,** Gating strategy for B cells and T cells in mouse DLN.

**f-i,** Quantification of MDPs and CDPs from bone marrow, monocytes from blood and B cells and T cells in DLN, $CD8^+$ T cells and $PD1^+CD8^+$ T cells in tumors by flow cytometry. n = 6 (WT), n = 6 (*Ptk2b* HET), n = 7 (*Ptk2b* HOM) mice. *P* values were determined by ordinary one-way ANOVA.

**Supplementary Fig. S18**

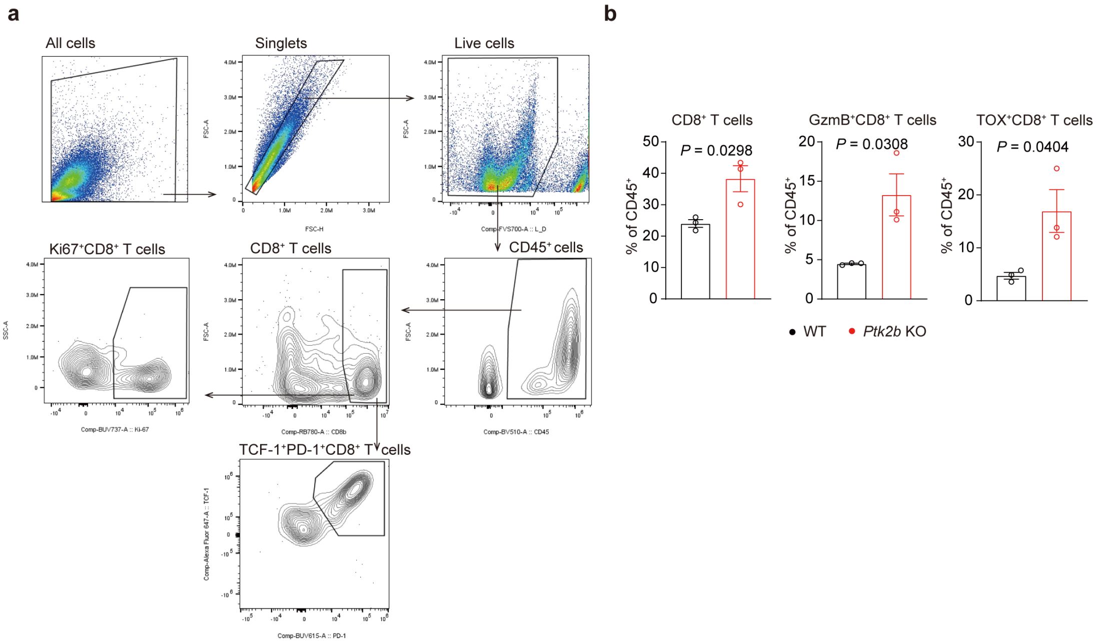


**Supplementary Fig. S18 Spectral cytometry analysis of T cells in co-culture system.**

**a,** Gating strategy of T cells in the co-culture system.

**b,** Quantificatioin of CD8$^+$ T cells, GzmB$^+$CD8$^+$ T cells, and TOX$^+$ CD8$^+$ T cell in co-culture system. *P* values were determined by two-tailed unpaired Student's t-test. Data shown are representative of three experiments.

**Supplementary Fig. S19**

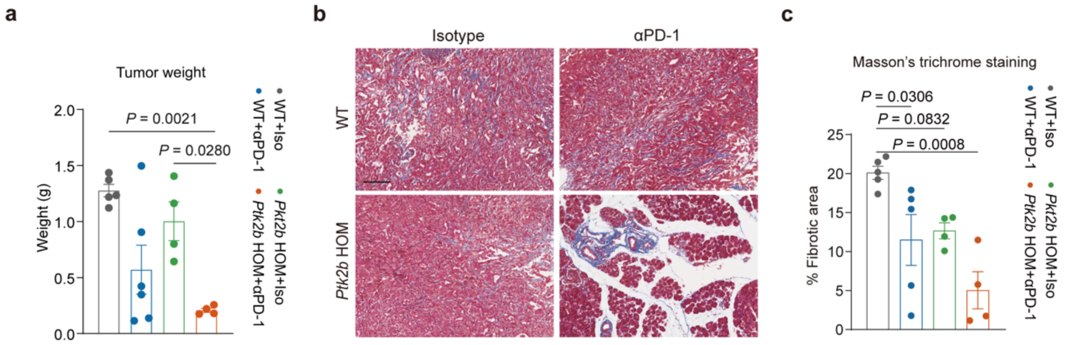


**Supplementary Fig. S19 Loss of PYK2 in BM progenitors alters TME and enhances the efficacy of immunotherapy.**

**a,** Quantification of tumor weight in WT and *Ptk2b* KO mice after 4 rounds of Isotype or αPD-1 treatment. n = 5 (WT+Iso), n = 6 (WT+αPD-1), n = 4 (*Ptk2b* HOM+Iso), n = 4 (*Ptk2b* HOM+αPD-1) mice.

**b,** Representative images of Masson's trichrome staining of PDAC in WT and *Ptk2b* KO mice after 4 rounds of Isotype or αPD-1 treatment. Scale bars, 100 µm.

**c,** Quantification of fibrotic area of PDAC in WT and *Ptk2b* KO mice after 4 rounds of Isotype or αPD-1 treatment. n = 5 (WT+Iso), n = 5 (WT+αPD-1), n = 4 (*Ptk2b* HOM+αPD-1), n = 4 (*Ptk2b* HOM+αPD-1) mice.

*P* values in **(a)** and **(c)** were determined by ordinary one-way ANOVA.

**Supplementary Fig. S20**

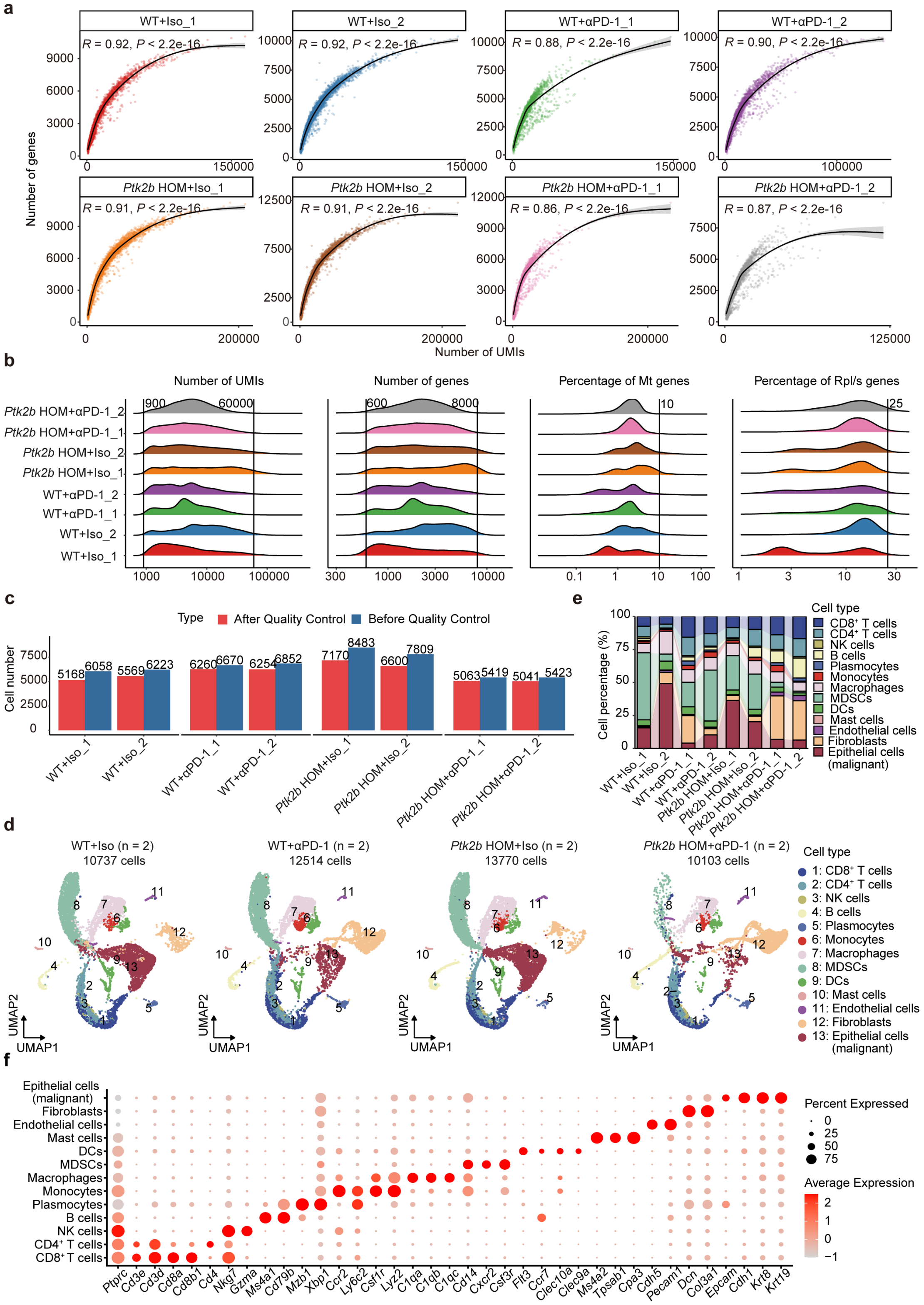


**Supplementary Fig. S20 Quality control scRNA-seq data of WT+Iso, WT+αPD-1, *Ptk2b* KO+Iso and *Ptk2b* KO+αPD-1 groups.**

**a,** Scatter plots showed Pearson's correlation between the numbers of genes and numbers of UMIs across WT+Iso (n = 2), WT+αPD-1 (n = 2), *Ptk2b* KO+Iso (n = 2), and *Ptk2b* KO+αPD-1 (n = 2) samples.

**b,** The distribution of UMIs, the numbers of genes, the percentage of Mitochondrial (Mt) genes, and the percentage of Ribosomal proteins large/small (Rpl/s) genes in single cells. The black line represents the threshold for each feature.

**c,** The bar chart displayed the number of cells in each sample before and after quality control, with blue representing the original cell count and red indicating the cell count after filtration.

**d,** UMAP of cell populations in WT+Iso (10,737 cells), WT+αPD-1 (12,514 cells), *Ptk2b* KO+Iso (13,770 cells), *Ptk2b* KO+αPD-1 (10,103 cells) group. A total of 13 different cell types were identified.

**e,** The bar plot representing the cell percentage for each cell type in each sample.

**f,** Dot plot showing the expression of canonical marker genes corresponding to each cell type from Supplementary Fig. S20d. The color indicates the normalized expression, and the dot size represents the percentage of expressed cells.

### Supplementary Fig. S21

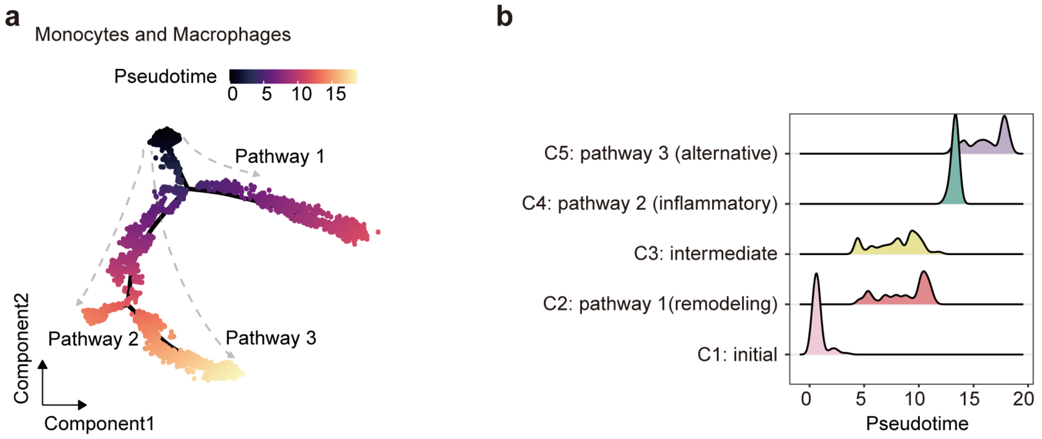


**Supplementary Fig. S21 Transcriptional landscape of monocytes and macrophages from WT+Iso, WT+αPD-1, *Ptk2b* KO+Iso and *Ptk2b* KO+αPD-1 groups.**

**a,** UMAP visualization showing the results of Monocle2 pseudotime analysis for all Mono/Macro cells from WT+Iso, WT+αPD-1, *Ptk2b* KO+Iso, and *Ptk2b* KO+αPD-1 groups. The cells are colored according to their pseudotime values. The dashed lines represent three inferred endpoints**.**

**b,** The ridge plots showed the temporal distribution of each cell type from Fig. 6c along the pseudotime axis.

**Supplementary Fig. S22**

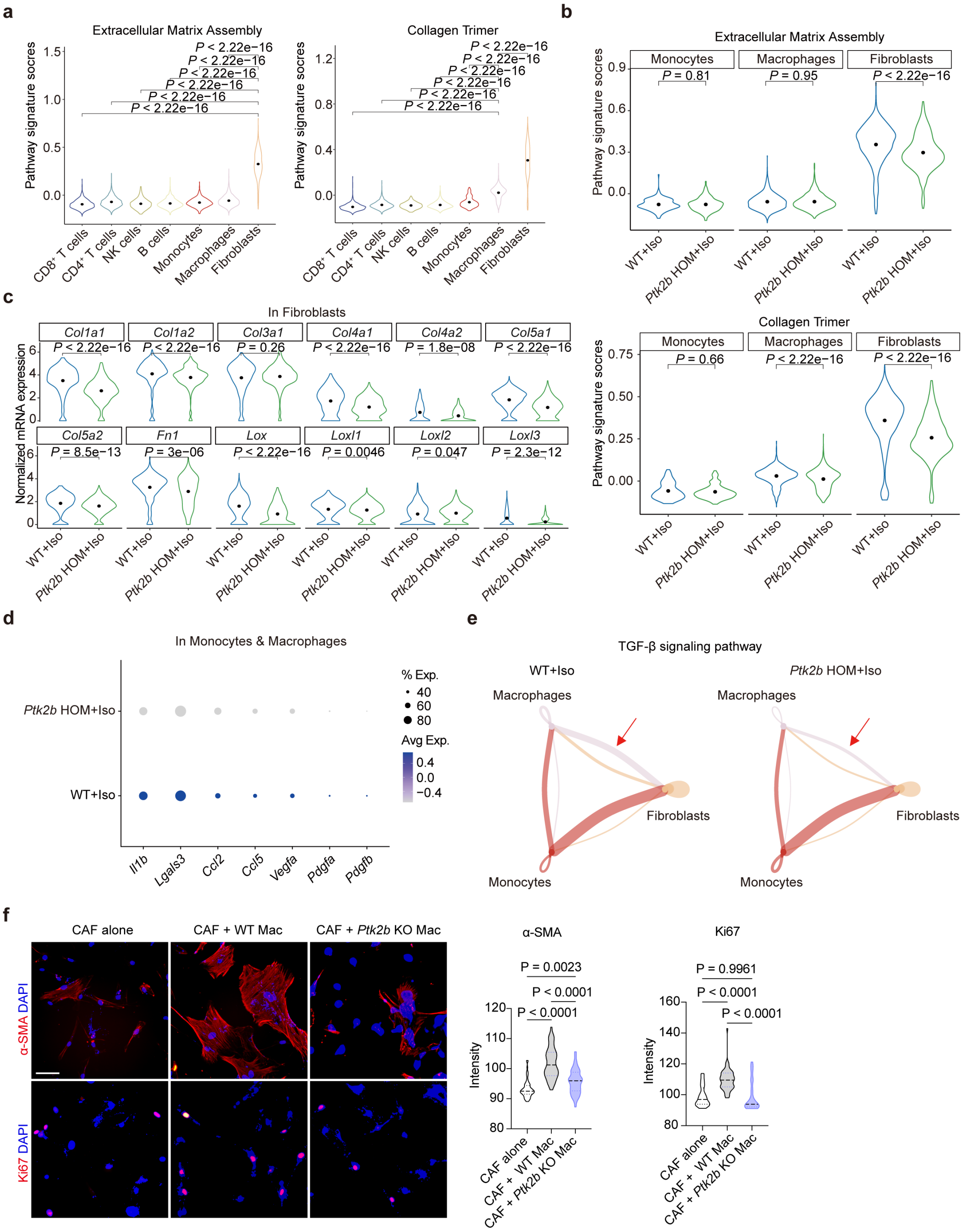


**Supplementary Fig. S22 Effects of *Ptk2b* deficiency in BM progenitors on fibrosis in PDAC.**

**a,** Signature scores for "Extracellular Matrix Assembly" pathway and "Collagen Trimer" pathway across immune cells and fibroblasts in mouse PDAC samples.

**b,** Comparison of signature scores for monocytes, macrophages, and fibroblast between the WT + Iso and *Ptk2b* HOM + Iso groups.

**c,** Normalized mRNA expression levels of *Col1a1*, *Col1a2*, *Col3a1*, *Col4a1*, *Col4a2*, *Col5a1*, *Col5a2*, *Fn1*, *Lox*, *Loxl1*, *Loxl2*, and *Loxl3* in fibroblasts from the WT + Iso and *Ptk2b* HOM + Iso groups.

**d,** Normalized mRNA expression levels of *Il1b*, *Lgals3*, *Ccl2*, *Ccl5*, *Vegfa*, *Pdgfa*, and *Pdgfb* in monocytes and macrophages from the WT + Iso and *Ptk2b* KO + Iso groups.

**e,** Predicted TGF-β signaling interactions among monocytes, macrophages, and fibroblasts in the WT + Iso and *Ptk2b* KO + Iso groups.

**f,** Representative immunofluorescent images (left) and quantification (right) of Ki67 and α-SMA in CAFs. CAFs were co-cultured with WT macrophages or *Ptk2b* KO macrophages in a Transwell system for 3 days, then fixed and stained for Ki67 and α-SMA. n = 58 (CAF alone), 53 (CAF + WT macrophages), and 68 (CAF + *Ptk2b* KO macrophages) cells for Ki67 quantification. n = 64 (CAF alone), 56 (CAF + WT macrophages), and 61 (CAF + *Ptk2b* KO macrophages) cells for α-SMA quantification. Scale bars, 100 μm. Data represents three independent experiments. *P* values were calculated using one-way ANOVA.

### Supplementary Fig. S23

a

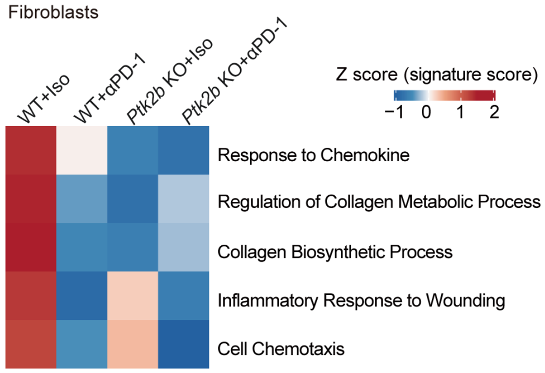


**Supplementary Fig. S23 Signature scores of biological pathways in fibroblasts across experimental groups**

**a,** Heatmap illustrating the signature scores of key biological pathways in fibroblasts from WT and *Ptk2b* KO mice treated with Isotype control or αPD-1. Pathways analyzed include the response to chemokine, regulation of collagen metabolic process, collagen biosynthetic process, inflammatory response to wounding, and cell chemotaxis.

### Supplementary Fig. S24

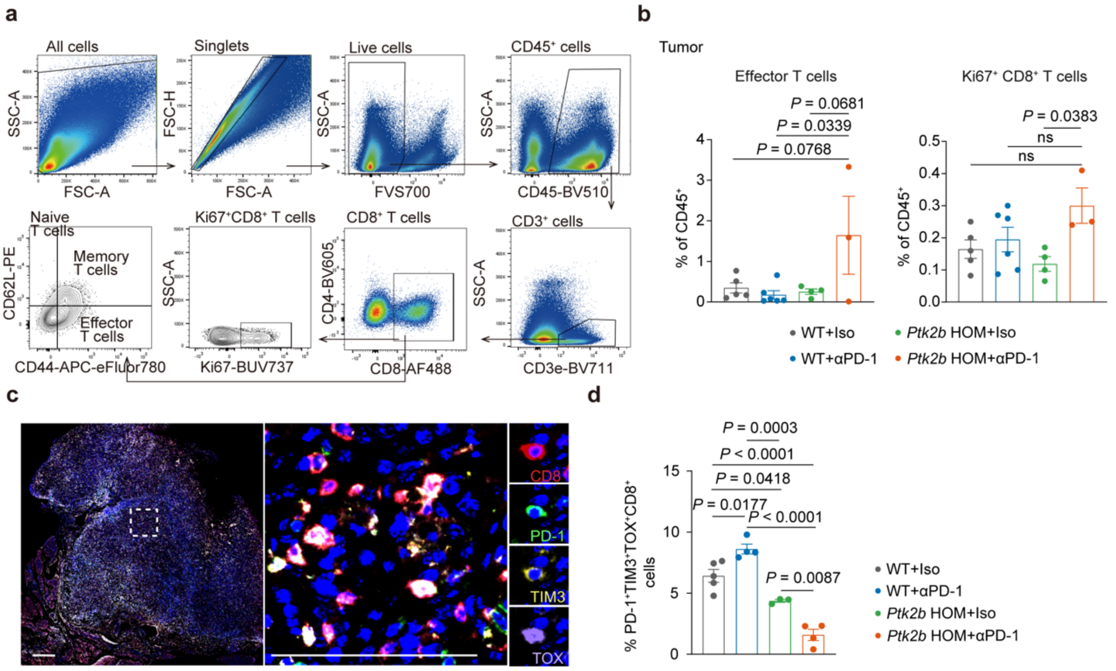


**Supplementary Fig. S24 Flow cytometry and mIHC analysis of $CD8^+$ T cells in WT+Iso, WT+αPD-1, *Ptk2b* KO+Iso and *Ptk2b* KO+αPD-1 groups.**

**a-b,** Flow cytometry gating strategy (a) and quantification (b) of T cells in WT+Iso, WT+αPD-1, *Ptk2b* KO+Iso and *Ptk2b* KO+αPD-1 groups. n = 5 (WT+Iso), n = 6 (WT+αPD-1), n = 4 (*Ptk2b* HOM+Iso), n = 3 (*Ptk2b* HOM+αPD-1) mice.

**c-d,** Representative images (c) and quantification (d) of mIHC staining of CD8, PD-1, TIM-3, TOX, and DAPI in PDAC in WT or *Ptk2b* HOM mice that received either Isotype control or αPD-1 treatment. n = 5 (WT+Iso), n = 4 (WT+αPD-1), n = 3 (*Ptk2b* HOM+ Iso), n = 4 (*Ptk2b* HOM+αPD-1) mice. Scale bars, 500 μm. *P* values were determined by ordinary one-way ANOVA.

**Supplementary Fig. S25**

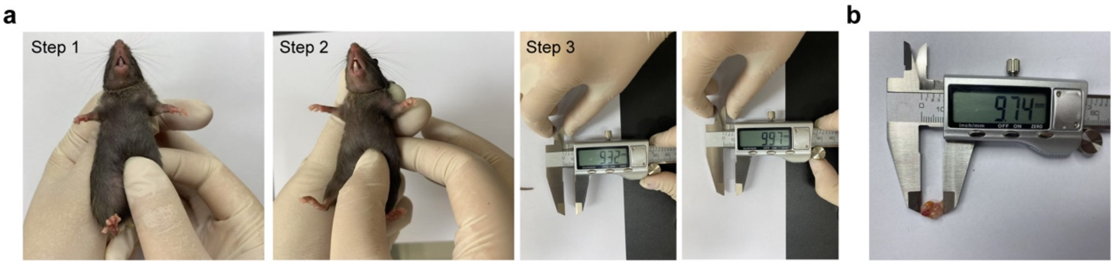


**Supplementary Fig. S25 Palpation-based tumor size measurement.**

**a,** Illustration of the palpation technique for measuring orthotopic pancreatic tumor size. The procedure includes Step 1: Gentle palpation of the implantation site near the spleen to localize the tumor; Step 2: Immobilization of the tumor between the thumb and forefinger; Step 3: Measurement of the tumor diameter between the thumb and forefinger via digital calipers, which is repeated 2–3 times for consistency.

**b,** Longitudinal tumor size measurements obtained at necropsy.

**Supplementary Fig. S26**

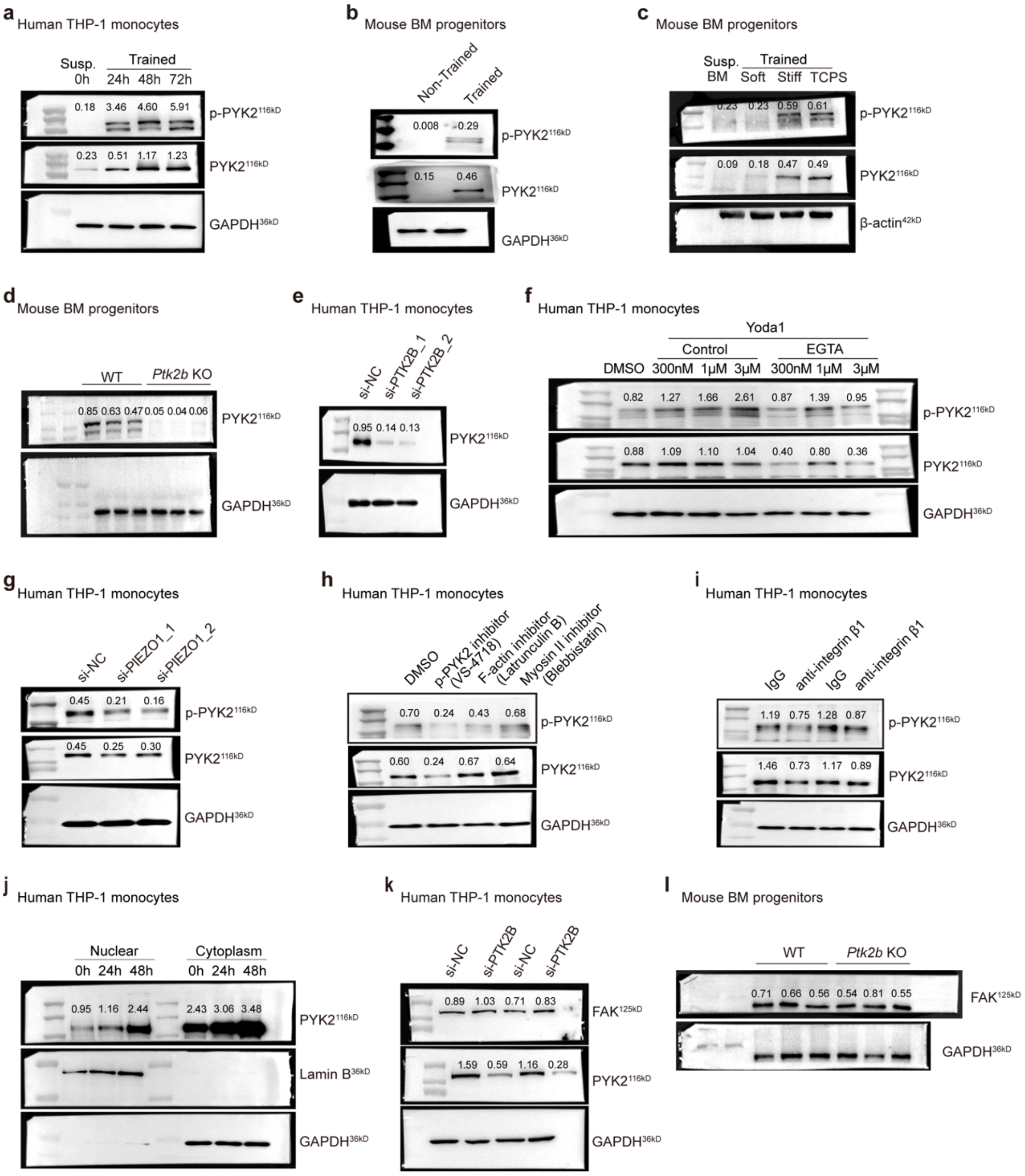


**Supplementary Fig. S26 Quantification of western blot data.**

**a-l,** The full blots and corresponding quantifications from the western blot analyses presented in Fig. 3a, Fig. 3d, Fig. 3e, Fig. 3h, Fig. 3m; Fig. 4a, Fig. 4d-f; Supplementary Fig. S10a and b. Quantification was performed using ImageJ and normalized to housekeeping proteins.
